# Differential tolerance for SEA domain misfolding encodes a MAPK pathway-specific response

**DOI:** 10.64898/2026.05.06.723240

**Authors:** Ankita Priyadarshini, Paul J. Cullen

## Abstract

Signaling pathways often share components yet produce highly specialized biological responses. How signaling specificity is achieved between pathways utilizing common components is a fundamental question. In budding yeast, the same transmembrane mucin, Msb2, regulates two Mitogen-Activated Protein Kinase (MAPK) pathways controlling filamentous growth (fMAPK) and the response to osmotic stress (HOG). How this shared sensor distinguishes between stimuli and regulates different pathways is not clear. Using structure-guided analysis, we identified a conserved SEA (Sea urchin sperm protein, Enterokinase, Agrin) domain in fungal mucins and found that mutations disrupting protein folding selectively impair one pathway (fMAPK) but were tolerated by another (HOG). Mechanistically, these differences revealed distinct modes of signal transmission. The fMAPK pathway required an intact SEA domain and the cytosolic tail, consistent with a cis signaling mechanism that required structural coupling across the membrane. In contrast, the HOG pathway functioned independently of the cytosolic tail and tolerated misfolded SEA domain variants, consistent with trans signaling mediated by extracellular domains of interacting partners. The HOG pathway may detect misfolding as part of its sensing mechanism, as stressors that induce protein misfolding required Msb2 for survival. This work reveals how differential tolerance to protein deformation confers signaling specificity and identifies sensor deformation as a general feature of mechanosensory pathways that respond to environmental stress.

**HIGHLIGHTS:**

- Signaling pathways differ in tolerance to misfolding of a sensory domain
- Misfolded SEA domains retain function in a stress pathway (HOG) pathway but not a cell differentiation pathway (fMAPK)
  - Misfolded SEA domain variants showed altered protein levels, mis-localization in the secretory pathway, and turnover by ERAD
  - Non-functional variants lacked residues that stabilize the structure through intramolecular bonds
- Differential tolerance for misfolding revealed distinct modes of signaling
  - Trans signaling predominated in the HOG pathway and did not require proper SEA domain folding or the mucin cytosolic tail
    - A dominant hyperactive variant next to the SEA domain revealed basal interactions with the CR domain of tetraspanin
    - AlphaFold modeling showed distinct interactions occur between the SEA domain and tetraspanin in the basal and activated states
  - Cis signaling predominated in the fMAPK pathway
    - Required a properly folded SEA domain and conformational coupling to the cytosolic tail
    - Yapsin processing was required for SEA domain activation and turnover of the mucin cytosolic tail
- HOG pathway may sense protein misfolding as part of its activation mechanism.
- SEA domains are conserved throughout fungal mucins and mammalian glycoprotein sensors suggesting a generalizable mechanism
- Protein deformation may provide information to survival pathways about environmental stress.

**GRAPHICAL ABSTRACT:** 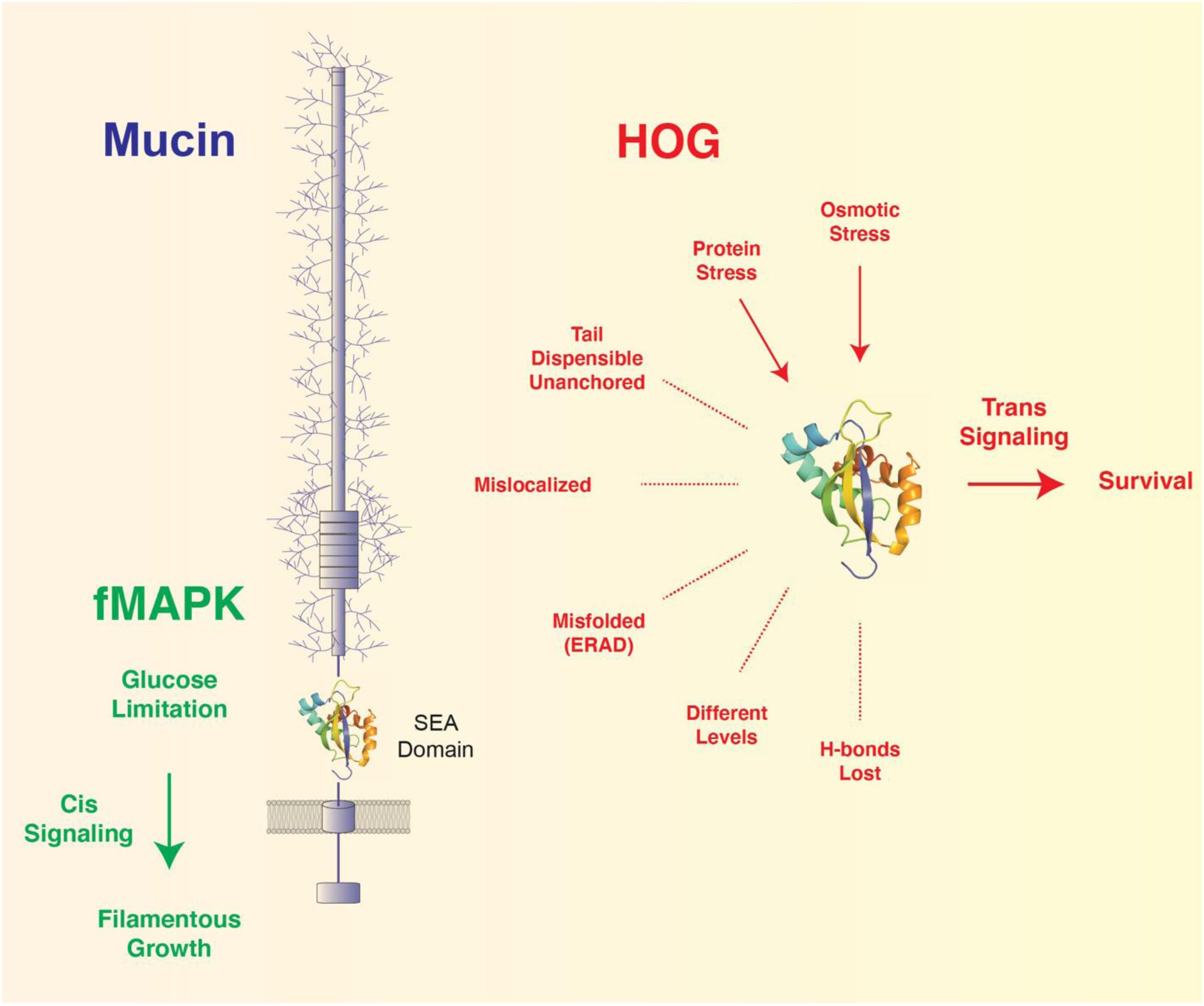

Signaling pathways often share components yet activate different effector processes through mechanisms that remain unclear. The same mucin regulates two MAPK pathways (red and green), and the discovery of a conserved SEA domain provided insights into specificity mechanisms. In the fMAPK pathway that regulates filamentous growth, the mucin works in a classical manner, where an external signal (in this case underglycosylation by glucose limitation) transduces a signal to the cytosolic domain in cis. By comparison, the HOG pathway that responds to osmotic stress displayed a remarkable tolerance for mucin and SEA domain deformation. Protein variants that caused SEA domain misfolding, mislocalization, and degradation by ERAD retained function in the HOG pathway. Truncations that removed the cytosolic tail and transmembrane anchor were also functional. These phenotypes support a trans activation mechanism with external partners that was preferential for activation of the HOG pathway. SEA domain deformation may be induced by environmental stress as a trigger for the HOG pathway. Cells may detect misfolding of protein domains to gain information about environmental stress.

## INTRODUCTION

Protein folding is critical for protein function (Rao and Bredesen 2004; Gidalevitz *et al*. 2011; Moreno-Gonzalez and Soto 2011). Protein folding occurs by intrinsic mechanisms to minimize entropic states (Anfinsen 1973; Dill *et al*. 2008; Senet *et al*. 2026) and can be facilitated by chaperones (Taipale *et al*. 2010). Proteins can become misfolded by damage due to genetic lesions and environmental stress. Protein misfolding is an underlying cause of neurological disorders and other diseases (Sweeney *et al*. 2017; Kuzu *et al*. 2025; Tregub *et al*. 2025). Although much effort has been devoted to understanding the mechanisms of protein folding, and the detrimental consequences affiliated with protein misfolding, relatively little is known about the functional capacities of proteins in their partially folded states. Here, we describe an example where protein misfolding of a sensory domain leads to differential signaling of one MAPK pathway over another.

Signaling pathways allow the detection and response to extracellular stimuli and are critical for survival. One feature of signaling pathways is their ability to share components yet generate highly specific biological responses (Rood *et al*. 2025). Conserved Mitogen-Activated Protein Kinase (MAPK) pathways regulate distinct biological processes, like the response to stress (Mordente *et al*. 2024) and cell differentiation (Sun *et al*. 2015; Peterson *et al*. 2022), yet function in networks containing common or shared protein modules (Bandyopadhyay *et al*. 2010; Johnson 2011; Bahar *et al*. 2023). How specificity is achieved in interconnected networks is a fundamental question in biology. In addition to producing the correct response, signaling systems must avoid inappropriate activation arising from fluctuations in protein levels, localization, or folding states. The loss of specificity by erroneous crosstalk is an underlying cause of cancer and other diseases (Brognard and Hunter 2011; Arthur and Ley 2013; Dalton 2013; Yuan *et al*. 2022; Hendrikse *et al*. 2023; Ng *et al*. 2024). Therefore, understanding how signaling pathways balance specificity with robustness to perturbation is a highly relevant question.

Like other eukaryotes, the budding yeast *Saccharomyces cerevisiae* contains evolutionarily conserved MAPK pathways that share components yet induce distinct responses (Schwartz and Madhani 2004; Wang and Dohlman 2004; Bardwell 2005; Saito 2010). For example, the same signaling mucin Msb2 regulates the fMAPK and HOG pathways. Mucins are highly glycosylated proteins with variable repeats rich in proline, threonine, and serine residues [PTS repeats (Borneman and Pretorius 2015; Dvela-Levitt *et al*. 2019; Kiss *et al*. 2019)]. Mucins also contain cytosolic tails that regulate effector pathways. Unlike G-protein coupled receptors (GPCRs), which typically bind to specific ligands (Velazhahan *et al*. 2021; Velazhahan *et al*. 2022), mucins function as mechanoreceptors that in an ill-defined manner detect mechanical perturbations, such as changes in pressure or hydration (Delarue *et al*. 2017; Hansson 2020). Misregulated mucins in humans cause toxic proteinopathies and are leading cause of diseases like cancers and kidney disease, which makes their study a priority (Sorensen *et al*. 2006; Bafna *et al*. 2010; Kosugi *et al*. 2011; Kirby *et al*. 2013; Kufe 2013; Kharbanda *et al*. 2014; Mukamel *et al*. 2021; Qing *et al*. 2022; Zhou *et al*. 2022; Xi *et al*. 2025).

One pathway in yeast that is regulated by Msb2 is the filamentous/invasive/pseudohyphal growth (fMAPK) pathway that controls differentiation to elongated and interconnected adhesive cells suited for nutrient foraging (Ryan *et al*. 2012; Kumar 2021). In pathogens like *Candida albicans*, filamentous growth is required for virulence (Lo *et al*. 1997; Swidergall *et al*. 2015a; Woolford *et al*. 2016). During filamentous growth, Msb2 responds to glucose limitation, which may result from changes in its glycosylation state (Karunanithi *et al*. 2010; Adhikari *et al*. 2015b). By comparison, Msb2 also regulates the High Osmolarity Glycerol response (HOG) pathway that controls the response to osmotic stress (Hohmann 2002; O’rourke and Herskowitz 2004; Tatebayashi *et al*. 2007; Yamamoto *et al*. 2015; Nishimura *et al*. 2016). The HOG pathway is composed of redundant branches that converge on a common MAPKK (Maeda *et al*. 1995; Posas *et al*. 1996; Posas and Saito 1997). A second mucin, called Hkr1, functions with Msb2 in the HOG pathway but does not regulate the fMAPK pathway (Pitoniak *et al*. 2009). It is not known what signal the mucins are sensing in the HOG pathway or how they induce specific responses. This puzzle is complicated because the HOG pathway is induced by multiple stimuli that are seemingly unrelated at the molecular level, including salt, temperature, pressure, and heavy metals.

Although the sensing and specificity mechanisms of yeast mucins remain elusive, several regulatory features have been defined. Msb2 contains a positive regulatory region (PRR) and negative regulatory region (NRR) with an inhibitory function (Cullen *et al*. 2004; Tatebayashi *et al*. 2007; Vadaie *et al*. 2008; Prabhakar *et al*. 2021). Proteolytic processing and release of the NRR is required for activation of the protein. Many other signaling glycoproteins are also proteolytically processed (Andersson *et al*. 2005; Johansson *et al*. 2008; Sandberg *et al*. 2008), and several exhibit a cleavage-dependent activation mechanism (Schoenwaelder *et al*. 1997; Logeat *et al*. 1998; Moloney *et al*. 2000; Chen *et al*. 2024). The cleavage of Msb2 is mediated by yapsins, aspartyl-type proteases that resemble sheddases in mammals (Kheradmand and Werb 2002; Krysan *et al*. 2005; Lichtenthaler *et al*. 2018). Msb2 also functions in a complex with proteins that resemble the tetraspanin adaptor in mammals (Maecker *et al*. 1997; Gueho *et al*. 2025). One of these proteins is Sho1, a four-span transmembrane protein with a cytosolic SH3 domain (Maeda *et al*. 1995; Cullen *et al*. 2004; Vadaie *et al*. 2008). Another is Opy2, which contains an extracellular cysteine-rich (CR) domain that forms intramolecular disulfide bonds (Wu *et al*. 2006). Opy2 also bears resemblance to fungal CFEM proteins, which play critical roles in fungal development and pathogenesis (Zhang *et al*. 2015). Msb2, Sho1, and Opy2 function in the fMAPK and HOG pathways. Therefore, the same mucin complex detects different signals and activates different MAPK pathways, providing a model for understanding signaling specificity (Posas *et al*. 1998; Wu *et al*. 1999; Jansen *et al*. 2001).

To explore how Msb2 regulates different MAPK pathways in pathway-specific contexts, a structure-guided approach was utilized, which identified conserved SEA domains in yeast and other fungal mucins. Functional analysis of the SEA domain of Msb2 identified protein misfolding as a critical point of discrimination between MAPK pathways. The fMAPK pathway required a properly folded SEA domain, whereas the HOG pathway showed broad tolerance for misfolded SEA domain variants present at different levels, which accumulated and were turned over in the endomembrane system. Specificity was also conferred by the mucin cytosolic tail, which was required in the fMAPK pathway but was dispensable in the HOG pathway. These observations revealed different activation mechanisms: the fMAPK pathway predominately functioned in cis through the cytosolic tail, whereas the HOG pathway operated in trans through interactions with tetraspanin. Protein misfolding may also be part of the sensing mechanism of the HOG pathway, which can account for its ability to detect multiple unrelated molecular stimuli. Protein domain deformation may be a general way that survival pathways detect and respond to environmental stress.

## RESULTS

### Discovery of SEA domains in fungal mucins as MAPK pathway regulatory modules

To understand how Msb2 regulates different MAPK pathways that detect and respond to different stimuli (**Fig. 1A**), a structure-guided approach was employed. The structure prediction program AlphaFold (Jumper *et al*. 2021; Varadi *et al*. 2024) identified a globular domain in the extracellular region of Msb2 next to the transmembrane helix (**Fig. 1A-B**, dashed box). The 110 amino acid (aa) domain had a per-residue confidence score of >90, denoting a high-accuracy prediction (Movie 1, blue, pLDDT score). The domain resembled conserved SEA domains found in the human mucin MUC1 (**Fig. 1B**) and other signaling glycoproteins [Movie 2A-B, *Fig. S1*, (Macao *et al*. 2006; Noguera *et al*. 2020)]. The SEA domain of Msb2 had a ferredoxin-like fold with a ßαßßαα arrangement that was similar to the ßαßßαß pattern of MUC1 (**Fig. 1B**). In both domains, the ß strands formed an anti-parallel arrangement creating a concave surface packed against the α helices in a dome-like structure. When overlaid, the SEA domain of Msb2 overlapped with the SEA domains of other signaling glycoproteins (*Fig. S2*, TM-scores >0.5).

**Figure 1.**
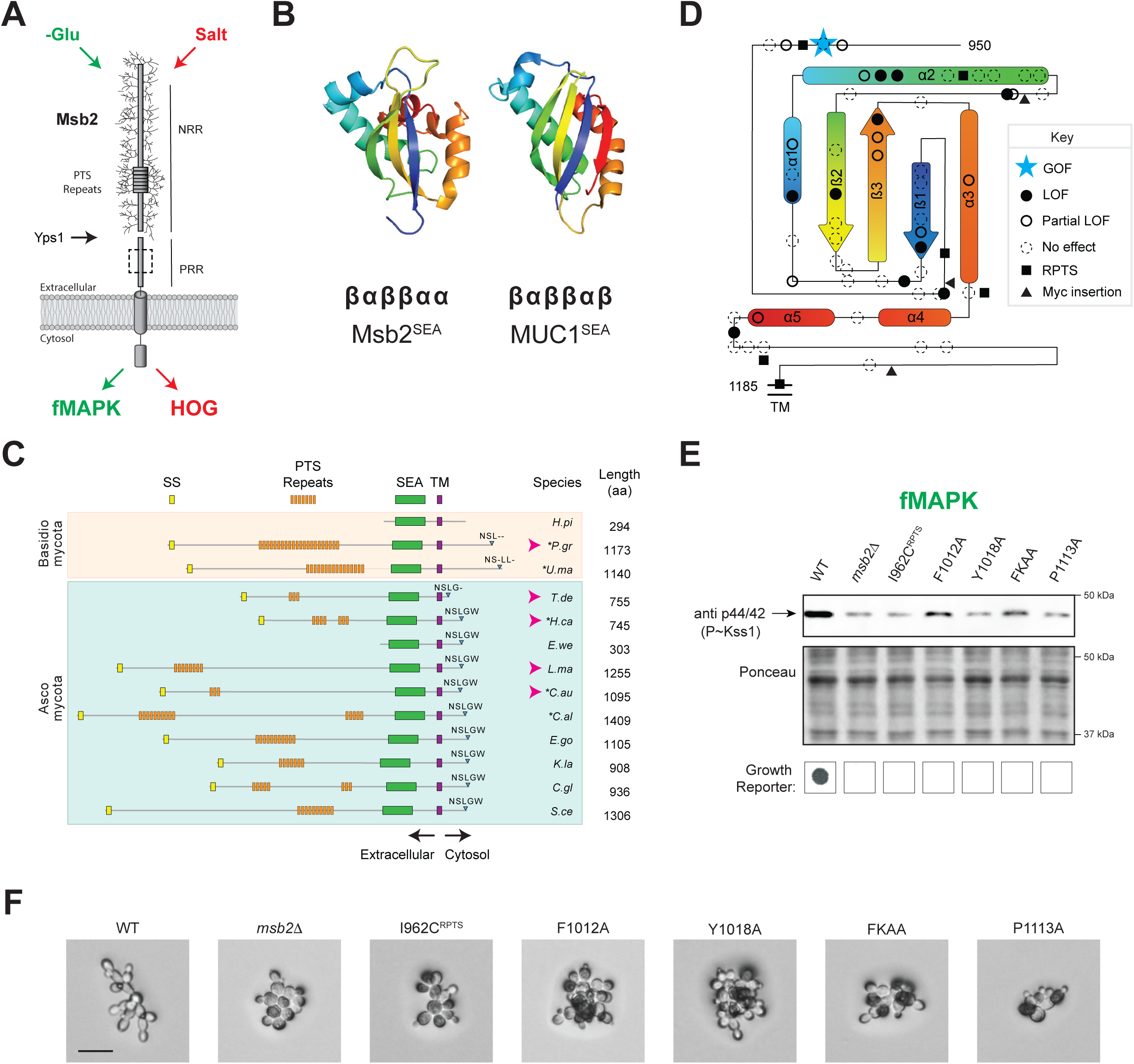
Discovery of SEA domain of Msb2 and analysis of its role as a MAPK pathway regulatory module. **(A)** Diagram of the Msb2 protein with the glycosylated extracellular region containing the PTS-rich repeats, approximate cleavage site of Yps1, positive regulatory region (PRR), and negative regulatory region (NRR) labelled. Msb2 regulates the fMAPK and HOG pathways. Dashed square, location of the SEA domain. **B)** AlphaFold models of the SEA domains of *S. cerevisiae* Msb2 (1000-1110 aa residues, AFDB# P32334) and human MUC1 (residue# 1039-1148, AFDB# P15941). The rainbow color shows the N-terminus (blue) to C-terminus (red). ß, beta sheet, α, alpha helix. **(C)** Fungal proteins containing SEA domains. Pink arrowheads, previously unidentified Msb2 homologs. Signal sequence (SS, yellow), S/T/P-rich repeat (orange), SEA domain (green), TM domain (purple), and cytoplasmic tail with conserved NSLGW sequence (blue) are shown. Dashes, residues different from the NSLGW sequence. Species abbreviations, *H.pi. Hydnomerulius pinastri, P.gr. Puccinia graminis, U.ma. Ustilago maydis, T.de. Taphrina deformans, H.ca. histoplasma capsulatum, E.we. Escovopsis weberi, L.ma. Lophiostoma macrostomum, C.au. Candidozyma auris*, *C.al. Candida albicans, E.go. Eremothecium gossypii, K.la. Kluyveromyces lactis, C.gl. Candida glabrata, and S.ce. Saccharomyces cerevisiae*). Asterisks, pathogenic species. Proteins without PTS repeats were not considered mucins, as they lack repeat regions. **(D)** Two-dimensional diagram of the SEA domain of Msb2. Residues tested in the study are marked by filled circles, not functional; open circles, partial loss of function; dashed circles, no phenotype detected; blue star, hyperactive allele. Loops are not to scale. Triangles, Myc insertions; squares, repeated sequences. **(E)** fMAPK analysis by phosphoblot showing P∼Kss1 levels by p42/44 antibodies compared to total protein levels detected by PonceauS staining. Below, *ste4 FUS1-HIS3* reporter gene for fMAPK activity **(F)** Filaments of the designated alleles by 20X microscopy. Bar, 10 microns.

SEA domains have not previously been identified in fungal species, perhaps due to low sequence conservation (*Fig. S3-S4A*). To determine whether SEA domains are present in other fungal species, the structural search algorithm Foldseek was applied (Van Kempen *et al*. 2024). Foldseek identified SEA domains in nearly 200 fungal species across two major phyla of fungi. SEA domains were uncovered in the Msb2 homolog in *Candida albicans*, a major human fungal pathogen (Swidergall *et al*. 2015b; Richardson 2022), and other relatives of *S. cerevisiae*. By searching for SEA domains, Foldseek identified new mucin homologs in *Candidozyma auris,* an emerging hospital pathogen (Mishra *et al*. 2023; Kim *et al*. 2024), *Histoplasma capsulatum*, a distantly related human pathogen (Valdez *et al*. 2022), and *Puccinia graminis*, a plant pathogen (Guo *et al*. 2022) (**Fig. 1C**, pink arrows). Fungal SEA domains had overlapping structures with high TM-scores (*Fig. S4B*, 0.87-0.96, *Movie 3*) and the same pattern of hydrophobic residues that occur in the SEA domains of other organisms [*Fig. S3*, (Pei and Grishin 2017)].

The SEA domain of Msb2 resides in the PRR region, which is critical for MAPK pathway signaling (**Fig. 1A**). Epitope fusions constructed in and around the SEA domain were defective for fMAPK pathway activity (*Fig. S5*) suggesting an important functional role. To more precisely map functional residues in the SEA domain, site-directed mutagenesis was performed by substituting residues of interest to alanine (**Fig. 1D**). In particular, SEA domain variants were generated in a functional epitope-tagged version of Msb2 expressed from the endogenous promoter (pHA-MSB2). SEA domain variants were evaluated for fMAPK pathway activity by measuring phosphorylation of the MAP kinase that regulates the fMAPK pathway by immunoblot (IB) analysis (**Fig. 1E**, top, Kss1). fMAPK pathway activity was also evaluated by a growth reporter, where growth on histidine-deficient media provides a readout of fMAPK pathway activity (**Fig. 1E**, *ste4 FUS1-HIS3,* bottom, *Fig. S6*). As a phenotypic test, fMAPK pathway activity was evaluated by examination of filament formation by microscopy (**Fig. 1F**, *Fig. S7A*). In total, 56 residues in the SEA domain and flanking regions were constructed and tested (**Fig. 1D**). Fifteen of the thirty-nine residues in the SEA domain (38%), and four of the seventeen residues in the flanking regions (25%) had a detectable phenotype (**Fig. 1D-F**, hyperactive, partial or complete loss of function), demonstrating a central role for the SEA domain in MAPK pathway regulation.

### SEA domain variants show MAPK pathway-specific phenotypes that correspond to defects in protein stability

Msb2 regulates MAPK pathways that control filamentous growth (fMAPK) and the response to osmotic stress (HOG). To identify pathway-specific features of the SEA domain, variants were compared by tests that evaluate the fMAPK (see **Fig. 1E-F**) and HOG pathways. To evaluate the HOG pathway, a strain was constructed that lacked the redundant mucins (Msb2 and Hkr1) and the redundant branch of the HOG pathway (Ssk1). The *msb2Δ hkr1Δ ssk1Δ* triple mutant was evaluated for the HOG pathway by phosphorylation of the MAPK Hog1 (see below) and growth on high osmolarity media (1M KCl). Compared to cells lacking Msb2 (*msb2Δ*), cells harboring the pHA-MSB2 plasmid (WT) complemented the signaling defects of the fMAPK (**Fig. 2A**, fMAPK) and HOG pathways (**Fig. 2A**, HOG). Several variants that were defective for the fMAPK pathway were also defective for the HOG pathway (**Fig. 2A**, e.g. I1055A). Other variants that were defective for the fMAPK pathway were functional in the HOG pathway (**Fig. 2A**, e.g. F1012A, *Fig. S8*). None of variants were only defective for the HOG pathway. The fMAPK and HOG pathways are also required for the response to endoplasmic reticulum (ER) stress, which can occur when misfolded proteins accumulate in the secretory pathway (Torres-Quiroz *et al*. 2010). ER stress can be induced by the N-linked glycosylation inhibitor tunicamycin (Oslowski and Urano 2011). In media supplemented with tunicamycin, SEA domain variants showed the same growth defects as seen for the fMAPK pathway (*Fig. S9*). This result highlights the differences in SEA domain function between the fMAPK and HOG pathways. Two N-linked glycosylation sites (N-X-S/T motif) that were tested were dispensable for Msb2 function (*Fig. S10A*, green), indicating that sensitivity to glycosylation stress occurs outside the SEA domain. These results reveal different roles for the SEA domain in the fMAPK and HOG pathways.

**Figure 2.**
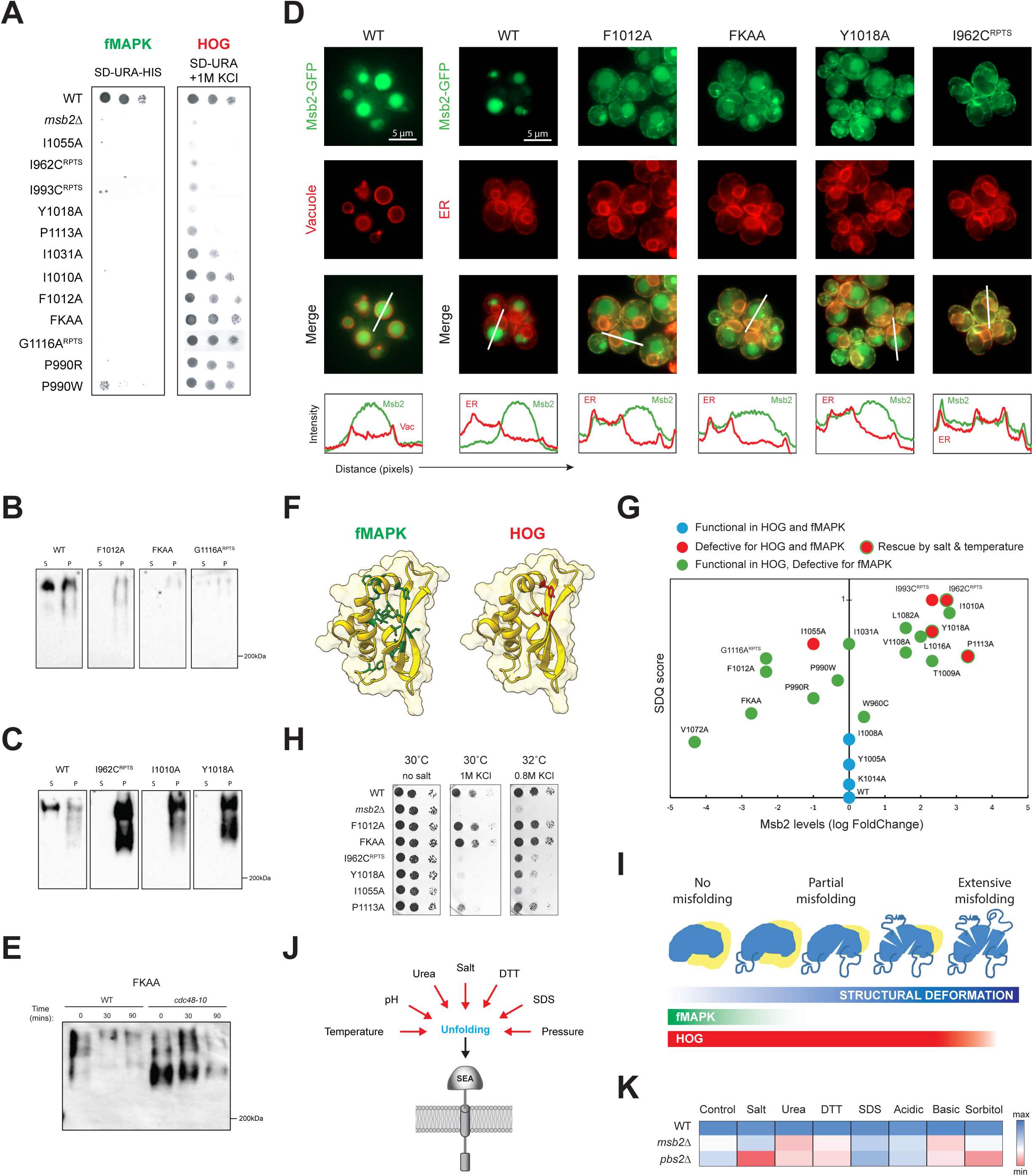
SEA domain variants with altered fMAPK and HOG pathway activity show altered protein levels, are mis-localized and accumulate in ERAD mutants. **(A)** Left, role of SEA domain variants in regulating the fMAPK pathway by the Msb2-dependent *FUS1-HIS3* growth reporter. Right, role of SEA domain variants in regulating the HOG pathway based on sensitivity to 1M KCl. Strain was *msb2Δ hkr1Δ ssk1Δ* (PC8102) containing positive control (pHA-Msb2 for WT) or negative control (pRS316 for *msb2Δ*). The serial dilutions were assembled from different plates incubated for the same duration. Refer to *Fig. S6* and *Fig. S8* for complete dataset with controls. **(B)** IB of supernatant (S) fractions for shed Msb2 and Pellet (P) fractions for cell-associated Msb2 of SEA domain variants found at lower levels in the cell. **(C)** IBs of SEA variants accumulating in cell pellets. Full set can be found in *Fig. S12.* (D) Top, co-localization of the indicated SEA domain variants (Green) with proteins that localize to the ER (Elo3, mCherry channel) and vacuole (Vph1, mCherry channel) as indicated. Merge represents superimposed pictures. Bar, 5μm. Line plots. White line in Merge shows the region analyzed for intensity values in the mCherry and GFP channels. **(E)** Levels of SEA domain variant FKAA in wild-type cells (WT) and the *cdc48-10* mutant incubated for the indicated times at 37°C after treatment with cycloheximide. Time 0, 30°C. Blots are probed with anti-HA antibodies, and the MWM at 200 kDa is marked. **(F)** AlphaFold models showing distribution of residues in the SEA domain required for fMAPK (green) and HOG (red) pathway signaling. **(G)** Plot showing Msb2 cell-associated levels (fold changes are shown as log values) and SDQ (capped at 1, refer Methods) scores. See the key for details. **(H)** Salt sensitivity rescue of HOG defective variants at elevated temperature (32°C) in the *msb2Δ hkr1Δ ssk1Δ* strain. **(I)** Model for the function of Msb2-dependent MAPK pathways in response to structural deformation of the SEA domain. Increasing structural deformations are tolerated by the HOG pathway, while preventing activation of the fMAPK pathway. At high structural deformations, both pathways are impacted. **(J)** Environmental stresses known to compromise protein folding may misfold SEA domain acting as trigger for HOG pathway activation. **(K)** Heat map generated from growth assays for strain with (WT) or without Msb2 (*msb2Δ*) and *pbs2Δ* under various proteostatic stresses such as Urea, DTT, SDS, pH. The *msb2Δ* is the *msb2Δ hkr1Δ ssk1Δ* strain.

To define the basis for differences in SEA domain function in the two MAPK pathways, the levels of the HA-Msb2 protein containing SEA domain variants were examined. Msb2 is proteolytically processed, and the NRR is shed from cells (Vadaie *et al*. 2008). Each variant with altered MAPK pathway activity was examined for shedding of HA-Msb2 by spot immunoblot (spotIB) analysis. By this test, nearly 40% of variants showed reduced shedding compared to wild-type HA-Msb2 (*Fig. S11*). Standard IB analysis revealed two groups of variants when comparing shed HA-Msb2 (supernatants, S) to cell-associated HA-Msb2 (pellets derived from cell extracts, P). One group showed lower overall levels of the HA-Msb2 protein (**Fig. 2B**, *Fig. S12*), which may be due to defects in protein stability. The other group showed high levels of HA-Msb2 in pellet fractions and were not shed, which may result from accumulation of the protein in the secretory pathway (**Fig. 2C**, *Fig. S12*). Both classes of variants may result from improper folding of the SEA domain. The fact that SEA domain variants with altered stability were defective in the fMAPK pathway was not surprising. However, most of these variants were functional in the HOG pathway (**Fig. 2A**, HOG). These included variants present at >5-fold lower levels than wild type (P990R, F1012A, FKAA, and G1116A^RPTS^) and variants present at >10-fold higher levels that were not shed (I1010A). These results indicate that the HOG pathway is more tolerant to defects in the stability of the SEA domain than the fMAPK pathway.

### SEA domain variants that function in the HOG pathway accumulate in the ER and are turned over by ERAD

We next examined the localization of SEA domain variants with altered protein levels. As a secreted glycoprotein, Msb2 is translated in the ER and delivered to the plasma membrane by the secretory pathway. By localization, wild-type Msb2-GFP is mainly found in the vacuole/lysosome due to rapid turnover from the plasma membrane (Vadaie *et al*. 2008; Adhikari *et al*. 2015a; Adhikari *et al*. 2015b). Co-localization of Msb2-GFP with a vacuolar protein marker, Vph1- mCherry, confirmed that Msb2 is mainly present in the vacuole (**Fig. 2D**, far left, see Merged Image and Intensity plot). In contrast, wild-type Msb2 was not enriched in the cortical and perinuclear compartments of the ER based on co-localization with the ER marker Elo3-mCherry (**Fig. 2D**, second panel from left). Compared to wild-type Msb2, the SEA domain variants with altered protein levels showed localization in the ER (**Fig. 2D**, variants compared with the ER marker). These included variants present at low levels, like F1012A and FKAA, and variants that accumulated at high levels, like Y1018A and I962C^RPTS^. Retention of these variants in the ER indicates that the variants are improperly folded. Despite their folding/localization defects, the variants retained function in the HOG pathway (**Fig. 2A**, FKAA and F1012A).

Misfolded proteins in the endomembrane system are turned over by quality control pathways such as ER-associated degradation [ERAD, (Lippincott-Schwartz *et al*. 1988; Klausner and Sitia 1990; Vembar and Brodsky 2008)]. As an independent test of protein misfolding, SEA domain variants were examined for turnover by ERAD. HA-Msb2 levels were examined in the temperature-sensitive *cdc48-1* mutant (Rabinovich *et al*. 2002; Tran and Brodsky 2012), which has a conditional defect in the ATPase that exports misfolded proteins from the ER for destruction by the 26S proteosome (Bays *et al*. 2001; Ye *et al*. 2001; Jarosch *et al*. 2002). SEA domain variants introduced in the *cdc48-10* mutant were examined at semi-permissive (30°C) and non-permissive (37°C) temperatures, with protein production being inhibited by cycloheximide. SEA domain variants accumulated in the *cdc48-10* mutant based on IB analysis (*Fig. S13A*, 30°C) and protein localization (*Fig. S13B*). For example, variant FKAA accumulated in the *cdc48-10* mutant compared to a wild-type strain (**Fig. 2E**, wild type). FKAA levels were reduced at 90 mins, which may be due to residual Cdc48 function at 37°C or turnover by a Cdc48-independent mechanism. Therefore, ERAD functions as a quality-control checkpoint for the SEA domain variants that accumulate in the secretory pathway. Accumulation of misfolded proteins in the ER might impact viability (Reiss *et al*. 2000), but no growth defects were observed, even under conditions that induce ER stress (*Fig. S14*). Several variants recognized as mis-folded proteins by ERAD were functional in the HOG pathway (**Fig. 2A**, FKAA and F1012A).

### SEA domain variants functional in the HOG pathway show predicted structural deformation

To gain structural insight into the differences between SEA domain function in the fMAPK and HOG pathways, functionally critical residues were mapped onto the predicted structure of the domain (**Fig. 2F**). Instead of clustering together most residues defective in the fMAPK pathway (60%, 9/15) were oriented towards the core of the structure (**Fig. 2F**, left, Movie 4A-B). Eight residues formed hydrogen bonds that when changed to alanine would be expected to destabilize the tertiary structure (*Fig. S10B*, yellow). Residues defective for both pathways were similarly buried in the structure (**Fig. 2F**, right). Residues dispensable in either pathway were located on the SEA domain surface (Movie 4C, pink). These results suggest that the functionality of the residues in the fMAPK pathway most likely stems from their contribution in maintaining the structural integrity of the domain. To further validate these findings, residues in the SEA domain were examined by MutateX, a FoldX based platform which predicts the folding free energy change (ΔΔG) for each amino acid change in a structure (Schymkowitz *et al*. 2005; Tiberti *et al*. 2022). Based on the average ΔΔG scores of each residue generated by MutateX, we identified residues important for structure (*Fig. S15*) that partially overlapped with conserved residues (*Fig. S16*). By these criteria, most residues important for the structure of the SEA domain were required for fMAPK pathway activity (80%, 12/15) supporting the idea that the structural integrity of the domain is critical for its function in the fMAPK but not HOG pathways.

To further examine the folding landscape of the SEA domain, a numerical score was generated for each variant, called the structural deformation quotient (SDQ). The relative defect in shedding of these variants was combined with the predicted structural factor (ΔΔG) as an estimate of the degree of structural deformation. Plotting SEA domain variants by SDQ scores to protein levels, we found that SEA domain variants that had low SDQ scores and were present at normal levels were functional in both MAPK pathways (**Fig. 2G**, blue circles). By comparison, variants functional in the HOG pathway tolerated a wide range of SDQ scores and changes in protein levels (green circles) and were functional in the HOG pathway. Non-functional variants in the HOG pathway had high SDQ scores and wide differences in protein levels (red circles). These results highlight the differential tolerance of the HOG and fMAPK pathways to deformation of the SEA domain. We also uncovered conditions in which three of the five SEA domain variants were at least partially functional (**Fig. 2G**, red circles with green outlines). Specifically, the partially functional variants were Y1018A, and P1113A, and I962C^RPTS^, which contained an insertion of ∼ 300 aas, accumulated to >20-fold higher levels in the cell, and co-localized tightly with an ER marker. Similarly, the signaling defect of P1113A was also rescued by intragenic loss of the NRR region of Msb2 in the HOG pathway (*Fig. S17A*). Only one variant, I1055A, was completely non-functional in the HOG pathway across all tested conditions (**Fig. 2H**) and may represent a key residue as discussed below. These tests demonstrate an extraordinary tolerance by the HOG pathway for structural deformation of the SEA domain (**Fig. 2I**).

### Exploring the folding landscape by environmental perturbation reveals a sensory role for the HOG pathway to inputs that compromise protein folding

In yeast and other fungal species, the HOG pathway is activated by diverse stresses, including osmotic stress (Brewster *et al*. 1993), high pressure (Delarue *et al*. 2017), heat stress (Winkler *et al*. 2002; Saraswat *et al*. 2016), ER stress (Bicknell *et al*. 2010; Adhikari and Cullen 2014) and exposure to heavy metals (Enjalbert *et al*. 2006; Thorsen *et al*. 2006; Yaakoub *et al*. 2023). Each of these stimuli are also known to compromise protein folding (Devi *et al*. 2022; Omkar *et al*. 2025). We therefore hypothesize that the HOG pathway may detect these unrelated stimuli by sensing defects in folding of the SEA domain (**Fig. 2J**). To test this possibility, the type cells (WT) and cells lacking Msb2 (*msb2Δ*) were compared to a HOG pathway mutant (*pbs2Δ*) across a panel of stresses specifically designed to compromise protein folding/stability. Exposure of cells to protein denaturing agents including urea, acidic and basic pH, detergents (sodium dodecyl sulfate, SDS), and reducing agents (dithiothreitol, DTT) induced growth defects that were ameliorated by Msb2 and the HOG pathway (**Fig. 2K**). These results demonstrate a role for the HOG pathway in the response to compromised protein folding.

The folding landscape of the SEA domain was also examined in cells grown in different conditions. Gene by environment (G X E) interactions can reveal unexpected functional relationships. Different concentrations of KCl (0.05, 0.2, 0.5, 0.8, and 1M) and a range of temperatures (26°C, 30°C, 32°C, 35°C) were varied in a phenotypic matrix. Combinatorial changes to salt and temperature were also examined, which revealed a wide range of Msb2-dependent function. Furthermore, we discovered an inverse sensitivity between salt and temperature, suggesting no single inducer is responsible for activation of the HOG pathway (**Fig. 2H**). These findings show that Msb2 in the HOG pathway detects and responds to a range of protein folding stress. Thus, SEA domain misfolding may be part of the sensory mechanism for Msb2 function in the HOG pathway.

### Mucin cytosolic tail is required by the fMAPK pathway but is dispensable for the HOG pathway

Yeast has two signaling mucins (Hohmann 2002; Tatebayashi *et al*. 2007; Pitoniak *et al*. 2009). Msb2 functions in the fMAPK and HOG pathways, and Hkr1 functions in the HOG pathway but not the fMAPK pathway. The mucins have the same topology but differ in primary aa sequence (*Fig. S18A*), and gene synteny (*Fig. S19*), and are not paralogs (Byrne and Wolfe 2005). Because Hkr1 does not function in the fMAPK pathway (*Fig. S20*), specificity determinants may exist that are unique to each mucin. AlphaFold analysis identified a SEA domain in Hkr1 that was similar to the SEA domain of Msb2, except for an extended loop (**Fig. 3A**, Movies 5-6, TM score 0.94). The loop may be dispensable in the HOG pathway because Msb2 functions in the HOG pathway without the loop. The SEA domain of Hkr1 may function in the HOG pathway in a similar manner as Msb2. For example, an Hkr1 SEA domain variant, Y1266A, showed similar salt sensitivity as the equivalent Y1018A variant of Msb2 (*Fig. S18B*). Moreover, like for Msb2, most SEA domain variants of Hkr1 retained function in the HOG pathway (*Fig. S18B*).

**Figure 3.**
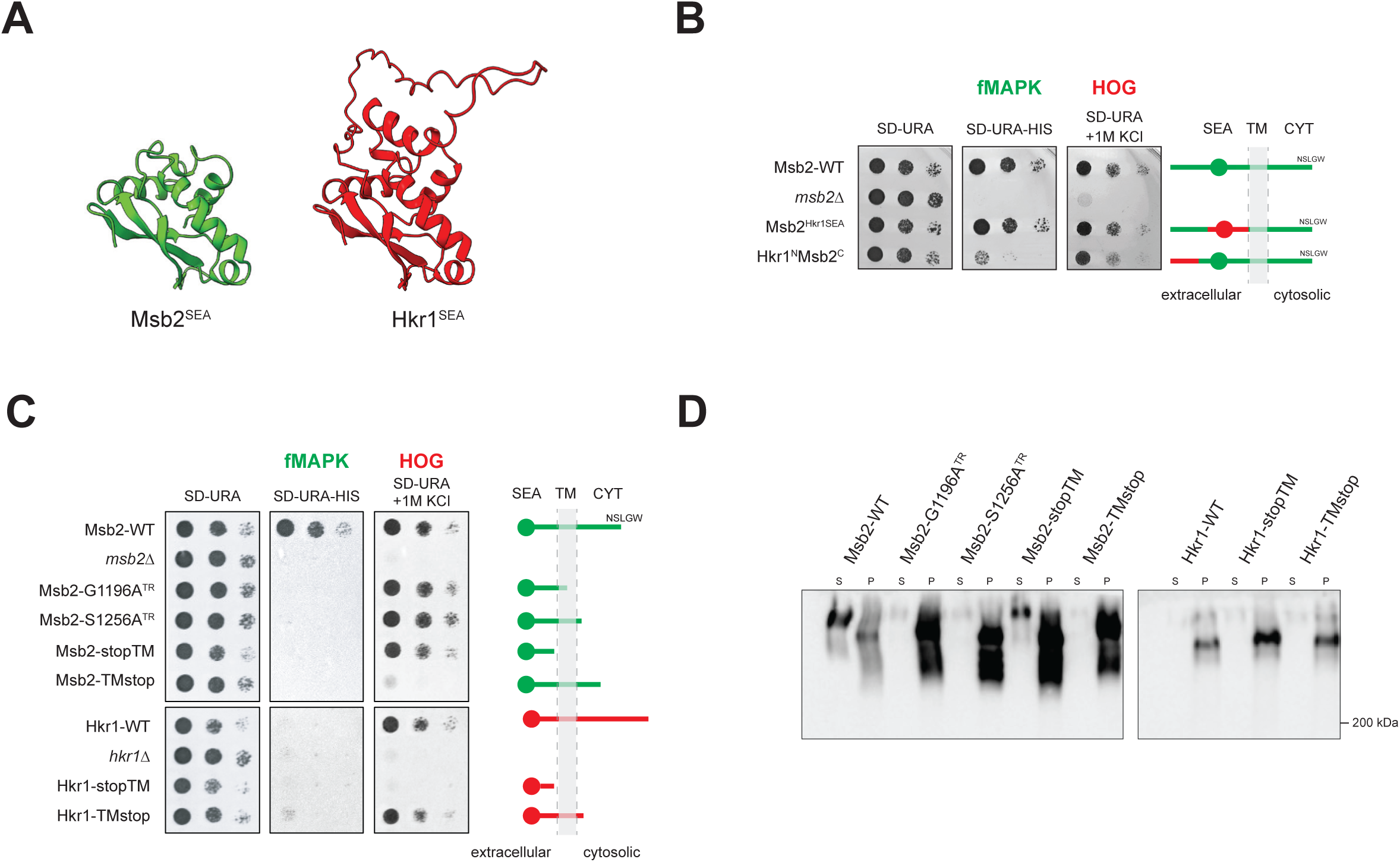
Role of cytosolic tails of signaling mucins in MAPK signaling. **(A)** AlphaFold models of Msb2 (green) and Hkr1 (Red) SEA domains. See also Movie 6. **(B)** Right, representation of Msb2-Hkr1 fusion proteins (portions of Msb2, Green and Hkr1, Red), Msb2^Hkr1SEA^ and Hkr1^N^-Msb2^C^. Left, function of the fusion proteins in the fMAPK (*FUS1-HIS3*) and HOG (1M KCl). See Fig. 2A for details. **(C)** Right, Representation of cytosolic deletion variants of Msb2 and Hkr1. Left, Function of the cytosolic deletion variants in the fMAPK (*FUS1-HIS3*) and HOG (1M KCl). **(D)** IB for supernatant and pellet fractions of cytosolic deletion variants. The *msb2Δ* and *hkr1Δ* are *msb2Δ hkr1Δ ssk1Δ* strain.

To uncover specificity determinants between the SEA domains of Msb2 and Hkr1, fusion proteins were constructed. In one fusion protein, the SEA domain and flanking regions of Msb2 were replaced with the SEA domain and flanking regions of Hkr1. Msb2^Hkr1SEA^ functioned in the fMAPK and HOG pathways (**Fig. 3B**), even though 45% of the residues were different between the two proteins in this region (*Fig. S18*). Therefore, amino acid sequence differences in the SEA domains of Msb2 and Hkr1 do not impart functional differences in the HOG pathway, suggesting that the SEA domain function instead depends on a common sequence motif or the domain architecture. Another fusion protein was constructed that contained the promoter and N-terminus (1-1197 aa) of Hkr1 and the C-terminus of Msb2. Hkr1^N^-Msb2^C^ was largely functional in the HOG pathway but only partially functional in the fMAPK pathway (**Fig. 3B**). The reduced function of Hkr1^N^-Msb2^C^ in the fMAPK pathway may be due to lower expression of the chimeric mucin by the *HKR1* promoter. The HOG pathway is tolerant of low mucin levels (**Fig. 2G**), and HA-Hkr1 is present at 3-fold lower levels than HA-Msb2 yet functional in the HOG pathway [*Fig. S21A*, (Pitoniak *et al*. 2009)]. By comparison, *MSB2* is expressed at high levels, and its expression is induced by positive feedback by the fMAPK pathway (Cullen *et al*. 2004). The fact that low mucin levels are sufficient for HOG but not fMAPK provides yet another example of the broad tolerance of the HOG pathway for compromised mucin function.

Another difference between the mucins are their cytosolic tails, which are different lengths and dissimilar by aa sequence analysis. To examine the role of the tails, we first examined tail-less versions of the Msb2 and Hkr1 proteins. As expected from previous findings (Adhikari *et al*. 2015b), versions of Msb2 lacking a portion (S1256A^TR^) or the entire cytosolic tail (G1196A^TR^, Msb2-TMstop, Msb2-stopTM) were defective in the fMAPK pathway (**Fig. 3C**, fMAPK). The same versions of Msb2 were functional in the HOG pathway (**Fig. 3C**, HOG, except Msb2-TMstop). Therefore, the cytosolic tail of Msb2 is needed for the fMAPK pathway but dispensable for the HOG pathway. Additionally, based on Msb2-stopTM, a version lacking both the TM helix and cytosolic tail, the SEA domain of Msb2 did not need be tethered to the membrane to function in the HOG pathway. Like for Msb2, the cytosolic tail of Hkr1 was also dispensable in the HOG pathway (**Fig. 3C**, Hkr1-TMstop). Another version of Hkr1 lacking the TM helix and cytosolic tail (Hkr1-stopTM) was not functional perhaps because the TM domain of Hkr1 contributes to formation of the Hkr1-Sho1-Opy2 complex (Tatebayashi *et al*. 2015). Mucins lacking their cytosolic tails were defective in shedding to varying degrees and accumulated in cell pellets (**Fig. 3D**); however, this did not correlate with mucin function in the HOG pathway. The dispensability of the mucin tails highlights the broad tolerance of the HOG pathway to deformation of the mucin sensors compared to the fMAPK pathway.

### Cis- and trans-activation mechanisms are favored by different MAPK pathways

The finding that the cytosolic tail is dispensable indicates that Msb2 regulates the HOG pathway in trans by interaction with other proteins in the mucin complex. Msb2 (and Hkr1) form a complex with Opy2 and Sho1, proteins that resemble mammalian tetraspanin [**Fig. 4A**, (Tatebayashi *et al*. 2007; Adhikari *et al*. 2015a; Yamamoto *et al*. 2016)]. Specifically, on the extracellular side of the plasma membrane, the PRR region of Msb2, which includes site S1023, functions proximal to the cysteine-rich (CR) domain of Opy2 (Yamamoto *et al*. 2016). Placed in the context of the present study, this means that the SEA domain and N-terminal flanking region of Msb2 (called the Regulatory arm, 950-1000 aas) function next to the CR domain of Opy2 (**Fig. 4A**). AlphaFold multimer modelling of the mucin sensor complex supports this conclusion (*Fig. S22*).

**Figure 4.**
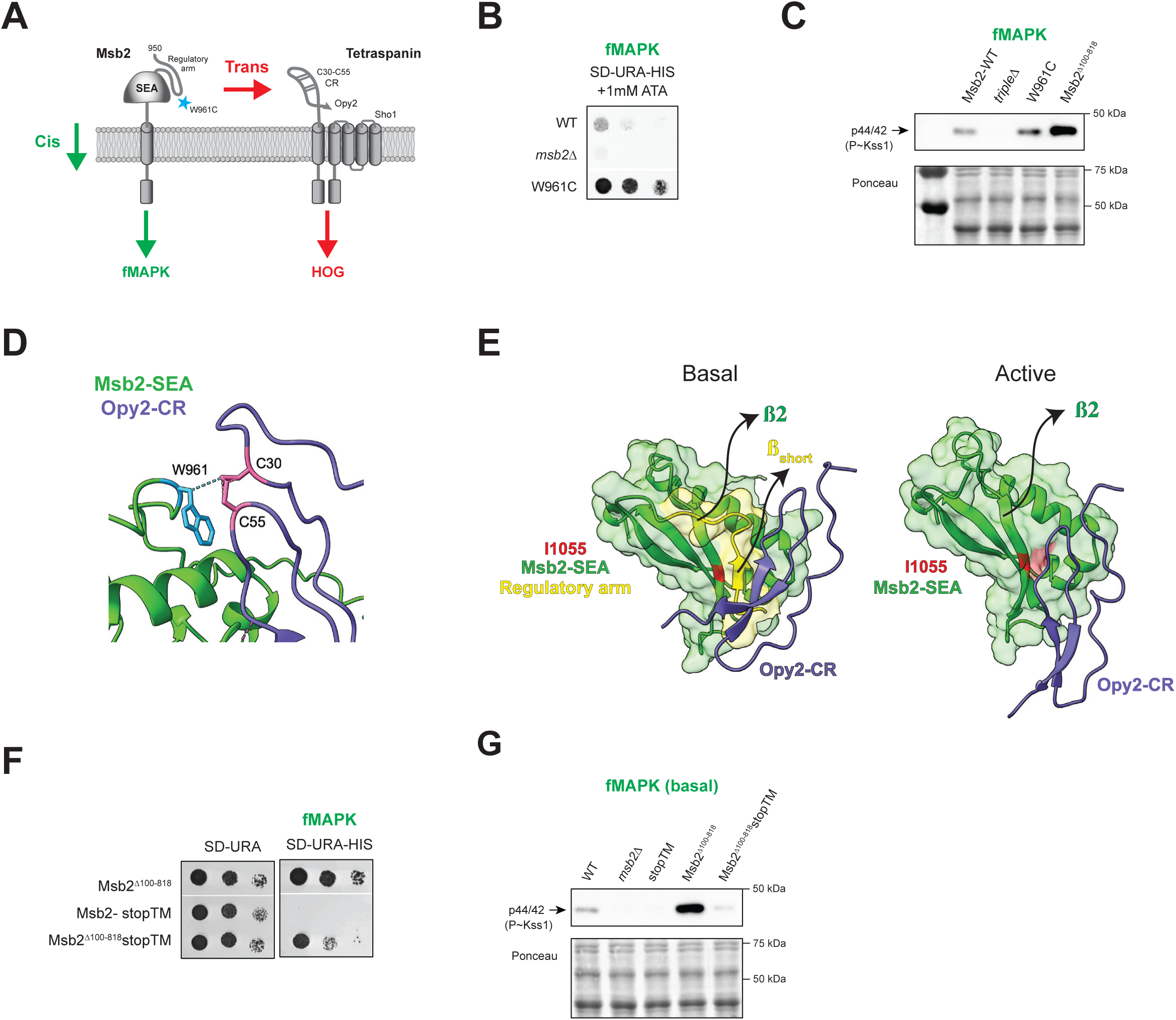
Msb2 functions in cis- and trans- based signaling in different pathways. **(A)** Schematic model showing cis- and trans- activation states. Blue star, Hyperactive allele. **(B)** *FUS1-HIS3* growth reporter assay for hyperactive allele W961C. **(C)** Left, P∼Kss1 blot for the indicated strains. *tripleΔ* indicates *msb2Δ hkr1Δ ssk1Δ*. Right, AlphaFold multimer model showing position of Msb2-W961 (green) with CR of Opy2 (purple) **(D)** Surface model of AlphaFold prediction of Msb2-SEA (green), and Regulatory arm (990-1000, yellow) and Opy2 (purple) interactions in basal or predicted activated states. Non bypassable variant I1055 (red) buried in the basal state/ exposed in the activated state. **(E)** Function in fMAPK shown by Growth reporter assays (*FUS1-HIS3*) for Msb2^Δ100-818/stopTM^. **(F)** P∼Kss1 blot for indicated strains. The *msb2Δ* is the *msb2Δ hkr1Δ ssk1Δ* strain.

In further support of this interaction, we identified a dominant variant in the regulatory arm that hyperactivates the fMAPK pathway. The W961C variant was identified by exploring variation of Msb2 across a large collection of wild and domesticated isolates (Peter *et al*. 2018). Among five single nucleotide variants (SNVs) that occur in and around the SEA domain in naturally occurring populations of yeast strains [resulting in changes W961C, P990A, P990H, T1006A, and F1012L (Vandermeulen *et al*. 2024)], W961C alone showed elevated fMAPK pathway activity and was the only hyperactive protein identified in the study. The W961C variant showed elevated activity of the fMAPK pathway based on the growth reporter (**Fig. 4B**) and phospho-IB analysis (**Fig. 4C**). The hyperactivity by W961C was less than the hyperactivity conferred by the loss of the NRR region (**Fig. 4C**, Msb2^Δ100-818^). AlphaFold modeling showed that W961 is located next to the disulfide pair C30-C55 of the CR domain of Opy2 (**Fig. 4D**, right, Movie 7). W961C may represent a highly specific change because substitution to alanine (W961A) or cysteine of adjacent residues (W960C, P963C, or T964C) did not hyperactivate the fMAPK pathway (*Fig. S6*).

We hypothesized that structural changes due to misfolding and shedding of the NRR region may alter the position of the regulatory arm and expose a latent interaction surface. This hypothesis is supported by the fact that osmotic stress alters the the CR domain relative to the PRR domain of Msb2 (Yamamoto *et al*. 2016), and by the location of the W961 residue in reference to the CR domain (**Fig. 4D**). To test this possibility, the SEA domain was modeled with and without the regulatory arm for interaction with the CR domain. In the basal state, a short ß-strand (ß_short_) in the regulatory arm interacted with the CR domain (**Fig. 4E**, Basal). No other interactions between these domains and the core SEA domain were observed. In the active state, modeled without the regulatory arm to mimic activation, the ß2 strand on the SEA domain was exposed and interacted with a separate region in the CR domain (**Fig. 4E**, Active). Interestingly, ß2 contained the single non-bypassable HOG pathway variant 1055A (**Fig. 2H**), which may represent a key contact point between the proteins. Therefore, previous studies, gain- and loss-of-function alleles, and AlphaFold modeling support a trans-activation mechanism for the HOG pathway.

Trans activation may be sufficient for the HOG pathway; which does not require a properly folded SEA domain or the cytosolic tail. However, activation of the fMAPK pathway requires a properly folded SEA domain and cytosolic tail. Activation of the fMAPK pathway presumably occurs by a cis mechanism by conformational coupling between the SEA domain and tail region mediated by the transmembrane domain (**Fig. 4A**). To define contributions of the SEA domain to cis and trans signaling, a version of Msb2 lacking the NRR (Δ100-818) and cytosolic tail (stopTM) was compared to a version lacking the NRR alone. While Msb2^Δ100-818^-stopTM was functional in the HOG pathway (*Fig. S17*), it transmitted ∼10% of the signal conveyed by Msb2^Δ100-818^ in the fMAPK pathway based on growth reporter activity (**Fig. 4F**) and P∼Kss1 levels (**Fig. 4G**). These cells also underwent filamentous growth to a lesser degree (*Fig. S7*). Therefore, trans signaling is sufficient for HOG pathway activation, whereas cis signaling predominates fMAPK pathway activation with trans signaling playing a minor role.

### Cleavage by yapsins make specific cis and trans contributions to mucin regulation

The proteolytic processing of Msb2 by yapsins is necessary for activation of the fMAPK pathway (Vadaie *et al*. 2008) and may represent a point of specificity between MAPK pathways. SEA domains include processing sites (Levitin *et al*. 2005; Akhavan *et al*. 2008; Brogan *et al*. 2023), and potential cleavage sites were identified in the SEA domain of Msb2 (*Fig. S10C*, pink). However, the sites were not needed for processing, which may occur elsewhere in the protein (*Fig. S23*). As a result of processing, the extracellular domains of Msb2 and Hkr1 are shed from cells (Pitoniak *et al*. 2009). Msb2 shedding requires yapsin aspartyl proteases (Vadaie *et al*. 2008), but whether yapsins are required for Hkr1 shedding has not been explored. Colony IB showed that the shedding of HA-Hkr1 in cells lacking the five redundant yapsin proteases [*5yps*Δ (Krysan *et al*. 2005)] was reduced to the same degree as for HA-Msb2 (**Fig. 5A**). The HA-Hkr1 protein also accumulated in cell pellets in the 5*yps*Δ by IB analysis (**Fig. 5B**, *Fig. S21B*). Supernatant fractions showed low levels of HA-Hkr1, perhaps due to impaired migration of the extracellular domain of Hkr1, which is larger than the extracellular domain of Msb2. Therefore, yapsin proteases are required for shedding of Msb2 and Hkr1.

**Figure 5.**
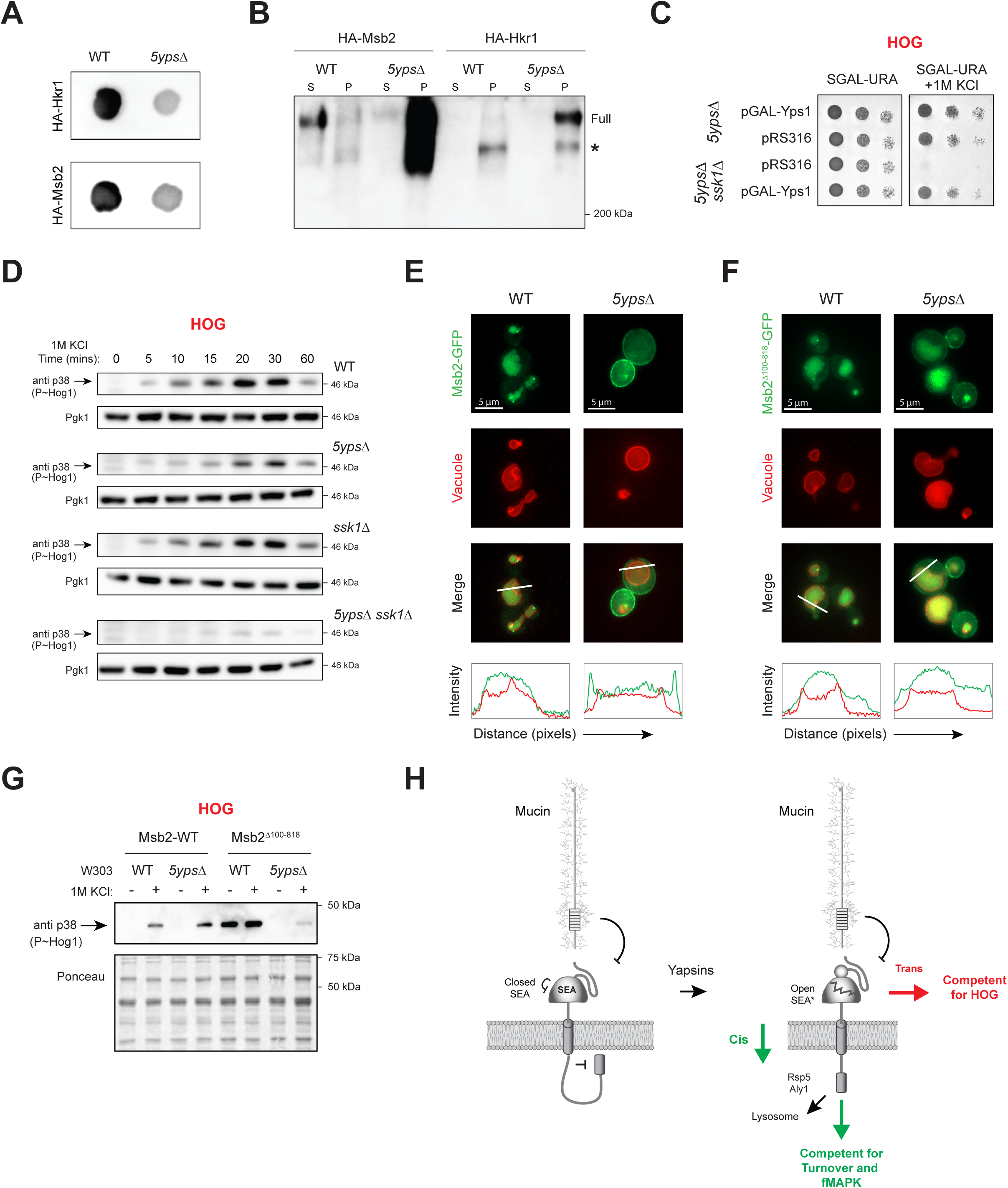
Role of yapsins in mucin signaling. **(A)** ColonyIB of wild-type cells (PC2212) and the *5yps*Δ mutant (PC2683) containing pHA-Hkr1 (PC8248) or pHA-MSB2 (PC1456). Cells were spotted onto nitrocellulose membranes atop SD-URA media for 36h. Filters were washed to remove cells and probed with anti-HA antibodies. **(B)** Supernatant (S) and Pellet (P) fractions from WT cells and the *5ypsΔ* mutant cell extracts containing HA-Msb2 or HA-Hkr1 revealed by anti-HA antibodies. asterisk, underglycosylated product. The MWM at 200 kDa is marked. **(C)** Salt sensitivity of the *5ypsΔ* (PC2683) and *5ypsΔ ssk1Δ* (PC8373) strains containing pRS316 (empty vector) or pGAL-Yps1(PC3507). Cells were spotted onto S-GAL-URA or S-GAL-URA+1M KCl media. **(D)** Phosphoblot analysis of P∼Hog1 levels in wild-type cells (WT, PC2212) and the *5ypsΔ* (PC2683), *ssk1Δ* (PC6580), and *5ypsΔssk1Δ* (PC8373) strains treated with 1M KCl for the indicated time points. Blots were probed with anti-phospho-antibodies and antibodies to Pgk1 as a control for protein loading. Background band is marked with an asterisk. **(E)** Co-localization of pMsb2-GFP in WT cells and the *5ypsΔ* mutant harboring vacuolar marker Vph1-mCherry. Bar, 5 microns. **(F)** Colocalization of pMsb2^Δ100-818^-GFP in WT cells and the *5ypsΔ* mutant harboring vacuolar marker Vph1-mCherry. Line plots are shown as described in Fig. 2. Bar, 5 microns. **(G)** Phosphoblot analysis of P∼Hog1 levels to show activation by Msb2-WT, Msb2^Δ100-818^ in WT and *5ypsΔ* under basal and inducing conditions. See panel (D) for details. **(H)** Model of Msb2 function in the fMAPK and HOG pathways. In the off state, a series of auto-inhibitory mechanisms prevent signaling. Cleavage potentiates the SEA domain for activation in cis- and trans- modes. Upon stimulus sensing, the extracellular domain is released, which leads to a conformational change resulting in trans-activation of the HOG pathway and cis activation of fMAPK pathway. To a lesser degree, both pathways utilize both cis- and trans- modes of signaling. Blue star, hyperactive allele W961C identified in the study. Processing and release of the extracellular domain also results in mucin turnover from the plasma membrane (PM).

Yapsins are required to activate the fMAPK pathway (Vadaie *et al*. 2008). To determine whether yapsins are also required to activate the HOG pathway, cells lacking yapsins (*5ypsΔ*) and the redundant branch of the HOG pathway (*ssk1*Δ) were tested for salt sensitivity. Compared to the *5yps*Δ strain, the *5ypsΔ ssk1Δ* mutant was defective for growth on 1M KCl (**Fig. 5C**). The growth defect was complemented by a plasmid containing Yps1 (**Fig. 5C**), the major yapsin that processes Msb2 (Vadaie *et al*. 2008). Phospho-IB analysis showed that the *5ypsΔ ssk1Δ* strain had reduced levels of P∼Hog1 compared to wild-type cells, the *5yps*Δ mutant, and the *ssk1*Δ mutant exposed to 1M KCl (**Fig. 5D**). Therefore, yapsins are required for activation of the HOG pathway, which is presumably required for trans interaction with tetraspanin.

Yapsin processing may impact other aspects of Msb2 function, including mucin localization. Compared to wild-type cells, which show Msb2-GFP localization in the vacuole due to turnover, Msb2-GFP was found at the plasma membrane in the *5ypsΔ* mutant (**Fig. 5E**). Therefore, yapsin processing is required for turnover of Msb2 from the plasma membrane. Hkr1-GFP was not detected due to low levels in the cell. Yapsin processing of Msb2 may lead to conformational changes in the cytosolic tail that allow recognition by Rsp5 (**Fig. 5F**), the ubiquitin ligase that regulates turnover of Msb2 (Adhikari *et al*. 2015b) and other plasma membrane proteins (Katzmann *et al*. 2004). Because the cytosolic tail of Msb2 is critical for the fMAPK pathway, the yapsins may impact specificity due to cis-based conformational changes that lead to fMAPK turnover. Although Msb2 is enriched at the plasma membrane in the *5yps*Δ mutant, it did not activate the HOG pathway (**Fig. 5D**), which supports the idea that yapsins are also required for trans signaling of Msb2 in the HOG pathway. Therefore, processing of the SEA domain is required for cis and trans activation of Msb2.

Loss of the NRR of Msb2 is required for activation of the protein and may impact yapsin-dependent signaling and turnover. The version of Msb2 that lacks NRR (Msb2 ^Δ100-818^-GFP), bypassed the internalization defect of the *5ypsΔ* mutant (**Fig. 5F**). This result supports data that release of the inhibitory region promotes cis signaling (see **Fig. 4G**). By comparison, activation of the HOG pathway by Msb2^Δ100-818^ was dependent on the yapsins (**Fig. 5G**). which indicates that release of the inhibitory domain cannot bypass trans signaling (**Fig. 5H**). These results highlight the mechanistic differences for how Msb2 functions in the two MAPK pathways that it regulates.

## DISCUSSION

Protein folding is critical for the function of most proteins. Protein misfolding disrupts function and is a major cause of protein aggregation and disease. Here, we describe an example where misfolding of a domain in an environmental sensor leads to differential signaling of one MAPK pathway over another. This mechanism indicates that protein misfolding is not inherently detrimental but may encode information about the external environment. Proteins in their misfolded states are recognized by quality control pathways, like the unfolded protein response (UPR) and ERAD, and by chaperones that promote protein re-folding or degradation (Cox *et al*. 1993; Mori *et al*. 1993; Causton *et al*. 2001; Lindholm *et al*. 2017; Abdel-Nour *et al*. 2019; Lam *et al*. 2020; Sitarik *et al*. 2025). The mechanism proposed here differs from quality control mechanisms because misfolding occurs on outside surface to provide cells with information about the environment. Here, misfolding is tolerated and may serve as a signal to a stress-sensing pathways about environments that could readily cause protein damage.

### SEA domains are conserved signaling modules in fungal species

SEA domains are present in a diverse family of glycosylated sensor proteins including mucins (Levitin *et al*. 2005), Notch (Pei and Grishin 2017), Dystroglycan (Akhavan *et al*. 2008), Enterokinase (Bork and Patthy 1995), Agrin (Patel *et al*. 2012), and cadherins (Pei and Grishin 2017). SEA domains have been identified in mammals, invertebrates [like *Drosophila melanogaster* and *Caenorhabditis elegans* (Pei and Grishin 2017)], protists, and even bacteria (Brogan *et al*. 2023; Chen *et al*. 2023). Structural predictions identified SEA domains in the mucins of yeast and other fungal species. The discovery of SEA domains in fungi fills an important gap in their distribution across species and may provide insights into their evolutionary origins and functional roles. SEA domains may have been missed in fungi due to their low sequence conservation. By searching for SEA domains with structure-based search algorithms, new mucin homologs in distantly related plant and animal pathogens were uncovered. In yeast, SEA domains regulate MAPK pathways that promote filamentous growth and osmotic tolerance, highlighting their importance in signal transduction. The identification of a structured domain in a sensory glycoprotein represents an important step towards understanding how signal sensing occurs, which is of particular interest for sensors that detect and integrate multiple stimuli. In fungal pathogens, SEA domains on the outside surface may represent potential targets for therapeutics.

### SEA domain misfolding allows discrimination between MAPK pathways

The yeast mucin Msb2 regulates different MAPK pathways, and the SEA domain plays a critical role in MAPK pathway specificity. Misfolded SEA domains did not function in a differentiation pathway (fMAPK) but functioned in a stress-response pathway (HOG), highlighting an unexpected tolerance by the HOG pathway for misfolding in the SEA domain. SEA domain stability was assessed by domain mapping, which identified residues buried in the structure that form hydrogen bonds needed to hold the domain together. To more comprehensively explore the folding landscape of the SEA domain, we developed a SDQ score by comparing protein levels/trafficking to the ΔΔG scores at each position in the domain. Misfolded SEA domains had high SDQ scores, were mis-localized in the endomembrane system, and were turned over by a quality control pathway (ERAD) that recognizes misfolded proteins. The fact that misfolded SEA domains are turned over by ERAD is itself an important finding. In humans, mutations in SEA domains are commonly associated with diseases such as cancer and muscular dystrophy (Gariballa *et al*. 2024), and pharmacological inhibition of ERAD may provide a strategy to restore glycoprotein function in clinical settings.

The differential tolerance by pathways for protein misfolding provides a mechanistic framework to explain how a single sensor can produce pathway-specific outcomes. Misfolding of the SEA domain is tolerated by the HOG pathway but prevents activation of a competing fMAPK pathway. As a result, the fMAPK cannot be activated under conditions unfavorable for filamentous growth, resulting in activation of a stress response pathway needed for survival. Protein misfolding may be the underlying cause for altered specificity, rather than a specific conformational change. This conclusion is supported by the fact that multiple variants were identified in distinct regions of the domain, which show broad variation in protein levels, including loss of attachment of the domain with the plasma membrane and dispensability of the cytosolic tail. Protein fusion experiments further support the idea that the stability of the SEA domain, not a specific aa sequence motif, is critical for specificity. The extraordinary tolerance of the HOG pathway to SEA domain deformation suggests the pathway may have developed tolerance to mucin deformation over evolutionary time due to chronic exposure of the SEA domain to unfavorable environments. Misfolded domains on the outside surface may be refolded by secreted chaperones or subject to irreparable damage. The rapid turnover of Msb2 may replace misfolded proteins by synthesis of new versions of the protein to re-balance sensory functions at the plasma membrane. The ability to tolerate misfolding of external domains may be common in microbes, which must constantly adapt to fluctuating environments. However, most organisms need to detect external changes, including mammals. Human MUC1 for example also resides on the outside surface of mucosal cavities and is directly exposed to the environment (Dhar and McAuley 2019; Chen *et al*. 2021). Therefore, regulatory pathways may detect proteins in their misfolded states to interpret information and respond to stress.

### Cis and Trans Signaling Occur Differently for Different MAPK Pathways

The different functional properties of mucin revealed different modes of MAPK pathway activation. One mode is dominated by trans-activation of the tetraspanin complex by the SEA domain to activate the HOG pathway. The trans mode of activation can be studied in isolation because of the dispensability of the mucin cytosolic tail. Since SEA domain misfolding is tolerated in HOG, we conclude that a significant portion of the SEA domain structure is dispensable for trans signaling. Moreover, misfolding of the SEA domain by extracellular stress may lead to conformational rearrangements and serve as part of the sensory mechanism. We present a model where this activation event is depicted as removal of a regulatory arm through environmental triggers or genetic perturbations leading to misfolding that reveals an altered interaction interface between the SEA and the CR domain of tetraspanin. Such changes presumably result in conformational changes in the cytosolic domains of tetraspanin, leading to recruitment of cytosolic proteins, like activator of the Rho GTPase Cdc42 (Gonzalez and Cullen 2022) and adaptor for the MAPKKK Ste11 to the plasma membrane (Truckses *et al*. 2006). The recruitment of these proteins activates the pathway (Yamamoto *et al*. 2010). The trans activation mechanism may be transient, which fits with the kinetics of the HOG pathway as being required for short-term adaptation to osmotic and other stresses. Although the cytosolic tail is dispensable in the HOG pathway, it may control aspects of HOG pathway signaling in certain settings (Tatebayashi *et al*. 2007).

By comparison, the fMAPK pathway requires a properly folded SEA domain for cis activation through conformational changes that impact the mucin cytosolic tail. The fMAPK pathway also requires some degree of trans signaling, which may amplify signaling to achieve sustained activation that is required for differentiation to the filamentous cell type. Although cleavage by yapsins, and release of the inhibitory domain, are prerequisites for activation of both pathways, cleavage is also required for mucin turnover, which specifically impacts cis signaling of the fMAPK pathway. The cytosolic tail of Msb2 may recruit pathway-specific adaptors (Pitoniak *et al*. 2015) and kinases (Madhani *et al*. 1997) to promote a pathway-specific response.

### Protein misfolding as a sensory mechanism

Mechanosensors function in different ways, from touch sensation by ion channels (Coste *et al*. 2010), to detection of aneuploidy by the environmental stress response (Coste *et al*. 2010; Kane *et al*. 2021; Omkar *et al*. 2025), to sensing thermal signals to generate a global response (Omkar *et al*. 2025). The HOG pathway in yeast is activated by chemically unrelated stimuli that share the common feature of compromising protein folding. We suggest that SEA domain misfolding may occur in response to harsh environmental conditions needed to activate the HOG pathway. In support of the possibility, stimuli that induce protein misfolding require Msb2 and the HOG pathway for survival.

In conclusion, properly folded domains represent the ideal functional state for most proteins. However, structural deformation of protein domains may not always be detrimental. Regulatory pathways may tolerate protein misfolding, particularly when folding is compromised upon exposure to harsh environments. Sensory pathways required under stress conditions may detect protein misfolding to gain information about the environment or other proteostatic stresses. The tolerance and recognition of protein misfolding by regulatory pathways may represent a general way to respond to stress.

## MATERIALS AND METHODS

### Strains and Plasmids

Yeast strains were grown and manipulated using published protocols (Rose *et al*. 1990). Yeast strains are described in Table S1. Plasmids are described in Table S2, and primers are listed in Table S3. Yeast gene deletions were made by homologous recombination with auxotrophic and antibiotic resistance markers amplified by polymerase chain reaction (PCR) using published templates (Goldstein and McCusker 1999). The *msb2Δ hkr1Δ ssk1Δ* triple mutant (PC8102) was made using homologous recombination of *msb2::HYG* cassette at the *MSB2* genomic locus in the *hkr1::KanMX6 ssk1::NAT* (PC7883) strain. The *hkr1::KanMX6 ssk1::NAT* was made from the *ssk1::NAT* strain (PC6580), which has been described (Pitoniak *et al*. 2009).

Plasmids were constructed by gap repair and Gibson Assembly [(Gibson *et al*. 2009), Gibson Assembly Master Mix, New England Biolabs, Beverly MA, Catalog # E5510S]. The plasmid containing the pHA-MSB2 gene (PC1456) has been described (Vadaie *et al*. 2008). The plasmid contains an internal epitope-tagged version of hemagglutinin (HA) at position 500 aa residues driven by the endogenous promoter. The plasmids pHA-MSB2^Δ100-818^ (PC7936) and pHA-MSB2^Δ100-818^-GFP (PC8169) were made by gap repair from strains containing Msb2^Δ100-818^ (PC1516) and Msb2^Δ100-818^-GFP (PC2586) using pHA-MSB2 digested with the HpaI enzyme.

p*GAL-YPS1* (PC3507) has been described (Krysan *et al*. 2005). To facilitate site-directed mutagenesis, pHA-MSB2 was cloned into pRS316 by Gibson Assembly. The HA-Msb2 gene and 1 kb of the promoter region were amplified from pHA-MSB2 and assembled into pRS316 to obtain the pRS316-HA-Msb2 (PC8338). Specifically, the Msb2 open reading frame (ORF) along with 1kb promoter region was divided into three parts ranging from ∼1.5-2 kb, and overlapping primers were designed for the PCR amplification. The amplified cassettes were assembled in a single reaction. The pHA-Hkr1 plasmid (PC8248) was also made by Gibson Assembly using primers to amplify the HA-Hkr1 gene and 1kb of upstream promoter region from strain HA-Hkr1 [PC2740, (Pitoniak *et al*. 2009)]. The PCR product was assembled into pRS316 to obtain the full length pHA-Hkr1.

The chimeras were made using Gibson Assembly in which overlapping primers were designed to contain portion of *MSB2* and *HKR1* at the chimera junctions. The positions of the primers were chosen based on the occurrence of the SEA domain in each protein. Three regions were amplified from pHA-MSB2 and one from a region containing the Hkr1 SEA from pHA-HKR1 were assembled into the pRS316 plasmid at SalI and BamHI sites to make pHA-MSB2^HKR1-SEA^ (PC8463). The resulting ORF was driven by the *MSB2* promoter. Similarly, two amplified regions from pHA-HKR1 and pHA-MSB2 each were assembled into the pRS316 plasmid at XhoI and BamHI sites to make pHA-HKR1^N^MSB2^C^ (PC8472) that was driven by the *HKR1* promoter.

### Site-directed Mutagenesis

To generate point mutations in the SEA domains and flanking regions of *MSB2* and *HKR1*, a GeneArt site directed mutagenesis kit (Life Technologies, Thermo Fisher, Cat #A13282) was used according to the manufacturer’s instructions. Complementary primer sets (∼40nt long, *Table S3*) carrying the nucleotide changes at the center of the primers were designed. The primers were used to amplify mucin genes from templates (pHA-MSB2 or pRS316-HA-MSB2 and pHA-HKR1) using the AccuPrime Pfx DNA polymerase (Thermo Fisher Cat #12344024) in a reaction containing DNA methylase and S-adenosyl methionine to methylate the template plasmid. The PCR product was transformed into *E. coli* cells (One Shot MAX Efficiency DH5α-T1^R^) that select for the newly synthesized unmethylated product by digesting the methylated parent plasmid with a specially designed *McrBc* endonuclease (Panne *et al*. 2001). The mutagenized plasmids were isolated and analyzed by whole plasmid sequencing by Plasmidsaurus (https://plasmidsaurus.com/). As a byproduct of the mutagenesis, short stretches of repeated insertions were generated in some cases (10-12 aa stretches ranging from 1-30 repeats, called Msb2^RPTS^) that were also included in functional tests. Some out-of-frame truncations were also generated that resulted in a truncated protein with the designation (TR, truncated).

### Bioinformatics analysis

The Uniprot protein database (https://www.uniprot.org/) was used to obtain information about proteins and protein domains. The protein structure prediction program AlphaFold (Jumper *et al*. 2021; Varadi *et al*. 2024) along with the version AF2 was used for structure predictions. AlphaFold protein database (AFDB) (https://alphafold.ebi.ac.uk/) was used to extract the predicted structures of the top hits from structural homology searches for additional analysis. PROSITE (https://prosite.expasy.org/) was used to identify reported SEA domains in other glycoproteins (Sigrist *et al*. 2013). SEA domains from the study will be included into the entry heading SEA domain (Accession ID- PS50024).

The protein-protein BLAST search algorithm (https://blast.ncbi.nlm.nih.gov/Blast.cgi) was used to identify homologous proteins in fungal species (Altschul *et al*. 1990). The sequence of the Msb2 SEA domain (1000-1110) was used to find similar sequences from the ClusteredNR (non-redundant) database of proteins. Multiple sequence alignment (MSA) was performed in ClustalW (https://www.genome.jp/tools-bin/clustalw) or ClustalΩ (http://www.clustal.org/omega/) using default parameters to annotate the conserved residues across the fungal SEA domains. The Yeast Gene Order Browser (http://ygob.ucd.ie/) was used to compare the *MSB2* and *HKR1* genes at their respective genomic loci to reveal gene synteny (Byrne and Wolfe 2005). Foldseek (https://search.foldseek.com/search) was used to perform proteome-wide structural homology searches (Van Kempen *et al*. 2024). The .pdb file of Msb2 SEA domain (1000-1110aa) was used for this analysis. Default parameters were used for this search. As indicated, the Foldseek plugin (under structural analysis) provided by ChimeraX was employed for searches. Hits that show an E-value > 10^-5^ were considered for analysis. The NCBI taxonomic database (https://www.ncbi.nlm.nih.gov/taxonomy) (Schoch *et al*. 2020) was used to analyze the list of Foldseek hits and a representative uncharacterized protein was analyzed for each group. A SEA domain was not found in the fission yeast *Schizosaccharomyces pombe*, which may have lost its Msb2 protein, as homologs were identified in closely related species.

The structures of SEA domains from different proteins were extracted as .pdb files from the AlphaFold prediction models. All protein structure visualizations were performed on a local computer application ChimeraX (Meng *et al*. 2023) or PyMol (The PyMOL Molecular Graphics System, Version 1.2r3pre, Schrödinger, LLC). Structural alignments were performed-using TM-align (https://zhanggroup.org/TM-align/) (Zhang and Skolnick 2005). The TM-score for each SEA domain was obtained by comparing the structure to the Msb2 SEA domain. The score was normalized to the length of reference protein (Msb2-SEA). TM-align also provided a root mean square deviation (RMSD) value indicating overlap.

In silico saturating mutagenesis (FOLDX, MUTATEX) (Schymkowitz *et al*. 2005; Tiberti *et al*. 2022) was performed using the high-performance computing center CCR at University at Buffalo. Structural analysis including hydrogen-bond analysis were performed with ChimeraX (Meng *et al*. 2023). The Structural Deformation Quotient (SDQ) score was calculated as weighted sum of the normalized factors from experimental and predicted scores. A greater weight was assigned to experimental component than predicted structural components because it captures a direct phenotypic/functional consequence of the variant.

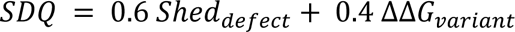

where 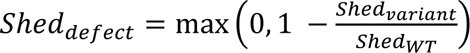, which increases when shedding is reduced as compared to wildtype. The structural term ΔΔ*G_variant_* was further defined as a weighted combination of the normalized mutation-specific and position-specific FoldX destabilization terms. Higher SDQ scores indicate high degree of misfolding through reduced shedding and high structural destabilization. Other Bioinformatics tools were employed to obtain domain information in the newly discovered potential Msb2 homologs. DeepTMHMM (https://dtu.biolib.com/DeepTMHMM) /TMHMM-2.0 (Krogh *et al*. 2001) (https://services.healthtech.dtu.dk/services/TMHMM-2.0/) was used to identify transmembrane domains. TargetP2.0 (https://services.healthtech.dtu.dk/services/TargetP-2.0/) (Almagro Armenteros *et al*. 2019) was used to identify N-terminal signal peptide sequences. T-REKS (https://bioinfo.crbm.cnrs.fr/index.php?route=tools&tool=3) (Jorda and Kajava 2009) was used to find repeated sequences. Unless otherwise indicated, SEA domain-containing proteins had a well-defined N-terminal signal sequence [TargetP-2.0 (Almagro Armenteros *et al*. 2019)] and a S/T/P– rich extracellular domain with tandem repeats [T-REKS, (Jorda and Kajava 2009)] with the SEA domain being adjacent to the transmembrane α-helix [TMHMM-2.0, (Krogh *et al*. 2001)].

### Growth Reporter, Salt, and Filamentation Assays for MAPK pathway activity

The *FUS1-HIS3* reporter is dependent on the transcription factor Ste12 (McCaffrey *et al*. 1987) and provides a readout of the mating and fMAPK pathways. In cells lacking an intact mating pathway (*ste4 FUS1-HIS3*), shows dependency on Msb2 (Cullen *et al*. 2004). A strain lacking Msb2 (*ste4Δ msb2Δ*, PC999) was transformed with plasmids expressing SEA domain variants, a wild-type version of Msb2 (pHA-MSB2) as a positive control, and an empty vector (pRS316) as a negative control. Serial dilutions of cells were spotted onto media lacking uracil (-URA) to maintain selection for the plasmid and media lacking histidine (-HIS) to assess fMAPK reporter activity. Cells (*ste4*Δ *msb2Δ hkr1Δ ssk1Δ*, PC8102) harboring Msb2-SEA variants were grown for 16h in selective media. Cells were diluted and adjusted to an optical density (OD_600_) of 0.1. Three serial dilutions were spotted onto SD-URA media as a control for growth and SD-URA-HIS media to assess reporter activity. For some experiments, cells were spotted onto SD-URA-HIS+ 3-Amino-1,2,4-triazole (ATA) to assess elevated reporter activity. Cells were spotted onto SD-URA+1M KCl to test for salt sensitivity as a readout of the HOG pathway (Tatebayashi *et al*. 2007). The single cell invasive growth assay was performed as described (Cullen and Sprague 2000).

Cells (*ste4Δ msb2Δ hkr1Δ ssk1Δ*; PC8102) harboring pHA-MSB2 or the empty vector pRS316 were grown for 48 h in selective media containing the indicated stressors (e.g., salt, urea, DTT) and compared with cells lacking an intact HOG pathway (*pbs2*Δ). Overnight cultures were diluted to an OD_600_ of 0.05 and grown in 96-well plates, with growth monitored every 30 min by measuring OD_600_ using a TECAN Spark® microplate reader. Absorbance at saturation (20h) was used to determine growth differences.

### Detection of shed mucins from yeast

Colony IB analysis was performed as described (Chavel *et al*. 2010). Cells containing plasmids harboring internal and functional epitope-tagged mucins were patched on a nitrocellulose filter atop semi-solid agar SD-URA media to maintain selection of the plasmids. Cells were incubated for 24h, and patches were washed off the filter to detect levels of shed protein. Immunoblot analysis was performed on filters using anti-HA antibodies (#12CA5). SpotIB analysis was performed as described (Vadaie *et al*. 2008). Supernatants derived from strains harboring epitope-tagged mucins grown for 16h in SD-URA media were diluted and normalized based on the OD_600_ of the cells and equal volumes spotted onto a nitrocellulose membrane. Membranes were probed with anti-HA antibodies to detect levels of shed mucins.

### Immunoblot Analysis

To detect HA-Msb2 and HA-Hkr1 proteins, proteins derived from yeast cell extracts were resolved by sodium dodecyl sulfate polyacrylamide gel electrophoresis (SDS-PAGE) with 6% acrylamide as described (Vadaie *et al*. 2008). Proteins were transferred to nitrocellulose membranes (Amersham Protran Premium 0.45 μm NC, GE Healthcare Life Sciences). Following the transfer, membranes were incubated with PonceauS staining solution (Thermo Scientific™, Catalog number A40000279) and were used as a proxy for loading controls. Membranes were incubated with blocking buffer 5% nonfat dry milk in TBST [10 mM Tris-HCl pH 8, 150 mM NaCl, and 0.05% Tween 20]. Antibodies were diluted with blocking buffer. Anti-HA antibodies (Roche Diagnostics, 12CA5) were used at 1:5000 dilution to detect proteins in supernatant and pellet. GFP-tagged proteins were detected with anti-GFP antibodies at 1:5000 dilution (Roche Diagnostics, clones 7.1 and 13.1, #11814460001). For secondary antibodies, HRP conjugated goat anti-mouse antibodies (Jackson ImmunoResearch Laboratories, #111-035-144) were used at 1:5000 dilution. Blots were washed three times for 5 min each in TBST in between primary and secondary antibody incubations. Membranes were incubated with primary antibodies for 16h at 4°C and with secondary antibodies for 1h at 22°C.

### Phospho-immunoblot Analysis

The detection of phosphorylated MAP kinases was performed as described (Karunanithi and Cullen 2012). Cells were grown to mid-log phase (3-4h) and shifted to inducing conditions for time points indicated. Cells were shifted to YEPGal (2% Galactose) media for 6h to induce fMAPK pathway and YEPD+1M KCl for up to 1h to induce HOG pathway activity. Cells were harvested and stored as cell pellets at -80°C. Cells were lysed by mechanical lysis using glass beads and vortexed at max for 15 mins with alternate 1 min pulse and 1 min rest on ice. TCA lysis buffer [10% trichloroacetic acid (TCA), 10mM Tris-HCl pH 8, 25mM Ammonium acetate, 1mM EDTA) was used to precipitate proteins. Protein extracts were run on 10% SDS-PAGE gels. ERK-type MAPKs (P∼Kss1, P∼Fus3, and P∼Slt2 as indicated) were detected using anti-p44/42 (Cell Signaling Technology, #4370, 1:10,000 dilution) antibodies. Anti-phospho-p38 MAPK antibodies (Cell Signaling Technology, #9211, 1:10,000) were used to detect P∼Hog1. Phospho-MAPK antibodies were diluted in 5% BSA in 1X TBST [10 mM TRIS-HCl (pH 8), 150 mM NaCl, 0.05% Tween 20]. Membranes were incubated with primary for 16h at 4°C and with secondary for 1hr at 22°C. For secondary antibodies, HRP conjugated goat anti-rabbit (Jackson ImmunoResearch, #111-035-144, 1:2000) diluted in 5% skim milk was used. The blot was imaged by a ChemiDoc MP Imaging System (BioRad, #12003154) and signal intensity was measured by using the Lane and Bands tool in the Image Lab Software.

### Fluorescence microscopy

For co-localization experiments with the vacuole, the vacuolar marker Vph1-mCherry [ZJOM65 (Plasmid #133654)] was introduced into strain PC2613 by digestion of plasmid PC8050 with the restriction enzyme PmlI to generate strain PC8205. For co-localization experiments with the endoplasmic reticulum, the marker Elo3-mCherry [ZJOM149 (Plasmid #133646)] digested with restriction enzyme BstBI was introduced into strain PC2613 to generate strain PC8206. These plasmids were obtained from Addgene (https://www.addgene.org/browse/article/28206918/) and have been described in (Zhu *et al*. 2019). Cells containing pMSB2-GFP in vacuolar-marked strains (PC8205) or endoplasmic reticulum marked strains (PC8206) were grown for 16h in selective minimal media (SD-URA-TRP) at 30°C, subcultured by dilution into fresh medium, and grown until mid-log phase 5h in minimal (SD-URA-TRP) media before imaging.

Brightfield and Fluorescence images were taken using a ZEISS Axioplan 2 fluorescent microscope equipped with a Plan-Apochromat 100×/1.4 (oil) objective (Carl Zeiss Microscopy, LLC). The ZEISS ZEN Microscopy Software (RRID:SCR_013672) was used to analyze the microscopy images. ImageJ was used adjust brightness and contrast of Brightfield and Fluorescence images. The profile plot setting of FIJI/ ImageJ (Schindelin *et al*. 2012) (RRID:SCR_002285) was used to measure protein localization across the length of the cell. The intensity of individual plots was adjusted to match scale.

## ABBREVIATIONS

aa: amino acid
AFDB: AlphaFold Protein Database
ATA: 3-Amino-1,2,4-triazole
CR: cysteine rich
ER: endoplasmic reticulum
GPCR: G-protein coupled receptor
GOF: gain-of-function
HA: hemagglutinin
GXE: Gene by Environment
HOG: high osmolarity glycerol response pathway
IB: immunoblot
kb: kilobase
kDa: kilodalton
K: kinase
LOF: loss-of-function
MAPK: mitogen-activated protein kinase pathway
MHD: Mucin Homology Domain
NRR: negative regulatory region
OD: optical density
ORF: open reading frame
P: pellet
PM: plasma membrane
PRR: positive regulatory region
RPTS: repeat
SEA: sea urchin sperm protein, enterokinase, agrin domain
SpotIB: spot immunoblot analysis
S: supernatant
SDQ: structural deformation quotient
SNV: single nucleotide variant
TCA: trichloroacetic acid
TM: transmembrane
TR: truncated
VM: vacuolar membrane; and
WT: wild type.

## ACKNOWLEDGMENTS

Thanks to J. Brodsky (Penn State University) for providing reagents. Thanks to Gunnar Hansson (University of Götenberg, Sweden), Henrik Dohlman (University of North Carolina), and Laura Rusche (University at Buffalo) for reading the manuscript and providing comments. Thanks to Aditi Prabhakar, Matthew Vandermeulen, and other laboratory members for suggestions. Thanks to Center for Computational Research (CCR) at University at Buffalo for high-performance computing resources.

