## Supplemental Materials for "Differential tolerance for SEA domain misfolding encodes a MAPK pathway-specific response"

#### SUPPLEMENTAL FIGURE LEGENDS

***Fig. S1.*** **Diagram of the SEA domain of Msb2 compared to the SEA domains in other proteins.** Blue, N terminus; Red, C terminus. (A) Msb2, (B) Agrin, (C) Enterokinase, (D) MUC1, (E) IMPG1, (F) Notch, (G) Dystroglycan, (H) PTPRN, (I) CD34, (J) Collectrin, and (K) MUC13. Structures were analyzed using ChimeraX.

***Fig. S2.*** **Structural overlap of the SEA domain of Msb2 (1000-1110) with the SEA domains in human proteins.** Proteins tested were (A) Enterokinase (AFDB# AF-P98073-F1), (B) Agrin (AFDB# AF-O00468-F1), (C) Notch (AFDB# AF-P46531-F1), (D-E) Dystroglycan (AFDB# AF-Q14118-F1) SEA domains I and II, (F) CD34 (AFDB# AF-P28906-F1), (G) Collectrin (AFDB# AF-Q9HBJ8-F1), (H) PTPRN (AFDB# AF-Q16849-F1) and (I) MUC1 (AFDB# P15941). Structures were visualized and aligned on ChimeraX. Structural similarity score (TM-scores) was calculated using TM-align (ZHANG AND SKOLNICK 2005).

***Fig. S3.*** **Alignment of hydrophobic residues (highlighted in yellow) in fungal and the indicated mammalian SEA domains.** Secondary structure predictions were marked by an "e" for ß-strands and "h" for α-helices. Alignments were based on (PEI AND GRISHIN 2017) and some sequences were aligned by eye. The Uniprot IDs/AFDB# for each protein are shown. The %Id denotes amino acid sequence identity with the SEA domain of Msb2. The term SEA denotes the length of the SEA domain based on the number of aa residues. Species designations are *H.sa*, Homo sapiens; *H.pi*, *Hydnomerulius pinastri; P.gr, Puccinia graminis; U.ma, Ustilago maydis; T.de, Taphrina deformans; H.ca, Histoplasma capsulatum; E.we, Escovopsis weberi; L.ma, Lophiostoma macrostomum; C. au, Candidozyma auris; C.al, Candida albicans; E.go. Eremothecium gossypii; K.la. Kluyveromyces lactis; C.gl. Candida glabrata;* and *S.ce, Saccharomyces cerevisiae.* Red asterisks, residues associated with human diseases including limb-girdle muscular dystrophy (HARA *et al.* 2011) and muscle-eye-brain disease (GEIS *et al.* 2013) in DAG1, vitelliform macular dystrophy (MANES *et al.* 2013) and retinitis pigmentosa (BANDAH-ROZENFELD *et al.* 2010; OLIVIER *et al.* 2021; MITCHELL *et al.* 2022) in IMPG1 and IMPG2, and congenital myasthenic syndrome in Agrin (PEI AND GRISHIN 2017).

***Fig. S4.*** **Sequence homology and structure of SEA domains in fungal Msb2 homologs. (A)** Alignment of fungal SEA domains by ClustalΩ, Dark blue, conserved residues; light blue, similar residues; Red arrows, residues predicted to be critical for structure; Blue arrows, residues conserved across fungal species; Purple arrows, residues that are both conserved and critical for structure. See key for details. Pink arrows, new Msb2 homologs identified in fungal species *Hydnomerulius pinastri, Puccinia graminis, Taphrina deformans, Histoplasma capsulatum, Escovopsis weberi, Lophiostoma macrostomum, and Candidozyma auris.* The Opy2 interaction site comes from (YAMAMOTO *et al.* 2016). Sp., species. **(B)** Structural overlap of the fungal Msb2 SEA domains. Structures were determined by AlphaFold and aligned using ChimeraX. The TM-scores were obtained from TM-align. Organisms are the same as shown in panel *S4A*. See also Movie 3.

***Fig. S5.*** **Functional impact of epitope tag insertions in and around the SEA domain.** Left, diagram of the Msb2 protein. Yellow star, site of the HA epitope tag. Rectangles denote the PTS-rich region. 2D structure of the SEA domain is shown with site of Myc epitope insertions denoted by pink arrows, whose positions were confirmed by DNA sequencing analysis. Right, IB analysis showing the shed (S, supernatant) and cell associated (P, pellet) fractions probed by antibodies against the HA, at top, and Myc epitopes, bottom blots. The plus and minus signs refer to the activity of the fMAPK pathway determined by the activity of the *FUS1-HIS3* reporter.

***Fig. S6.*** **Evaluation of the function of the SEA domain of Msb2 in the fMAPK pathway by a transcriptional growth reporter.** The *FUS1-HIS3* growth reporter was examined in the *msb2*Δ (PC948) mutant containing plasmids with the indicated SEA domain variants on control (SD-URA) and reporter (SD-URA-HIS) media. White boxes were put into the figure to replace duplicates. The key describes different classes of alleles: cyt- cytoplasmic tail residue mutants; RPTS, introduction of one or more repeats/insertions; TR, truncated protein due to frameshift following the residue changed; green circle, point mutants defective for HOG; double green circle, and RPTS mutants defective for fMAPK signaling. pRS316 is an empty plasmid that used as a negative control.

***Fig. S7.*** **Evaluating the role of the SEA domain in regulation cell differentiation to filamentous growth. (A)** Single cell invasive growth assay for selected SEA domain variants that were defective for fMAPK pathway function. 20X magnification, bar, 5 microns. **(B)** Single cell assay of cells expressing a hyperactive version of Msb2, W961C or Msb2$^{\Delta 100\text{-}818}$, in combination with loss of function mutants as indicated. 20X magnification, bar, 5 microns. **(C)** Single cell assay showing the degree of filamentous growth in cells exposed to HOG pathway inducers, 1M KCl and 2M sorbitol. 20X magnification, bar, 5 microns.

***Fig. S8.*** **Evaluation of the function of the SEA domain of Msb2 in the HOG pathway.** The *msb2*Δ *hkr1*Δ *ssk1*Δ triple mutant (PC8102) containing Msb2 variants were grown on control (SD-URA) and SD-URA+1M KCl media. Salt sensitivity provides a readout of the HOG pathway. See the key and *Fig. S6* for details.

***Fig. S9.*** **Comparison of SEA domain function in three biological contexts.** Spot plates showing growth of Msb2 SEA domain variants on 0.5µg/ml TUN plates as compared to growth based on *FUS1-HIS3* reporter (fMAPK) and salt sensitivity test based on 1M KCl (HOG). All plates were incubated at 30˚C.

***Fig. S10.*** **Predicted SEA domain structure showing potential post-translational modification sites and sites of cleavage investigated in the study.** For each panel, the structure of the SEA domain was determined by AlphaFold, and residues tested for function were marked as indicated. **(A)** Potential N-linked glycosylation sites are shown and labelled (green). **(B)** Residues important for hydrogen bonding are shown (yellow) based on default parameters on ChimeraX. **(C)** Potential cleavage sites for yapsin aspartyl proteases are shown and labelled (pink).

***Fig. S11.*** **SpotIB for selected HA-Msb2 SEA domain variants.** Supernatants from cell cultures grown to the same density were spotted onto nitrocellulose filters. Shed protein levels were detected using the anti-HA antibody. Spots are labelled for each variant tested.

***Fig. S12.*** **Immunoblot of shed and cell-associated fractions of HA-Msb2 and HA-Hkr1 in selected SEA domain variants.** Cells were grown in SD-URA media to mid-log phase (OD $A_{600}$ = 0.8-1) to maintain selection of the plasmids. Supernatant (S) and Pellet (P). Proteins levels were detected using anti-HA antibody. No HA control came from cells harboring an empty plasmid, pRS316. Allele designations are for Msb2 unless otherwise indicated. The same amount of protein was loaded onto each lane based on cell number as determined by OD $A_{600}$.

***Fig. S13.*** **Role of ERAD in regulating the levels and localization of SEA domain variants. (A)** SEA domain variants were examined in wild-type cells (WT) or the *cdc48-10* mutant and tested for protein levels using anti-HA antibody. The same amount of protein was loaded onto each lane based on cell number as determined by OD $A_{600}$. **(B)** Localization of wildtype Msb2-GFP and the SEA domain variant Msb2-F1012A-GFP in WT cells and the *cdc48-10* mutant. Bar, 5 microns.

***Fig. S14.*** **Viability test for cells exposed to various ER stress conditions.** Cells containing the indicated Msb2 SEA variants were tested for growth on high temperature (37˚C), oxidative stress (1mM $H_2O_2$), the ER-Golgi protein trafficking inhibitor BrefeldinA (20 µg/ml) and reducing environment (4mM DTT). Plates were incubated for 48 h and photographed.

***Fig. S15.*** **Heat map of residues in the SEA domain of Msb2 based on structure.** The heat map was generated by MutateX. X-axis, each residue in the SEA domain. Y-axis, the predicted folding free energy change (ΔΔG) for each amino acid. Larger numbers (red) are less favorable for structure. Red arrows, residues predicted to be critical for structure; Blue arrows, residues conserved across fungal species; Purple arrows, conserved residues critical for structure. Circles show residues that impact function. See the key for details. Opy2 interaction site comes from (YAMAMOTO *et al.* 2016).

***Fig. S16.*** **Comparison of residues required for structure to residues conserved between species for the SEA domain of Msb2.** Top, Venn diagram showing number of conserved and structurally critical residues as derived from the ΔΔG scores. Bottom, representative region (1003-1022) comparing homology and heatmap of predicted ΔΔG scores. Refer to Key for details.

***Fig. S17.*** **Intragenic suppression of SEA domain alleles by activated version of Msb2. (A)** Spot plates showing fMAPK and HOG activity of SEA domain variants combined with the hyperactive allele $Msb2^{\Delta 100\text{-}818}$. Cells were spotted onto the indicated media and photographed after 48h. **(B)** Intragenic suppression analysis with cytosolic deletion variants and SEA domain variants of Msb2 (P990H, F1012A and P1113A). Cells were spotted onto the indicated media and photographed after 48h.

***Fig. S18.*** **Comparative functional analysis of Msb2 and Hkr1 SEA domains. (A)** Alignment of the SEA domains of Msb2 and Hkr1 performed ClustalΩ. **(B)** Spot plates showing redundant function of Msb2 and Hkr1 in HOG pathway activity and the role of the SEA domain of Hkr1. The *msb2*Δ *hkr1*Δ *ssk1*Δ (PC8102) triple mutant containing SEA variants of Hkr1 were spotted onto control (SD-URA) and SD-URA+1M KCl media to test for HOG pathway activity.

***Fig. S19.*** **Gene synteny analysis of Msb2 and Hkr1.** Gene synteny was performed using the yeast gene order browser showing the order of genes in the neighborhood of the Msb2 and Hkr1 genomic loci as indicated.

***Fig. S20.*** **Evaluation of invasive growth by the plate-washing assay.** The indicated strains, WT, *hkr1*Δ, *msb2*Δ, *hkr1*Δ *sho1*Δ, *ste12*Δ, *sho1*Δ, *msb2*Δ *sho1*Δ, and *msb2*Δ *hkr1*Δ *sho1*Δ were spotted on YEPD media for 3d at 30˚C. Cells were imaged before and after washing the cells off of the plates. An inverted image is also shown.

***Fig. S21.*** **Comparison of the levels of the HA-Msb2 and HA-Hkr1 proteins. (A)** Total levels of the HA-Msb2 and HA-Hkr1 proteins in the cells calculated using the sum of supernatant and pellet fractions. **(B)** Quantification of shedding levels of HA-Msb2 and HA-Hkr1 in wild-type cells (WT) and the *5yps*Δ mutant.

***Fig. S22.*** **AlphaFold model of the mucin sensor complex.** Msb2 (951-1205aa); Opy2 (1-138aa) and Sho1 (1-145aa).

***Fig. S23.*** **Immunoblot analysis of cell-associated HA-Msb2-GFP to test proteolytic processing in selected SEA domain variants.** Pellet fractions of the SEA variants were probed using anti-GFP antibodies alongside control strains.

#### SUPPLEMENTAL MOVIES

**Movie 1.** AlphaFold model of the 1306 aa Msb2 protein. Repeats 687-820 residues, globular domain (1000-1110), transmembrane (TM) domain 1184-1223 residues.

**Movie 2A-B.** Msb2 and MUC1 SEA domains.

**Movie 3.** Structural alignment of the SEA domains of Msb2 homologs in closely related fungal species (*S. cerevisiae*, yellow; *C. glabrata*, pink; *K. lactis*, blue; *A. gossypii*, green; *C. albicans*, orange).

**Movie 4.** AlphaFold model of the 1802 aa Hkr1 protein. Repeats 456-786 residues, globular domain (1249-1420), transmembrane domain (1485-1509).

**Movie 5A-B.** Space filling models showing location of residues in the Msb2 SEA domain critical for fMAPK (green) and HOG (red). Note- most green residues are buried (not exposed to the surface).

**Movie 6.** Structural alignment of the SEA domains of Msb2 and Hkr1.

**Movie 7.** contact site at W961 residue in proximity (4-8Å) with C30-C55 of Opy2.

#### SUPPLEMENTAL TABLES

Table S1. Yeast strains used in the study.

| Strain [a] | Genotype | Reference |
|---|---|---|
| PC313 | *MAT***a** *ura3-52* | (LIU *et al.* 1993) |
| PC538 | *MAT***a** *ste4 FUS1-lacZ FUS1-HIS3 ura3-52* | (CULLEN *et al.* 2004) |
| PC539 | *MAT***a** *ste4 FUS1-lacZ FUS1-HIS3 ura3-52 ste12::URA3* | (PITONIAK *et al.* 2015) |

| | | |
|---|---|---|
| PC948 | *MAT***a** *ste4 FUS1-lacZ FUS1-HIS3 ura3-52 msb2Δ::KanMX6* | (CULLEN *et al.* 2004) |
| PC998 | *MAT***a** *ste4 FUS1-lacZ FUS1-HIS3 ura3-52 MYC-MSB2* | (VADAIE *et al.* 2008) |
| PC999 | *MAT***a** *ste4 FUS1-lacZ FUS1-HIS3 ura3-52 HA-MSB2* | (VADAIE *et al.* 2008) |
| PC1558 | *MAT***a** *ste4 FUS1-lacZ FUS1-HIS3 ura3-52 sho1::HYG ssk1::NAT* | (PITONIAK *et al.* 2015) |
| PC2053 | *MAT***a** *ste4 FUS1-lacZ FUS1-HIS3 ura3-52 pbs2::KanMX6* | (PITONIAK *et al.* 2009) |
| PC2212 [b] | *MAT***a** *ade2-1 his3-11,15 leu2-3,112 ura3-1 trp1-1 can1-100* | (KRYSAN *et al.* 2005) |
| PC2613 | *MAT***a** *ste4 FUS1-lacZ FUS1-HIS3 ura3-52 trp1::NAT* | This study |
| PC2625 | *MAT***a** *ste4 FUS1-lacZ FUS1-HIS3 ura3-52 hkr1::HYG* | (PITONIAK *et al.* 2009) |
| PC2662 | *MAT***a** *ste4 FUS1-lacZ FUS1-HIS3 ura3-52 hkr1::HYG sho1::URA3* | (PITONIAK *et al.* 2009) |
| PC2683 [b] | *MAT***a** *ade2-1 his3-11,15 leu2-3,112 ura3-1 trp1-1 can1-100 yps1Δ::LEU2 yps2Δ::HIS3 yps3Δ::KanMX4 yps6Δ::KanMX4 yps7Δ::KanMX4* (*5yps*Δ strain) | (KRYSAN *et al.* 2005) |
| PC2740 | *MAT***a** *ste4 FUS1-lacZ FUS1-HIS3 ura3-52 HA-HKR1* | (PITONIAK *et al.* 2009) |
| PC2850 | *MAT***a** *ste4 FUS1-lacZ FUS1-HIS3 ura3-52 hkr1::HYG sho1::URA3 msb2::NAT* | (PITONIAK *et al.* 2009) |
| PC3861 | *MAT***a** *ste4 FUS1-lacZ FUS1-HIS3 ura3-52 ste11::NAT* | (CULLEN *et al.* 2004) |
| PC6580 | *MAT***a** *ste4 FUS1-lacZ FUS1-HIS3 ura3-52 ssk1::NAT* | (PITONIAK *et al.* 2009) |
| PC7883 | *MAT***a** *ste4 FUS1-lacZ FUS1-HIS3 ura3-52 hkr1::KanMX6 ssk1::NAT* | This study |
| PC8102 | *MAT***a** *ste4 FUS1-lacZ FUS1-HIS3 ura3-52 msb2Δ::HYG hkr1Δ::KanMX6 ssk1Δ::NAT* | This study |
| PC8193 [c] | *MAT***a** *his4-619 leu2-3,112 ura3-52* | (TRAN AND BRODSKY 2012) |
| PC8195 [c] | *MAT***a** *lys2-801 leu2-3 ura3-52 cdc48-10(ts)* | (TRAN AND BRODSKY 2012) |
| PC8205 | *MAT***a** *ste4 FUS1-lacZ FUS1-HIS3 ura3-52 trp1::NAT Vph1-mCherry::TRP1* | This study |
| PC8206 | *MAT***a** *ste4 FUS1-lacZ FUS1-HIS3 ura3-52 trp1::NAT* Elo3-mCherry::*TRP1* | This study |
| PC8371 [b] | *MAT***a** *ade2-1 his3-11,15 leu2-3,112 ura3-1 trp1-1 can1-100 ssk1*Δ::*NAT* | This study |
| PC8373 [b] | *MAT***a** *ade2-1 his3-11,15 leu2-3,112 ura3-1 trp1-1 can1-100 yps1Δ::LEU2 yps2Δ::HIS3 yps3Δ::KanMX4 yps6Δ::KanMX4 yps7Δ::KanMX4* (*5yps*Δ strain) *ssk1*Δ::*NAT* | This study |
| PC8493 | *MAT***a** *ste4 FUS1-lacZ FUS1-HIS3 ura3-52 sho1::HYG ssk1::NAT msb2::KanMX6* | This study |

a. All strains are from the Σ1278b background unless otherwise indicated.
b. Strains from the W303 background.
c. Brodsky lab strain.

Table S2. Plasmids used in the study.

| Plasmid Number | Description | Reference |
|---|---|---|
| PC8248 | pHA-HKR1CEN/URA | This study |
| PC3507 | pGAL-YPS1 CEN/URA | (KRYSAN *et al.* 2005) |
| PC1456 | pHA-MSB2 CEN/URA | (VADAIE *et al.* 2008) |
| PC2582 | pHA-MSB2-GFP CEN/URA | (VADAIE *et al.* 2008) |
| PC4204 | pRS316 CEN/URA | (SIKORSKI AND HIETER 1989) |
| PC8050 | ZJOM65, VPH1-mCherry CEN/TRP | (ZHU *et al.* 2019) |
| PC8053 | ZJOM149, ELO3-mCherry CEN/TRP | (ZHU *et al.* 2019) |
| PC7936 | pHA-MSB2$^{\Delta 100\text{-}818}$ CEN/URA | This study |
| PC8169 | pHA-MSB2$^{\Delta 100\text{-}818}$-GFP CEN/URA | This study |
| PC8338 | pHA-MSB2$^{Gibson}$ | This study |
| PC7931 | pHA-MSB2-W961C CEN/URA | This study |
| PC7932 | pHA-MSB2-L989F CEN/URA | This study |
| PC7933 | pHA-MSB2-P990A CEN/URA | This study |
| PC7935 | pHA-MSB2-P990H CEN/URA | This study |
| PC7950 | pHA-MSB2-N1088A CEN/URA | This study |
| PC7951 | pHA-MSB2-N1049A CEN/URA | This study |
| PC7952 | pHA-MSB2-M1098G CEN/URA | This study |
| PC7953 | pHA-MSB2-N1175A CEN/URA | This study |
| PC7954 | pHA-MSB2-D1060A CEN/URA | This study |
| PC7956 | pHA-MSB2-F1012L CEN/URA | This study |
| PC7958 | pHA-MSB2-W960C CEN/URA | This study |
| PC7959 | pHA-MSB2-T964C CEN/URA | This study |
| PC7960 | pHA-MSB2-I962C$^{RPTS}$ CEN/URA | This study |
| PC7961 | pHA-MSB2-P963C CEN/URA | This study |
| PC7962 | pHA-MSB2-F1012A CEN/URA | This study |
| PC7963 | pHA-MSB2-K1014A CEN/URA | This study |
| PC7964 | pHA-MSB2-T1278A CEN/URA | This study |
| PC7965 | pHA-MSB2-T1280A CEN/URA | This study |
| PC7966 | pHA-MSB2-S1286A CEN/URA | This study |
| PC7967 | pHA-MSB2-NN1049/1088AA$^{RPTS}$ CEN/URA | This study |
| PC7968 | pHA-MSB2-NN1088/1175AA CEN/URA | This study |
| PC7969 | pHA-MSB2-1043FK/AA CEN/URA | This study |
| PC7970 | pHA-MSB2-N1175A CEN/URA | This study |
| PC7971 | pHA-MSB2-S1233A CEN/URA | This study |
| PC7973 | pHA-MSB2-S1277A CEN/URA | This study |
| PC7974 | pHA-MSB2-NN1049/1175AA CEN/URA | This study |
| PC7976 | pMYC-500-Msb2-GFP CEN/URA | This study |
| PC8045 | pHA-MSB2-NN1049/1088AA$^{TR}$ CEN/URA | This study |
| PC8046 | pHA-MSB2-NN1049/1088AA CEN/URA | This study |
| PC8054 | pHA-MSB2-W960A$^{RPTS}$ CEN/URA | This study |
| PC8055 | pHA-MSB2-S930A$^{RPTS}$ CEN/URA | This study |
| PC8056 | pHA-MSB2-I993C$^{RPTS}$ CEN/URA | This study |
| PC8057 | pHA-MSB2-V1072A$^{RPTS}$ CEN/URA | This study |
| PC8058 | pHA-MSB2-G1116A$^{RPTS}$ CEN/URA | This study |
| PC8059 | pHA-MSB2-N945A CEN/URA | This study |
| PC8060 | pHA-MSB2-L989A CEN/URA | This study |
| PC8061 | pHA-MSB2-P990R CEN/URA | This study |
| PC8062 | pHA-MSB2-P990W CEN/URA | This study |
| PC8063 | pHA-MSB2-T1009A CEN/URA | This study |
| PC8064 | pHA-MSB2-I1010A CEN/URA | This study |

| | | |
|---|---|---|
| PC8065 | pHA-MSB2-L1016A CEN/URA | This study |
| PC8066 | pHA-MSB2-N1017A CEN/URA | This study |
| PC8067 | pHA-MSB2-Y1018A CEN/URA | This study |
| PC8068 | pHA-MSB2-E1019A CEN/URA | This study |
| PC8069 | pHA-MSB2-F1020A CEN/URA | This study |
| PC8070 | pHA-MSB2-P1036A CEN/URA | This study |
| PC8071 | pHA-MSB2-L1082A CEN/URA | This study |
| PC8072 | pHA-MSB2-V1108A CEN/URA | This study |
| PC8073 | pHA-MSB2-I1112A CEN/URA | This study |
| PC8074 | pHA-MSB2-T1115A CEN/URA | This study |
| PC8075 | pHA-MSB2-G1126A CEN/URA | This study |
| PC8076 | pHA-MSB2-S1128A CEN/URA | This study |
| PC8077 | pHA-MSB2-S1232A CEN/URA | This study |
| PC8078 | pHA-MSB2-C1198A CEN/URA | This study |
| PC8079 | pHA-MSB2-S1256A$^{TR}$ CEN/URA | This study |
| PC8080 | pHA-MSB2-G1196A$^{TR}$ CEN/URA | This study |
| PC8081 | pHA-MSB2-S935A CEN/URA | This study |
| PC8082 | pHA-MSB2-P941A$^{RPTS}$ CEN/URA | This study |
| PC8083 | pHA-MSB2-L966A CEN/URA | This study |
| PC8084 | pHA-MSB2-G1011A CEN/URA | This study |
| PC8085 | pHA-MSB2-N1040A CEN/URA | This study |
| PC8086 | pHA-MSB2-P1113A CEN/URA | This study |
| PC8087 | pHA-MSB2-L1114A$^{RPTS}$ CEN/URA | This study |
| PC8088 | pHA-MSB2-G1126A$^{TR}$ CEN/URA | This study |
| PC8089 | pHA-MSB2-S1129A$^{RPTS}$ CEN/URA | This study |
| PC8090 | pHA-MSB2-G1196A$^{RPTS}$ CEN/URA | This study |
| PC8157 | pHA-MSB2-I962C$^{RPTS}$-GFP CEN/URA | This study |
| PC8224 | pHA-MSB2-F1012A-GFP CEN/URA | This study |
| PC8225 | pHA-MSB2-FKAA-GFP CEN/URA | This study |
| PC8226 | pHA-MSB2-P1113A-GFP CEN/URA | This study |
| PC8227 | pHA-MSB2-Y1018A-GFP CEN/URA | This study |
| PC8228 | pHA-MSB2-G1116A$^{RPTS}$-GFP CEN/URA | This study |
| PC8241 | pHA-MSB2-Q1030A CEN/URA | This study |
| PC8242 | pHA-MSB2-F1032A CEN/URA | This study |
| PC8243 | pHA-MSB2-A1037T CEN/URA | This study |
| PC8244 | pHA-MSB2-P1057A CEN/URA | This study |
| PC8245 | pHA-MSB2-D1061A CEN/URA | This study |
| PC8246 | pHA-MSB2-Y1073A CEN/URA | This study |
| PC8247 | pHA-MSB2-F1074A CEN/URA | This study |
| PC8250 | pHA-MSB2-P990H-GFP CEN/URA | This study |
| PC8251 | pHA-MSB2-I993C$^{RPTS}$-GFP CEN/URA | This study |
| PC8284 | pHA-MSB2-T1006A CEN/URA | This study |
| PC8285 | pHA-MSB2-P1075A CEN/URA | This study |
| PC8308 | pHA-MSB2-S1292D CEN/URA | This study |
| PC8309 | pHA-MSB2-S1297D CEN/URA | This study |
| PC8311 | pHA-MSB2-TMstop (Y1210*) CEN/URA | This study |
| PC8312 | pHA-Hkr1-stopTM (G1480*) CEN/URA | This study |
| PC8313 | pHA-Hkr1- TMstop$^{TR}$ (L1515*) CEN/URA | This study |
| PC8330 | pHA-Hkr1-Y1266A CEN/URA | This study |
| PC8331 | pHA-Hkr1-K1334A/T CEN/URA | This study |
| PC8332 | pHA-Hkr1-S1358A CEN/URA | This study |
| PC8333 | pHA-Hkr1-NP1406-1407AA CEN/URA | This study |
| PC8334 | pHA-Hkr1-K1288A CEN/URA | This study |

| | | |
|---|---|---|
| PC8335 | pHA-Hkr1-F1282A CEN/URA | This study |
| PC8336 | pHA-Hkr1-W1210C CEN/URA | This study |
| PC8337 | pHA-Hkr1-TMstop (L1515*) CEN/URA | This study |
| PC8342 | pHA-MSB2-W961A CEN/URA | This study |
| PC8343 | pHA-MSB2-Y1005A CEN/URA | This study |
| PC8344 | pHA-MSB2-I1008A CEN/URA | This study |
| PC8345 | pHA-MSB2-K1013A CEN/URA | This study |
| PC8346 | pHA-MSB2-I1031A CEN/URA | This study |
| PC8347 | pHA-MSB2-Y1034A CEN/URA | This study |
| PC8348 | pHA-MSB2-L1035A$^{RPTS}$ CEN/URA | This study |
| PC8349 | pHA-MSB2$^{\Delta100\text{-}818}$-F1012A CEN/URA | This study |
| PC8350 | pHA-MSB2$^{\Delta100\text{-}818}$-P990H CEN/URA | This study |
| PC8351 | pHA-MSB2$^{\Delta100\text{-}818}$-TMstop (Y1210*) CEN/URA | This study |
| PC8353 | pHA-MSB2$^{\Delta100\text{-}818}$-P1113A CEN/URA | This study |
| PC8354 | pHA-MSB2-stopTM (S1180*) CEN/URA | This study |
| PC8356 | pHA-MSB2-L1039A CEN/URA | This study |
| PC8357 | pHA-MSB2-P1042A CEN/URA | This study |
| PC8358 | pHA-MSB2-F1043A CEN/URA | This study |
| PC8359 | pHA-MSB2-V1052A CEN/URA | This study |
| PC8360 | pHA-MSB2-I1055A (+F1043V) CEN/URA | This study |
| PC8361 | pHA-MSB2-V1058A CEN/URA | This study |
| PC8362 | pHA-MSB2-V1072A CEN/URA | This study |
| PC8364 | pHA-MSB2-L1114A CEN/URA | This study |
| PC8365 | pHA-MSB2-G1116A CEN/URA | This study |
| PC8366 | pHA-MSB2-S1129A CEN/URA | This study |
| PC8367 | pHA-MSB2-S1135A CEN/URA | This study |
| PC8456 | pHA-MSB2$^{\Delta100\text{-}818}$-G1196A$^{TR}$ CEN/URA | This study |
| PC8460 | pHA-MSB2$^{\Delta100\text{-}818}$-stopTM (S1180*) CEN/URA | This study |
| PC8463 | pHA-MSB2$^{HKR1\text{-}SEA}$ CEN/URA | This study |
| PC8472 | pHA-HKR1$^{N}$MSB2$^{C}$ CEN/URA | This study |

Table S3. Primers used in the study.

| Primer Number | Name | Nucleotide Sequence (5' to 3') |
|---|---|---|
| 41 | Msb2-N1088 to A_fwd | CTGTCAAATCTAATTACCGCCTCTTCAAGCGCTTTTTAC |
| 42 | Msb2-N1088 to A_rev | GTAAAAAGCGCTTGAAGAGGCGGTAATTAGATTTGACAG |
| 45 | Msb2-N1049 to A_fwd | TTTAAGAACGTATTCACAGCCATTACGGTACTACAAATA |
| 46 | Msb2- N1049 to A_rev | TATTTGTAGTACCGTAATGGCTGTGAATACGTTCTTAAA |
| 49 | Msb2-1043FK to AA_fwd | GAAGCTCTGAACACACCTGCTGCGAACGTATTCACAAACATT |
| 50 | Msb2-1043FK to AA_rev | AATGTTTGTGAATACGTTCGCAGCAGGTGTGTTCAGAGCTTC |
| 53 | Msb2-1060D to A_fwd | ATAGTGCCATTACAGGCTGACTCACTCAACTAC |
| 54 | Msb2-1060D to A_rev | GTAGTTGAGTGAGTCAGCCTGTAATGGCACTAT |
| 55 | Msb2-1098M to G_fwd | TTTTACACGGATGGAGGGGGTACAGCAAAATCT |
| 56 | Msb2-1098M to G_rev | AGATTTTGCTGTACCCCCTCCATCCGTGTAAAA |
| 57 | Msb2-1175N to A_fwd | TCCAAATCTAACTCCGCCGTATCCACTTCTAGC |
| 58 | Msb2-1175N to A_rev | GCTAGAAGTGGATACGGCGGAGTTAGATTTGGA |
| 71 | Msb2-W960C_fwd | TCAAGTGACAACAATTGCTGGATTCCAACTGAG |

| | | |
|---|---|---|
| 72 | Msb2-W960C_rev | CTCAGTTGGAATCCAGCAATTGTTGTCACTTGA |
| 73 | Msb2-I962C_fwd | GACAACAATTGGTGGTGTCCAACTGAGTTAATC |
| 74 | Msb2_I962C_rev | GATTAACTCAGTTGGACACCACCAATTGTTGTC |
| 75 | Msb2-P963C_fwd | AACAATTGGTGGATTTGCACTGAGTTAATCACG |
| 76 | Msb2-P963C_rev | CGTGATTAACTCAGTGCAAATCCACCAATTGTT |
| 77 | Msb2-T964C_fwd | AATTGGTGGATTCCATGTGAGTTAATCACGCAG |
| 78 | Msb2-T964C_rev | CTGCGTGATTAACTCACATGGAATCCACCAATT |
| 81 | Msb2-F1012A_fwd | ACCCTAATCACAATAGGGGCCAAAAAGCTTTGAACTAC |
| 82 | Msb2-F1012A_rev | GTAGTTCAAAGCTTTTTGGCCCCTATTGTGATTAGGGT |
| 83 | Msb2-K1013A_fwd | CTAATCACAATAGGGTTCGCAAAAGCTTTGAACTACGAA |
| 84 | Msb2-K1013A_rev | TTCGTAGTTCAAAGCTTTTGCGAACCCTATTGTGATTAG |
| 85 | Msb2-K1014A_fwd | ATCACAATAGGGTTCAAAGCAGCTTTGAACTACGAATTT |
| 86 | Msb2-K1014A_rev | AAATTCGTAGTTCAAAGCTGCTTTGAACCCTATTGTGAT |
| 87 | Msb2-D1061A_fwd | ATAGTGCCATTACAGGATGCCTCACTCAACTACTTAGTA |
| 88 | Msb2-D1061A_rev | TACTAAGTAGTTGAGTGAGGCATCCTGTAATGGCACTAT |
| 89 | Msb2-D1109A_fwd | ATGGCTGCAATGGTTGCTTCCTCAATACCGCTA |
| 90 | Msb2-D1109A_rev | TAGCGGTATTGAGGAAGCAACCATTGCAGCCAT |
| 91 | Msb2-S1233A_fwd | ATTTCCAGTATCAGTTCAGCTGAATTTGGTGGAGAGAAA |
| 92 | Msb2-S1233A_rev | TTTCTCTCCACCAAATTCAGCTGAACTGATACTGGAAAT |
| 93 | Msb2-S1277A_fwd | GGGTTGACAAATAATGACGCAACTCCAACCAGGCACAAT |
| 94 | Msb2-S1277A_rev | ATTGTGCCTGGTTGGAGTTGCGTCATTATTTGTCAACCC |
| 95 | Msb2-T1278A_fwd | TTGACAAATAATGACTCAGCTCCAACCAGGCACAATACA |
| 96 | Msb2-T1278A_rev | TGTATTGTGCCTGGTTGGAGCTGAGTCATTATTTGTCAA |
| 97 | Msb2-T1280A_fwd | AATAATGACTCAACTCCAGCCAGGCACAATACATCGAGT |
| 98 | Msb2-T1280A_rev | ACTCGATGTATTGTGCCTGGCTGGAGTTGAGTCATTATT |
| 99 | Msb2-S1286A_fwd | ACCAGGCACAATACATCGGCTTCCATACCAAAAATTTCA |
| 100 | Msb2-S1286A_rev | TGAAATTTTTGGTATGGAAGCCGATGTATTGTGCCTGGT |
| 113 | S930A_fwd | ACTAATGTACAGACGGCTTTAACAACGGAATCG |
| 114 | S930A_rev | CGATTCCGTTGTTAAAGCCGTCTGTACATTAGT |
| 115 | S935A_fwd | AGTTTAACAACGGAAGCGACGACCGTTTTAGAA |
| 116 | S935A_rev | TTCTAAAACGGTCGTCGCTTCCGTTGTTAAACT |
| 117 | P941A_fwd | ACGACCGTTTTAGAAGCATCAACGACTAACAGT |
| 118 | P941A_rev | ACTGTTAGTCGTTGATGCTTCTAAAACGGTCGT |
| 119 | N945A_fwd | GAACCATCAACGACTGCCAGTTCCAGTACGTTT |
| 120 | N945A_rev | AAACGTACTGGAACTGGCAGTCGTTGATGGTTC |
| 123 | L989A_fwd | ACACAAACTATGACTGCGCCCCATGCAATTGCA |
| 124 | L989A_rev | TGCAATTGCATGGGGCGCAGTCATAGTTTGTGT |

| | | |
|---|---|---|
| 125 | P990R_fwd | CAAACTATGACTTTGCGCCATGCAATTGCAGCC |
| 126 | P990R_rev | GGCTGCAATTGCATGGCGCAAAGTCATAGTTTG |
| 127 | P990W_fwd | CAAACTATGACTTTGTGGCATGCAATTGCAGCC |
| 128 | P990W_rev | GGCTGCAATTGCATGCCACAAAGTCATAGTTTG |
| 131 | I993C_fwd | ACTTTGCCCCATGCATGTGCAGCCGCGACACAA |
| 132 | I993C_rev | TTGTGTCGCGGCTGCACATGCATGGGGCAAAGT |
| 133 | Y1005A_fwd | CCCGAGCCTGAGGGCGCCACCCTAATCACAATA |
| 134 | Y1005A_rev | TATTGTGATTAGGGTGGCGCCCTCAGGCTCGGG |
| 135 | T1006A_fwd | GAGCCTGAGGGCTACGCCCTAATCACAATAGGG |
| 136 | T1006A_rev | CCCTATTGTGATTAGGGCGTAGCCCTCAGGCTC |
| 137 | I1008A_fwd | GAGGGCTACACCCTAGCCACAATAGGGTTCAAA |
| 138 | I1008A_rev | TTTGAACCCTATTGTGGCTAGGGTGTAGCCCTC |
| 139 | T1009A_fwd | GGCTACACCCTAATCGCAATAGGGTTCAAAAAA |
| 140 | T1009A_rev | TTTTTTGAACCCTATTGCGATTAGGGTGTAGCC |
| 141 | I1010A_fwd | TACACCCTAATCACAGCAGGGTTCAAAAAAGCT |
| 142 | I1010A_rev | AGCTTTTTTGAACCCTGCTGTGATTAGGGTGTA |
| 143 | G1011A_fwd | ACCCTAATCACAATAGCGTTCAAAAAAGCTTTG |
| 144 | G1011A_rev | CAAAGCTTTTTTGAACGCTATTGTGATTAGGGT |
| 145 | L1016A_fwd | GGGTTCAAAAAAGCTGCGAACTACGAATTTGTT |
| 146 | L1016A_rev | AACAAATTCGTAGTTCGCAGCTTTTTTGAACCC |
| 147 | N1017A_fwd | TTCAAAAAAGCTTTGGCCTACGAATTTGTTGTA |
| 148 | N1017A_rev | TACAACAAATTCGTAGGCCAAAGCTTTTTTGAA |
| 149 | Y1018A_fwd | AAAAAAGCTTTGAACGCCGAATTTGTTGTATCA |
| 150 | Y1018A_rev | TGATACAACAAATTCGGCGTTCAAAGCTTTTTT |
| 151 | E1019A_fwd | AAAGCTTTGAACTACGCATTTGTTGTATCAGAA |
| 152 | E1019A_rev | TTCTGATACAACAAATGCGTAGTTCAAAGCTTT |
| 153 | F1020A_fwd | GCTTTGAACTACGAAGCTGTTGTATCAGAACCA |
| 154 | F1020A_rev | TGGTTCTGATACAACAGCTTCGTAGTTCAAAGC |
| 155 | Q1030A_fwd | CCAAAATCATCGGCTGCAATCTTCGGATACTTG |
| 156 | Q1030A_rev | CAAGTATCCGAAGATTGCAGCCGATGATTTTGG |
| 157 | I1031A_fwd | AAATCATCGGCTCAAGCCTTCGGATACTTGCCT |
| 158 | I1031A_rev | AGGCAAGTATCCGAAGGCTTGAGCCGATGATTT |
| 159 | F1032A_fwd | TCATCGGCTCAAATCGCCGGATACTTGCCTGAA |
| 160 | F1032A_rev | TTCAGGCAAGTATCCGGCGATTTGAGCCGATGA |
| 161 | Y1034A_fwd | GCTCAAATCTTCGGAGCCTTGCCTGAAGCTCTG |
| 162 | Y1034A_rev | CAGAGCTTCAGGCAAGGCTCCGAAGATTTGAGC |
| 163 | L1035A_fwd | CAAATCTTCGGATACGCGCCTGAAGCTCTGAAC |

| | | |
|---|---|---|
| 164 | L1035A_rev | GTTCAGAGCTTCAGGCGCGTATCCGAAGATTTG |
| 165 | P1036A_fwd | ATCTTCGGATACTTGGCTGAAGCTCTGAACACA |
| 166 | P1036A_rev | TGTGTTCAGAGCTTCAGCCAAGTATCCGAAGAT |
| 167 | L1039A_fwd | TACTTGCCTGAAGCTGCGAACACACCTTTTAAG |
| 168 | L1039A_rev | CTTAAAAGGTGTGTTCGCAGCTTCAGGCAAGTA |
| 169 | N1040A_fwd | TTGCCTGAAGCTCTGGCCACACCTTTTAAGAAC |
| 170 | N1040A_rev | GTTCTTAAAAGGTGTGGCCAGAGCTTCAGGCAA |
| 171 | P1042A_fwd | GAAGCTCTGAACACAGCTTTTAAGAACGTATTC |
| 172 | P1042A_rev | GAATACGTTCTTAAAAGCTGTGTTCAGAGCTTC |
| 173 | F1043A_fwd | GCTCTGAACACACCTGCTAAGAACGTATTCACA |
| 174 | F1043A_rev | TGTGAATACGTTCTTAGCAGGTGTGTTCAGAGC |
| 175 | V1052A_fwd | TTCACAAACATTACGGCACTACAAATAGTGCCA |
| 176 | V1052A_rev | TGGCACTATTTGTAGTGCCGTAATGTTTGTGAA |
| 177 | I1055A_fwd | ATTACGGTACTACAAGCAGTGCCATTACAGGAT |
| 178 | I1055A_rev | ATCCTGTAATGGCACTGCTTGTAGTACCGTAAT |
| 179 | P1057A_fwd | GTACTACAAATAGTGGCATTACAGGATGACTCA |
| 180 | P1057A_rev | TGAGTCATCCTGTAATGCCACTATTTGTAGTAC |
| 181 | L1058A_fwd | CTACAAATAGTGCCAGCACAGGATGACTCACTC |
| 182 | L1058A_rev | GAGTGAGTCATCCTGTGCTGGCACTATTTGTAG |
| 183 | Y1065A_fwd | GATGACTCACTCAACGCCTTAGTAAGTGTTGCT |
| 184 | Y1065A_rev | AGCAACACTTACTAAGGCGTTGAGTGAGTCATC |
| 185 | S1068A_fwd | CTCAACTACTTAGTAGCTGTTGCTGAAGTATAC |
| 186 | S1068A_rev | GTATACTTCAGCAACAGCTACTAAGTAGTTGAG |
| 187 | V1069A_fwd | AACTACTTAGTAAGTGCTGCTGAAGTATACTTT |
| 188 | V1069A_rev | AAAGTATACTTCAGCAGCACTTACTAAGTAGTT |
| 189 | V1072A_fwd | GTAAGTGTTGCTGAAGCATACTTTCCAACTGCA |
| 190 | V1072A_rev | TGCAGTTGGAAAGTATGCTTCAGCAACACTTAC |
| 191 | Y1073A_fwd | AGTGTTGCTGAAGTAGCCTTTCCAACTGCAGAA |
| 192 | Y1073A_rev | TTCTGCAGTTGGAAAGGCTACTTCAGCAACACT |
| 193 | F1074A_fwd | GTTGCTGAAGTATACGCTCCAACTGCAGAAATA |
| 194 | F1074A_rev | TATTTCTGCAGTTGGAGCGTATACTTCAGCAAC |
| 195 | P1075A_fwd | GCTGAAGTATACTTTGCAACTGCAGAAATAGAG |
| 196 | P1075A_rev | CTCTATTTCTGCAGTTGCAAAGTATACTTCAGC |
| 197 | L1082A_fwd | GCAGAAATAGAGGAGGCGTCAAATCTAATTACC |
| 198 | L1082A_rev | GGTAATTAGATTTGACGCCTCCTCTATTTCTGC |
| 199 | S1091A_fwd | ATTACCAACTCTTCAGCCGCTTTTTACACGGAT |
| 200 | S1091A_rev | ATCCGTGTAAAAAGCGGCTGAAGAGTTGGTAAT |

| | | |
|---|---|---|
| 201 | Y1094A_fwd | TCTTCAAGCGCTTTTGCCACGGATGGAATGGGT |
| 202 | Y1094A_rev | ACCCATTCCATCCGTGGCAAAAGCGCTTGAAGA |
| 203 | V1108A_fwd | TCTATGGCTGCAATGGCTGATTCCTCAATACCG |
| 204 | V1108A_rev | CGGTATTGAGGAATCAGCCATTGCAGCCATAGA |
| 205 | I1112A_fwd | ATGGTTGATTCCTCAGCACCGCTAACGGGCCTC |
| 206 | I1112A_rev | GAGGCCCGTTAGCGGTGCTGAGGAATCAACCAT |
| 207 | P1113A_fwd | GTTGATTCCTCAATAGCGCTAACGGGCCTCTTA |
| 208 | P1113A_rev | TAAGAGGCCCGTTAGCGCTATTGAGGAATCAAC |
| 209 | L1114A_fwd | GATTCCTCAATACCGGCAACGGGCCTCTTACAC |
| 210 | L1114A_rev | GTGTAAGAGGCCCGTTGCCGGTATTGAGGAATC |
| 211 | T1115A_fwd | TCCTCAATACCGCTAGCGGGCCTCTTACACGAT |
| 212 | T1115A_rev | ATCGTGTAAGAGGCCCGCTAGCGGTATTGAGGA |
| 213 | G1116A_fwd | TCAATACCGCTAACGGCCCTCTTACACGATAGT |
| 214 | G1116A_rev | ACTATCGTGTAAGAGGGCCGTTAGCGGTATTGA |
| 217 | G1126A_fwd | AGTAACAGCAACTCTGCCGGATCTTCGGACGGA |
| 218 | G1126A_rev | TCCGTCCGAAGATCCGGCAGAGTTGCTGTTACT |
| 219 | S1128A_fwd | AGCAACTCTGGCGGAGCTTCGGACGGATCCTCC |
| 220 | S1128A_rev | GGAGGATCCGTCCGAAGCTCCGCCAGAGTTGCT |
| 221 | S1129A_fwd | AACTCTGGCGGATCTGCGGACGGATCCTCCTCC |
| 222 | S1129A_rev | GGAGGAGGATCCGTCCGCAGATCCGCCAGAGTT |
| 223 | S1135A_fwd | GACGGATCCTCCTCCGCTAATTCGAACTCAGGA |
| 224 | S1135A_rev | TCCTGAGTTCGAATTAGCGGAGGAGGATCCGTC |
| 225 | N1224A_fwd | CAAGAAATTATCAAGGCCCCAGAAATTTCCAGT |
| 226 | N1224A_rev | ACTGGAAATTTCTGGGGCCTTGATAATTTCTTG |
| 227 | S1232A_fwd | ATTTCCAGTATCAGTGCAAGTGAATTTGGTGGA |
| 228 | S1232A_rev | TCCACCAAATTCACTTGCACTGATACTGGAAAT |
| 229 | N1243A_fwd | GAGAAAAATTACAATGCTGAAAAGAGAATGAGC |
| 230 | N1243A_rev | GCTCATTCTCTTTTCAGCATTGTAATTTTTCTC |
| 231 | S1252A_fwd | ATGAGCGTTCAAGAAGCCATAACACAATCTATG |
| 232 | S1252A_rev | CATAGATTGTGTTATGGCTTCTTGAACGCTCAT |
| 233 | S1256A_fwd | GAATCCATAACACAAGCTATGCGAATTCAAAAT |
| 234 | S1256A_rev | ATTTTGAATTCGCATAGCTTGTGTTATGGATTC |
| 235 | W961A_fwd | AGTGACAACAATTGGGCGATTCCAACTGAGTTA |
| 236 | W961A_rev | TAACTCAGTTGGAATCGCCCAATTGTTGTCACT |
| 239 | G1196A_fwd | ATCGGCGTTGTTGTTGCTGGATGCTTATATATT |
| 240 | G1196A_rev | AATATATAAGCATCCAGCAACAACAACGCCGAT |

| | | |
|---|---|---|
| 247 | Msb2_popin_990aa_fwd | ACTGCATCTTCTACCGTTGGAGGAACACAAACTATGACTTTGC CCAGGGAACAAAAGCTGG |
| 248 | Msb2_popin_926aa_rev | TGGTTCTAAAACGGTCGTCGATTCCGTTGTTAAACTCGTCTGT ACCTGTAGGGCGAATTGG |
| 249 | Msb2_popin_1174aa_fwd | TATCAAGATGCCGGTACTTTGGAATATTCATCCAAATCTAACT CCAGGGAACAAAAGCTGG |
| 250 | Msb2_popin_1110aa_rev | AGAGTTGCTGTTACTATCGTGTAAGAGGCCCGTTAGCGGTATT GACTGTAGGGCGAATTGG |
| 251 | Msb2_popin_1043aa_fwd | GCTCAAATCTTCGGATACTTGCCTGAAGCTCTGAACACACCTT TTAGGGAACAAAAGCTGG |
| 252 | Msb2_popin_1043aa_rev | TAATGGCACTATTTGTAGTACCGTAATGTTTGTGAATACGTTC TTCTGTAGGGCGAATTGG |
| 259 | Hkr1_frag1_F | AGGTCGACGGTATCGATAAGCTTGATATCGATGTACAATATT CATACTAG |
| 260 | Hkr1_frag1_R | AGCACTTAAGTCATAATATGTGGAAAGCTC |
| 261 | Hkr1_frag2_F | GAGCTTTCCACATATTATGACTTAAGTGCT |
| 262 | Hkr1_frag2_R | AATTGAGATTTTGTAAAAGACCGAGAATA |
| 263 | Hkr1_frag3_F | TATTCTCGGTCTTTTACAAAATCTCAATT |
| 264 | Hkr1_frag3_R | CTAGAACTAGTGGATCCCCCGGGCTGCAGGATTCCGAAAATA TTAAAAAA |
| 265 | Hkr1_frag1_F2_XhoI | AGGGAACAAAAGCTGGTACCGGGCCCCCCCATGTACAATATT CATACTAG |
| 266 | Hkr1_frag3_R2_BamHI | ACCGCGGTGGCGGCCGCTCTAGAACTAGTGATTCCGAAAATA TTAAAAAA |
| 275 | Msb2-S1180*_f | AACGTATCCACTTCTTGAAAATCAAAGAAAAAA |
| 276 | Msb2-S1180*_r | TTTTTTCTTTGATTTTCAAGAAGTGGATACGTT |
| 277 | Msb2-Y1210*_f | ATTTTTGCTTTCAAGTAAATCATAAGAAGGCGG |
| 278 | Msb2-Y1210*_r | CCGCCTTCTTATGATTTACTTGAAAGCAAAAAT |
| 279 | Hkr1-G1480*_f | GATTCTGCATCTGCATGAAAGTATGCTGTAAAA |
| 280 | Hkr1-G1480*_r | TTTTACAGCATACTTTCATGCAGATGCAGAATC |
| 281 | Hkr1-L1515*_f | CATAGAAATATATTATAAAAAAGACACCCAAGG |
| 282 | Hkr1-L1515*_r | CCTTGGGTGTCTTTTTTATAATATATTTCTATG |
| 283 | Msb2-frag1_SalI_f | CAAAAGCTGGTACCGGGCCCCCCCTCGAGGTCTGCAGGGAAG GCTGCCGC |
| 284 | Msb2-frag1_r | AGTTGGAGAACTAGGGCTACTGAAATCCGC |
| 285 | Msb2-frag2_f | GCGGATTTCAGTAGCCCTAGTTCTCCAACT |
| 286 | Msb2-frag2_r | GGAACTGTTAGTCGTTGATGGTTCTAAAAC |
| 287 | Msb2-frag3_f | GTTTTAGAACCATCAACGACTAACAGTTCC |
| 288 | Msb2-frag3_BamHI_r | ACCGCGGTGGCGGCCGCTCTAGAACTAGTGAAAGGCCTGCAG TCTAACTT |
| 291 | Hkr1-Y1266A_f | ACGGCCGCTTTGAATGCTGTATTTTTGGTTCAA |
| 292 | Hkr1-Y1266A_r | TTGAACCAAAAATACAGCATTCAAAGCGGCCGT |
| 293 | Hkr1-F1260A_f | TTAATTACCATCGGAGCCACGGCCGCTTTGAAT |
| 294 | Hkr1-F1260A_r | ATTCAAAGCGGCCGTGGCTCCGATGGTAATTAA |

| | | |
|---|---|---|
| 295 | Hkr1-1334A_f | TTATCAGTTGTCAAGGCAAAAAAAAACCAGCAG |
| 296 | Hkr1-1334A_r | CTGCTGGTTTTTTTTTGCCTTGACAACTGATAA |
| 297 | Hkr1-S1358A_f | CCACAAGTCGACACAGCCTCAATAGCGGTGAAA |
| 298 | Hkr1-S1358A_r | TTTCACCGCTATTGAGGCTGTGTCGACTTGTGG |
| 299 | Hkr1-NP1406/07AA_f | TCGACTCTTTATAGCGCTGCTCAAACCCCACTGCGA |
| 300 | Hkr1-NP1406/07AA_r | TCGCAGTGGGGTTTGAGCAGCGCTATAAAGAGTCGA |
| 301 | Hkr1-K1288A_f | TTACCTCTCGTTTTGGCGTATCCCTTTTCAAAC |
| 302 | Hkr1-K1288A_r | GTTTGAAAAGGGATACGCCAAAACGAGAGGTAA |
| 303 | Hkr1-F1282A_f | GCTCAAATCTTCAACGCTTTACCTCTCGTTTTG |
| 304 | Hkr1-F1282A_r | CAAAACGAGAGGTAAAGCGTTGAAGATTTGAGC |
| 305 | Hkr1-W1210C_f | CCAAATAGTTACGCATGCTTACCTACCGCTATT |
| 306 | Hkr1-W1210C_r | AATAGCGGTAGGTAAGCATGCGTAACTATTTGG |

*Fig._S1*

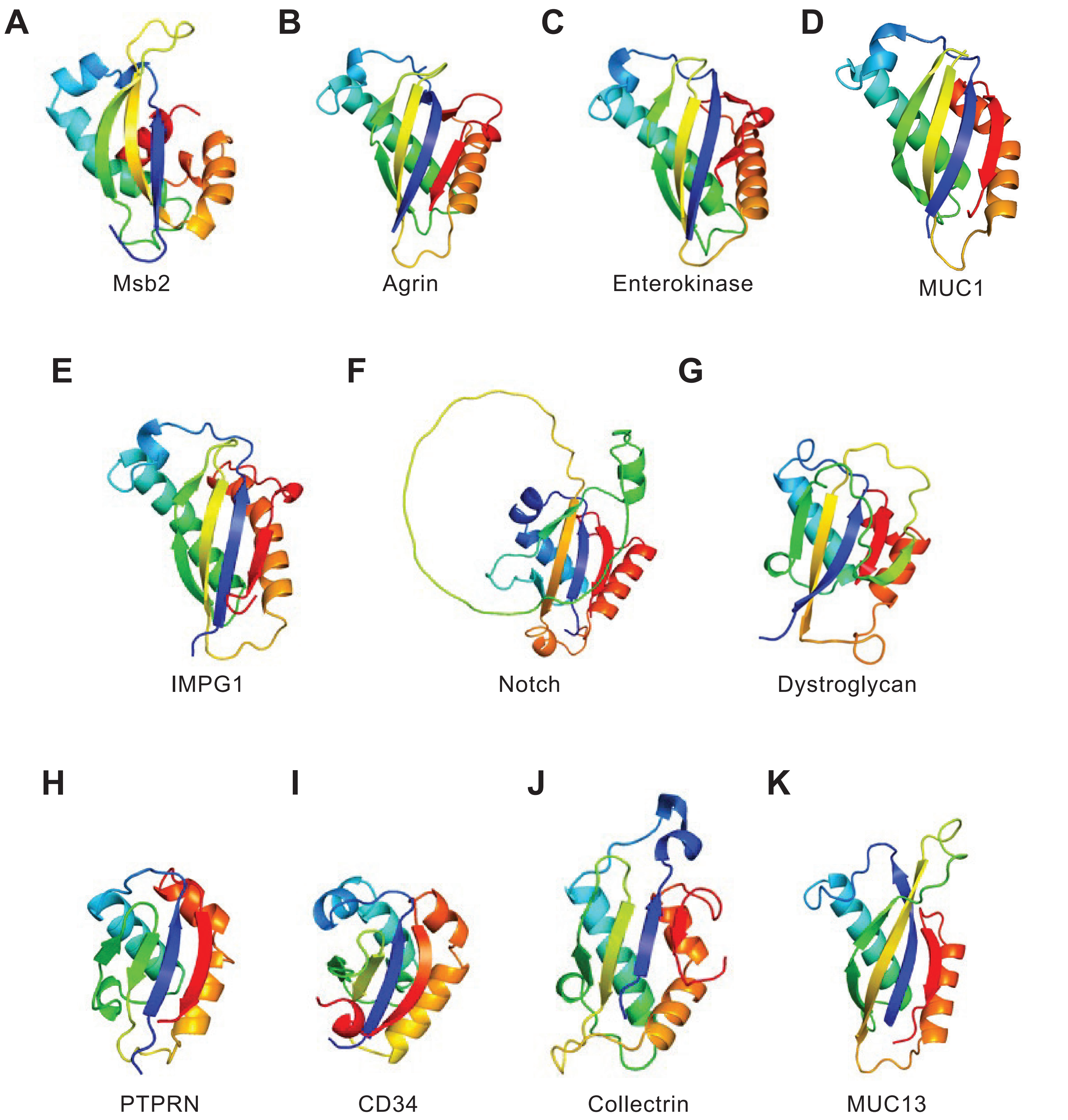
A
Msb2
B
Agrin
C
Enterokinase
D
MUC1
E
IMPG1
F
Notch
G
Dystroglycan
H
PTPRN
I
CD34
J
Collectrin
K
MUC13

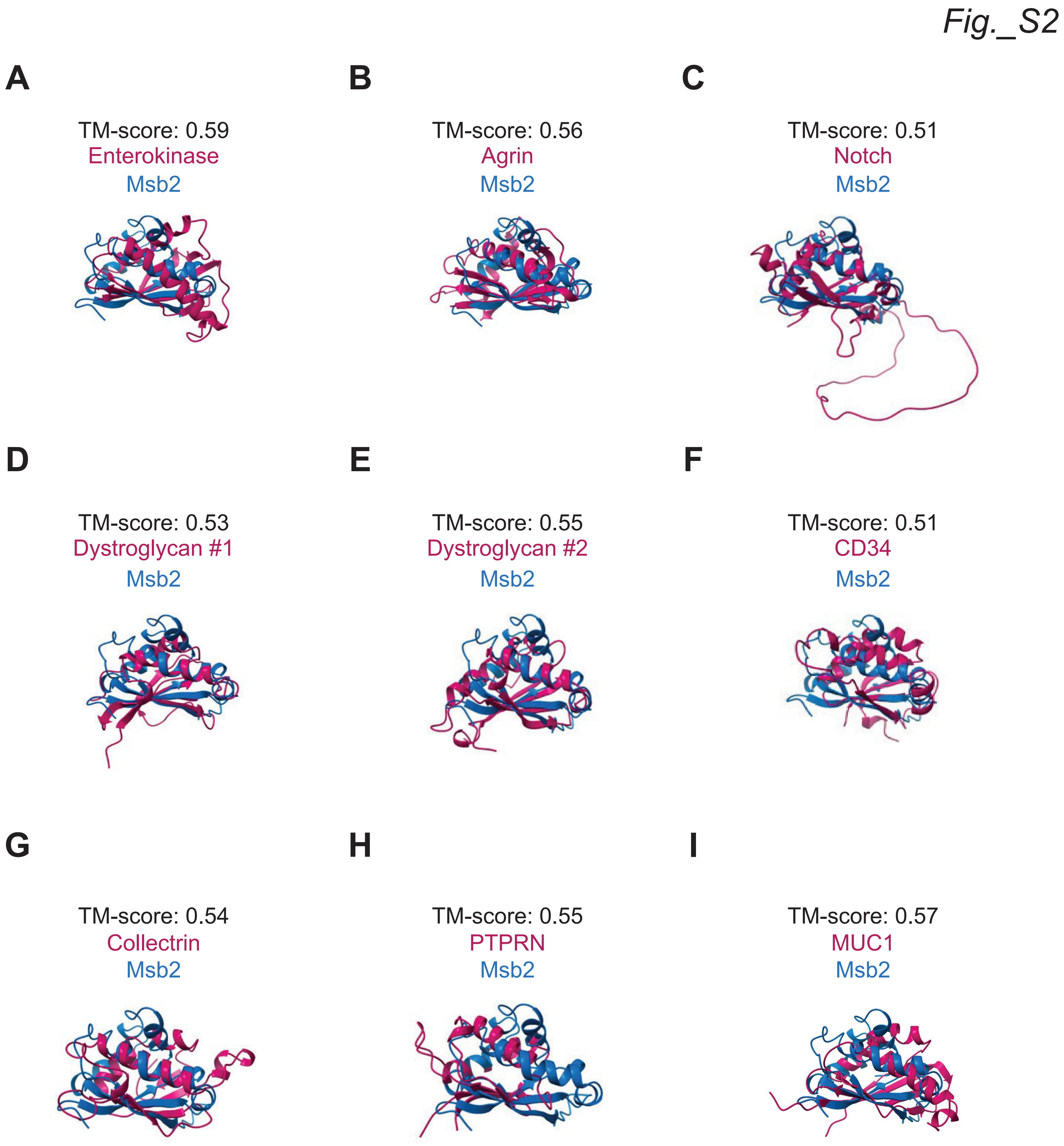
Fig._S2
A
TM-score: 0.59
Enterokinase
Msb2
B
TM-score: 0.56
Agrin
Msb2
C
TM-score: 0.51
Notch
Msb2
D
TM-score: 0.53
Dystroglycan #1
Msb2
E
TM-score: 0.55
Dystroglycan #2
Msb2
F
TM-score: 0.51
CD34
Msb2
G
TM-score: 0.54
Collectrin
Msb2
H
TM-score: 0.55
PTPRN
Msb2
I
TM-score: 0.57
MUC1
Msb2

### Fig._S3

| Name (ID) | Sp. | %Id. (SEA) | (SEA) | |
|---|---|---|---|---|
| | | | | *** * * * ** ** * |
| | | | | eeeeee hhhhh hhhhhhhhhhhhhh eeeeeeee eeeeeeeee hhhhhhhhhhhhhh eeeeee |
| P15941_MUC1 | *H.sa* | 14 | (1040-1148) | -SFFFLSFHIS(13)DY YQE--LQRDISEMFLQIY KQGGFLG LSNIKFRP--GSVVVQLTLAFREGTI-NVHDVETQFNQYKTEAASRYNLT ISDVSVSDVPFPF |
| Q14118_DAG1 #1 | *H.sa* | 9 | (187-301) | -PVTVLTVILDADLTKM TPK--QRIDLLHRMRSFSEV---- E LHNMKLVP(26)ALLSWKLGCSLNQ-NSVPDI-HGVEAPAREGAMS(5)P VVGWHIA |
| Q14118_DAG1 #2 | *H.sa* | 4 | (604-709) | -APARFKAKFVG(6)ND IHK---KIALVKKLAFAF GDRNCST ITLQNITR--GSIVVEWTNN(6)CPKEQIAGLSRRI AEDDGKPRP(10)K ATSITVTGSGSCRHLQ |
| Q17R60_IMPG1 #1 | *H.sa* | 6 | (232-354) | -EEQRVELSVSL(6)AE LAD(5)YQELAGKSQLQM Q(8)FKK IHVLGFRPK(7)TEMQLTAIFK(40)LTATDLKRLI SKALEEEQSLDVGT IQ-FTDEI |
| Q17R60_IMPG1 #2 | *H.sa* | 10 | (571-684) | -RELVVFFSLRV(6)ND LFN(5)YRALEQQFTQLL VPYL(6) FKQLEILNF(4)VIVNSKMKFA(4)YNLTKAVHGVLEDFRSAAAQQLHLE IDSYSLNIEPADQ |
| O00468_AGRN | *H.sa* | 4 | (1131-1252) | TKVFQGVLELE(17)SE LFGE-TARSIESTLDDLF RNS(7)S VRLRDLGPG-KSVRAIVDVHFDPTTAFRAPDVARAL LRQIQVSRRR(7)L QEHVRFMDFDW |
| A0A0C9W2Q7 | *H.pi* | 26 | (34-149) | LPTSDSLINVL(4)LNWEWVA(5)PGQIFEYFPQVIA(6)AGDVQNFELIVY(12)QLGTIWQGYMP----SDK V-STLASLISNQK-SVFY(6)IPSTLAA--CVD |
| A0A5B0R8H7 | **P.gr* | 24 | (808-920) | PANSTLISILFTDALNWPWLV(5)SSQVLVFMPELIAS(6)DQVVTQSLQAY(11)SMLTLWLGYIP----SQY V-GQLQAMIRAPQ-SRFY(6)LRALAK---TVD |
| A0A0D1E8D6 | **U.ma* | 21 | (742-848) | QPPNTTSIGLL(4)MKWTWVI(5)TAQIFAFMPGLVG(6)SSKVSTVKLVGY(16)TARTLYMAYVP----SSS V-DDIQAMVSNIS-SPFY(6)APQQLAS--QVD |
| R4X9N6 | *T.de* | 28 | (539-645) | PNSSQTIVQVG(4)LNYPFVV(5)TAQILAYTPKAVA(5)PQSGILANSLRP(4)DYVATVAYLVIPSADLSQ- L-EELWNNTN----SVLY(6)LQSLVV---LID |
| C0NVK4 | **H.ca* | 28 | (469-577) | -PADSRLIQVG(4)LSYEFVV(5)NLQIFTFTPKGVS(5)SLDLVKAHSIQ(9)RMYTPTLVLMYIP-----ED LVNELSLQIHSPY-SRLY(6)IAKLMS---MID |
| A0A0M8N1N7 | *E.we* | 32 | (37-144) | -PDGTTLIQIG(4)LNWYFVV(5)ATQIFAELPIAMC(8)KIKFARLEPLDT(4)GYYTTLARVYYP----TNL V-DKLRMDLWTPN-SLLY(6)VQNLTAQI |
| A0A6A6TRK9 | *L.ma* | 24 | (994-1100) | -PANSFLGQIG(4)LNYPFVS(4)GNQIFAYTVPALT(5)SADKVVMQGLKP(7)GFITTLAMFYIP-----AD LKTTLQAQLLTPADNLWH(5)VRSLTDLI |
| A0A2H1A754 | **C.au* | 44 | (866-976) | PGKDYDLIYIG(4)LNYPFVV(5)SAQIFEYLPRVLQ(8)YNGTLVSRLVP(5)INYVITVAEVYFPSNAVKA- L-SSLILDDS----SKLY(3)QTEAKLAA-LID |
| A0A1D8PGF8 | **C.al* | 40 | (1141-1252) | PGFDYQVITVG(4)LNYPFVV(5)SAQIFQYLPRVLK(7)SLQNVSVRRLI(6)IDYTITVAEVYFP----KDS V-KALGSFITTPG-SAIY(4)SV-LQALASLIDS |
| Q750D6 | *E.go* | 42 | (806-915) | PKKDYTLITIG(4)LNYPFIA(5)NAQIFDYLPGILN(8)DISVIQLVPLKANSKDYIATIAQVYFP----SEK V-DELAKLVVDTN-SELY(4)AN-MKVFASLIDP |
| Q6CST7 | *K.la* | 50 | (576-687) | QPAGYTLITVG(4)LNYPFTV(5)NAQIFEFLPKVLN(11)TSVLELVPLKVTSLDYLVTVAKVYFP----EDS V-STLQAQILNSS-SVIYNNDVTALKQLSSYIDA |
| A0A0W0CGP0 | *C.gl* | 47 | (634-744) | VPEGYSVITIG(4)LNYPFVV(5)SAQIMYYLPKLLN(8)DIKTVQLMPLEIQDSGYLSTVALLFFP----TSE I-QSLSNMVKNTS-SSLY(6)ITQSLSA--LIDP |
| P32334_Msb2 | *S.ce* | 100 | (1000-1110) | EPEGYTLITIG(4)LNYEFVV(5)SAQIFGYLPEALNT(8)ITVLQIVPLQDDSLNYLVSVAEVYFP----TAE I-EELSNLITNSS-SAFY(6)AKSMAA---MVDS |
| | | | | hhh hhhhh |

### Fig._S4

**A**

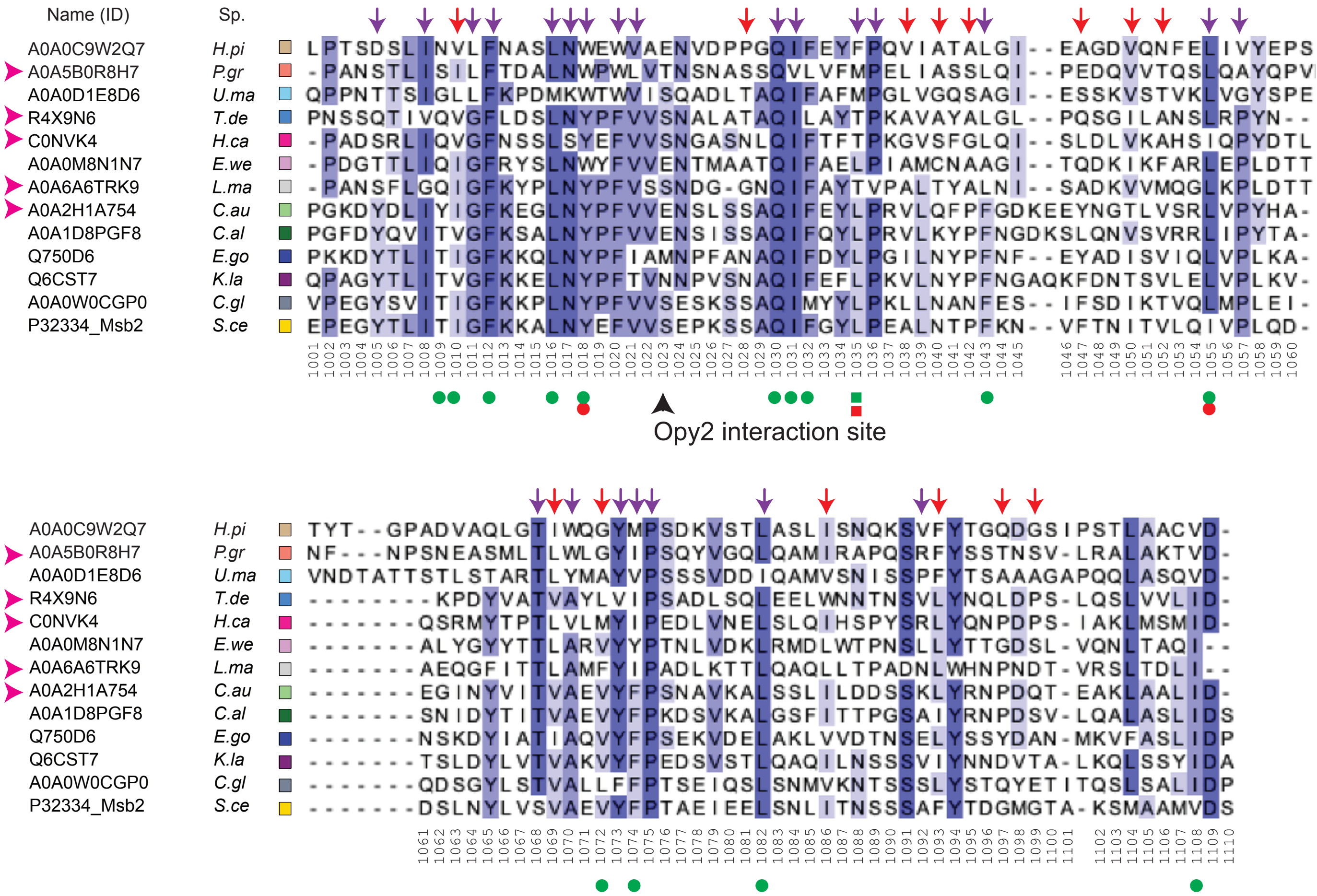


Key
- (green dot) critical for fMAPK(LOF and partial LOF)
- (red dot) critical for HOG
- (purple arrow) conserved and critical for structure
- (red arrow) critical for structure
- (pink arrowhead) new Msb2 homologs

**B**

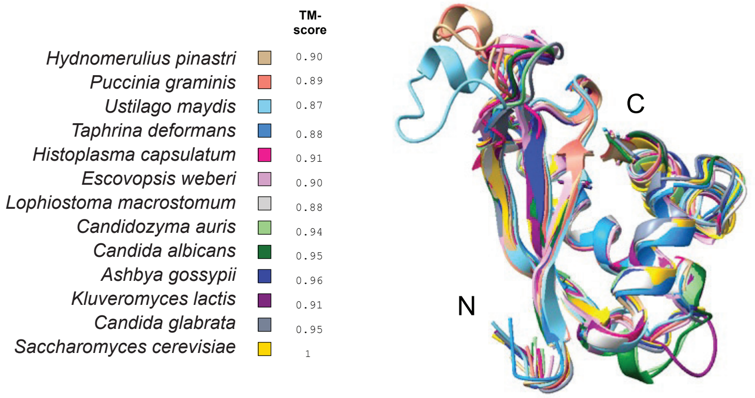

Fig._S5

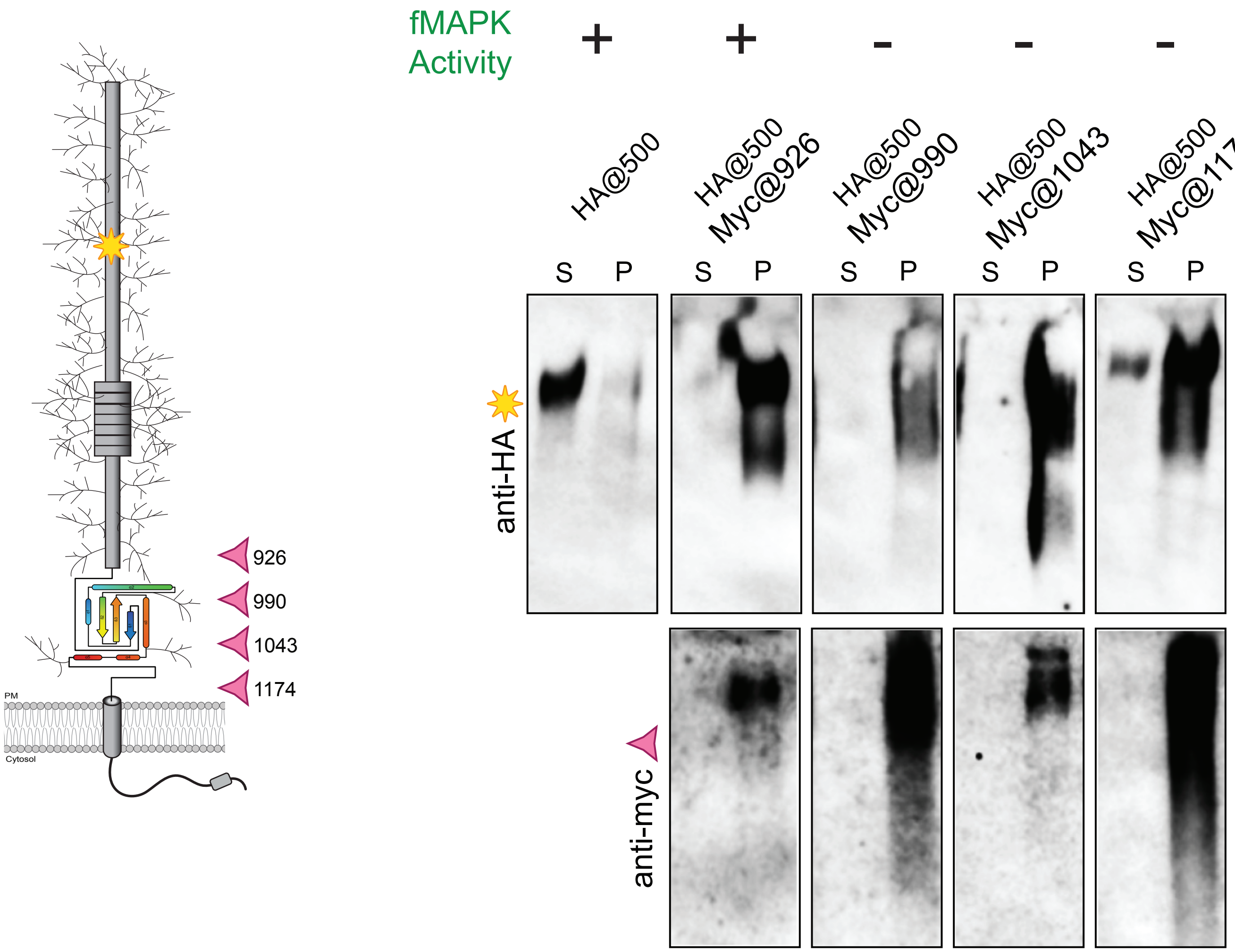

926
990
1043
1174
PM
Cytosol
fMAPK Activity
+
+
-
-
-
HA@500
HA@500 Myc@926
HA@500 Myc@990
HA@500 Myc@1043
HA@500 Myc@1174
S P
S P
S P
S P
S P
anti-HA
anti-myc

Fig._S6

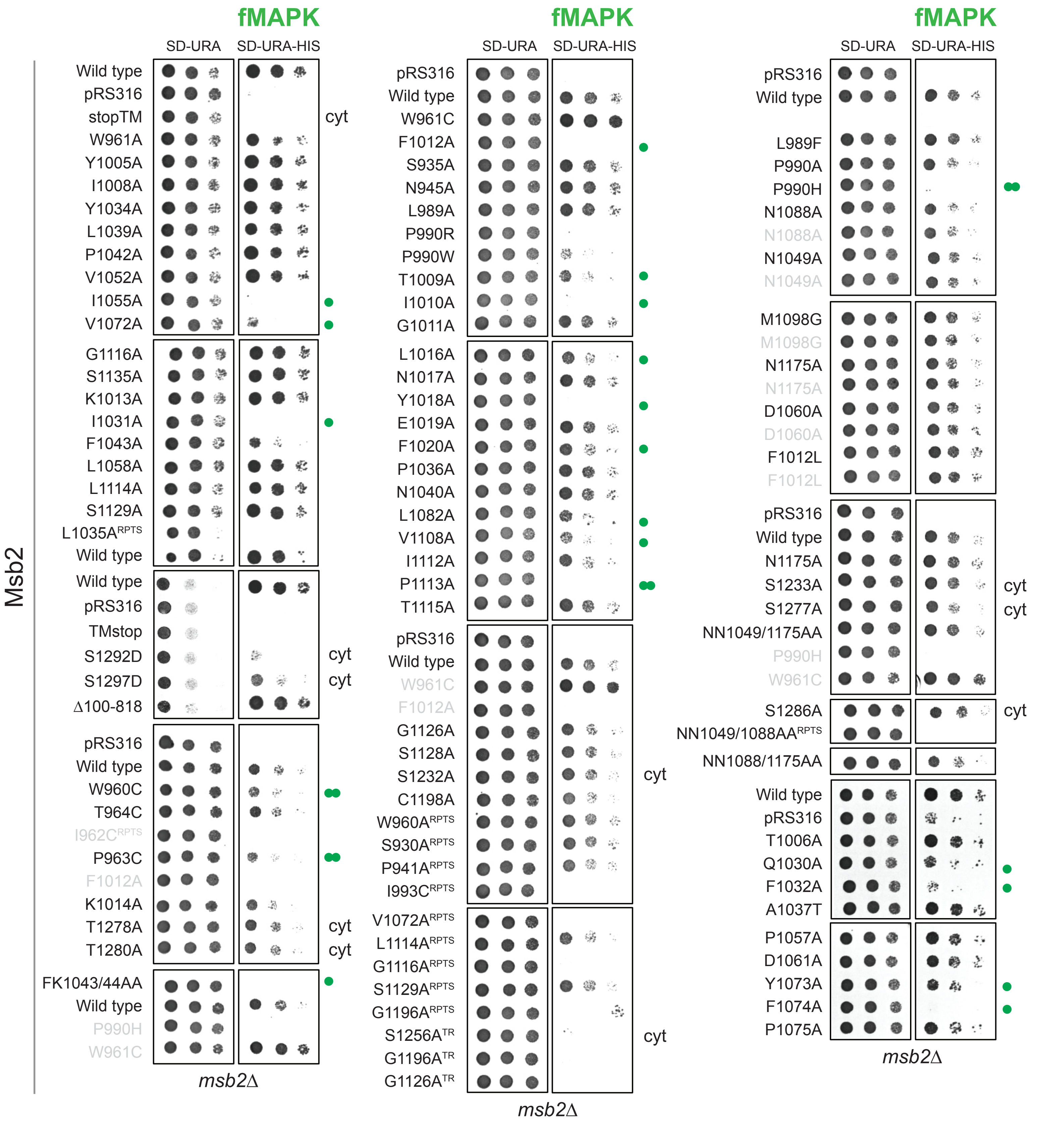


Key:

cyt cytoplasmic tail residue

RPTS contains 1 or more repeats/insertions

TR truncated due to frameshift after the residue

● point mutations defective for fMAPK (LOF and partial LOF)

●● variants in the flanking region defective for fMAPK

Summary of tested residues:

39 in the SEA domain
17 in the flanking region
14 RPTS insertions
9 cytoplasmic

Note- variant duplicates are in grey; spots were cropped from different plates and arranged

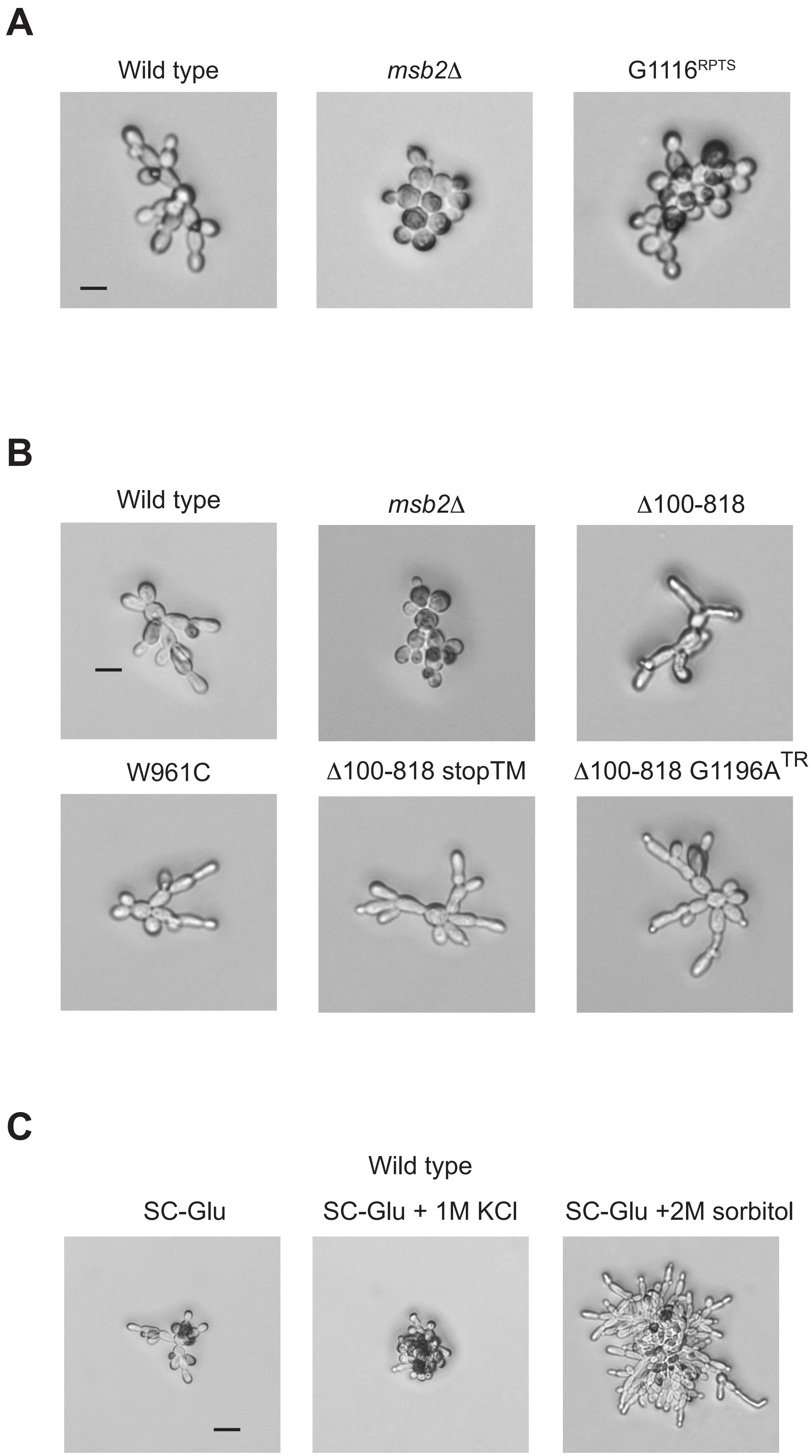
A
Wild type
msb2Δ
G1116RPTS
B
Wild type
msb2Δ
Δ100-818
W961C
Δ100-818 stopTM
Δ100-818 G1196ATR
C
Wild type
SC-Glu
SC-Glu + 1M KCl
SC-Glu +2M sorbitol

*Fig._S7*

*Fig._S8*

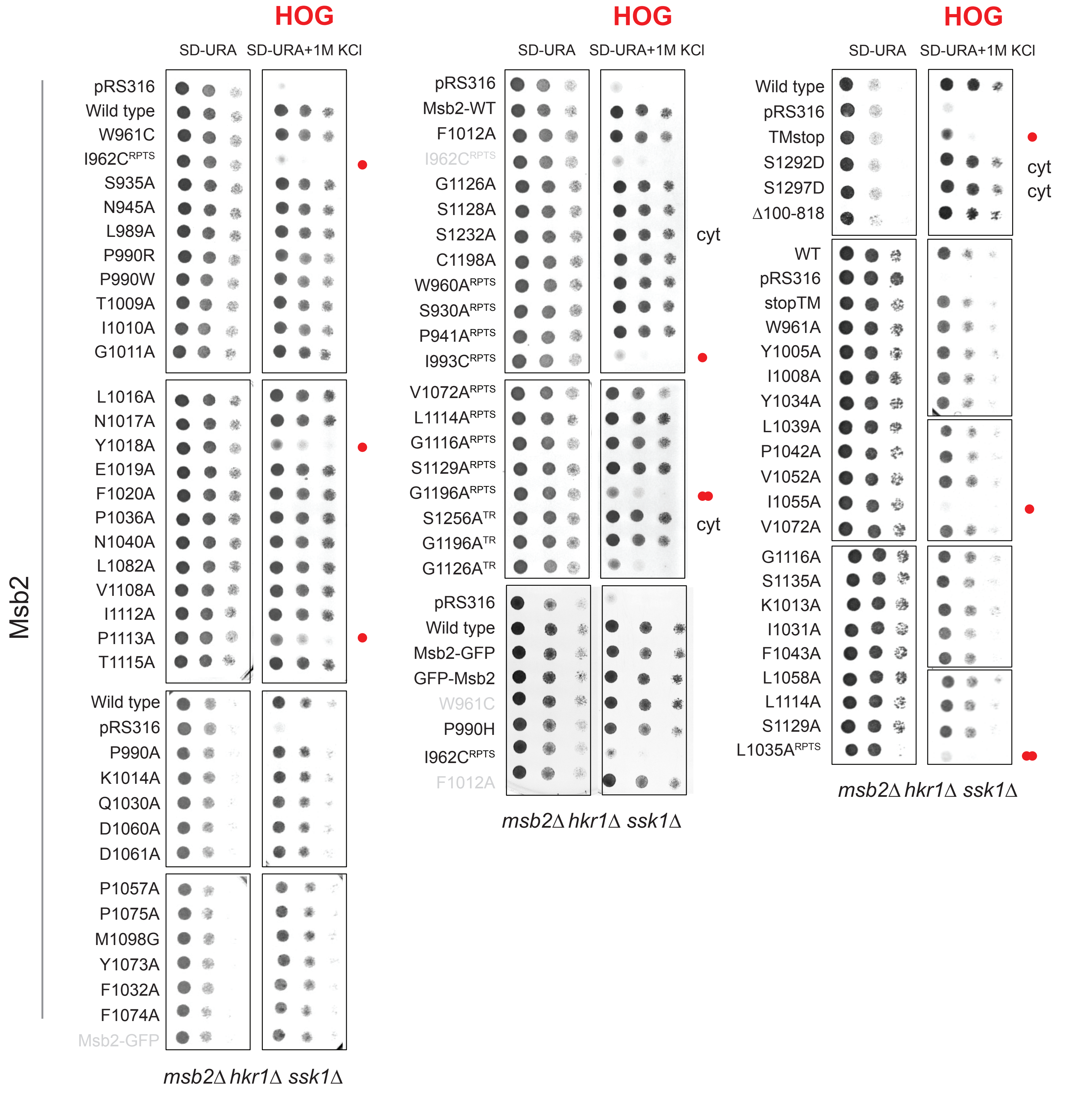


Key:

cyt cytoplasmic tail residue

RPTS contains 1 or more repeats/insertions

TR truncated due to frameshift after the residue

● point mutations defective for HOG activity

●● RPTS mutants defective for HOG activity

Note- variant duplicates are in grey; spots were cropped from different plates and arranged

Fig._S9

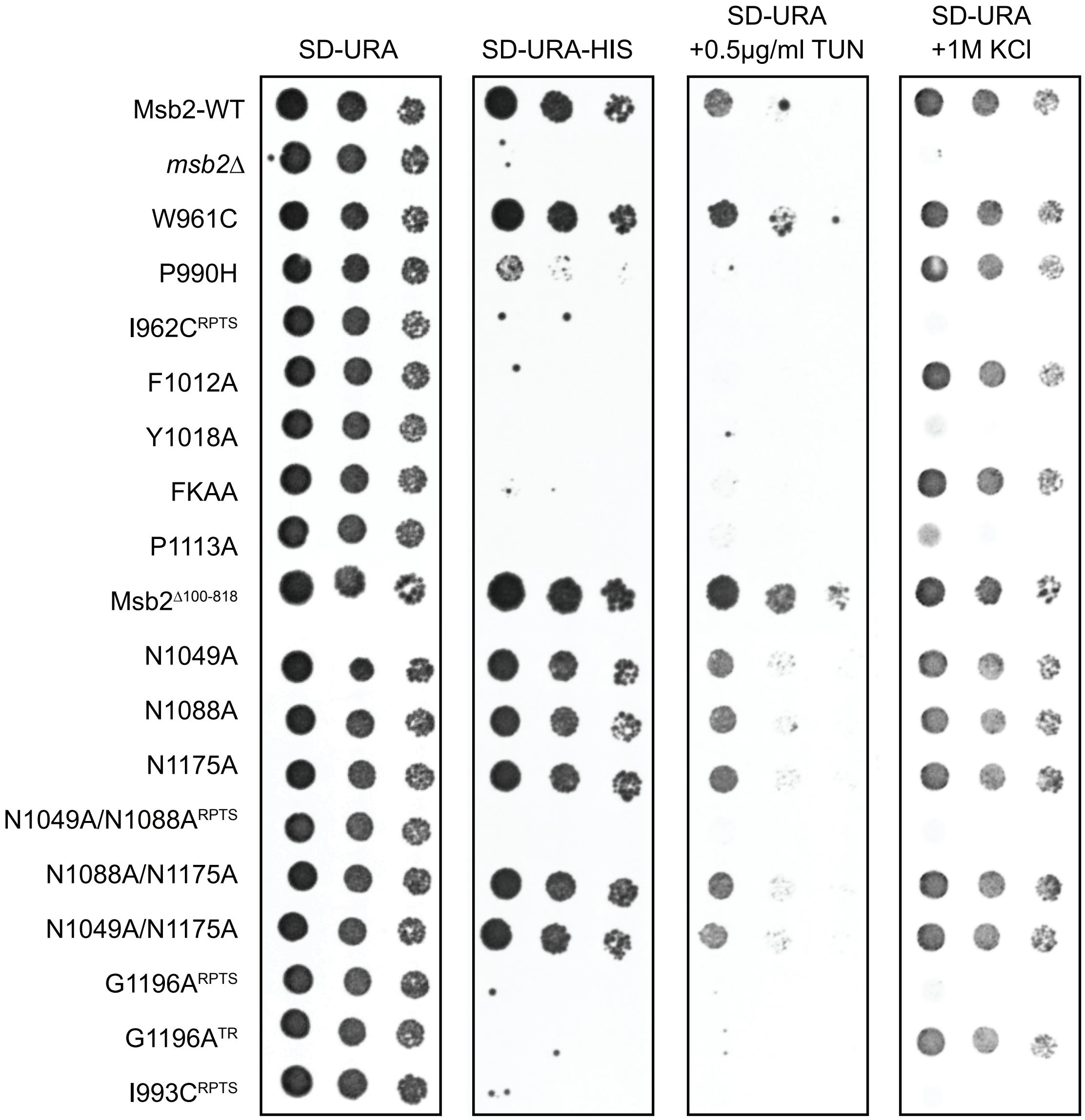
SD-URA
SD-URA-HIS
SD-URA
+0.5µg/ml TUN
SD-URA
+1M KCl
Msb2-WT
msb2Δ
W961C
P990H
I962CRPTS
F1012A
Y1018A
FKAA
P1113A
Msb2Δ100-818
N1049A
N1088A
N1175A
N1049A/N1088ARPTS
N1088A/N1175A
N1049A/N1175A
G1196ARPTS
G1196ATR
I993CRPTS

*Fig._S10*

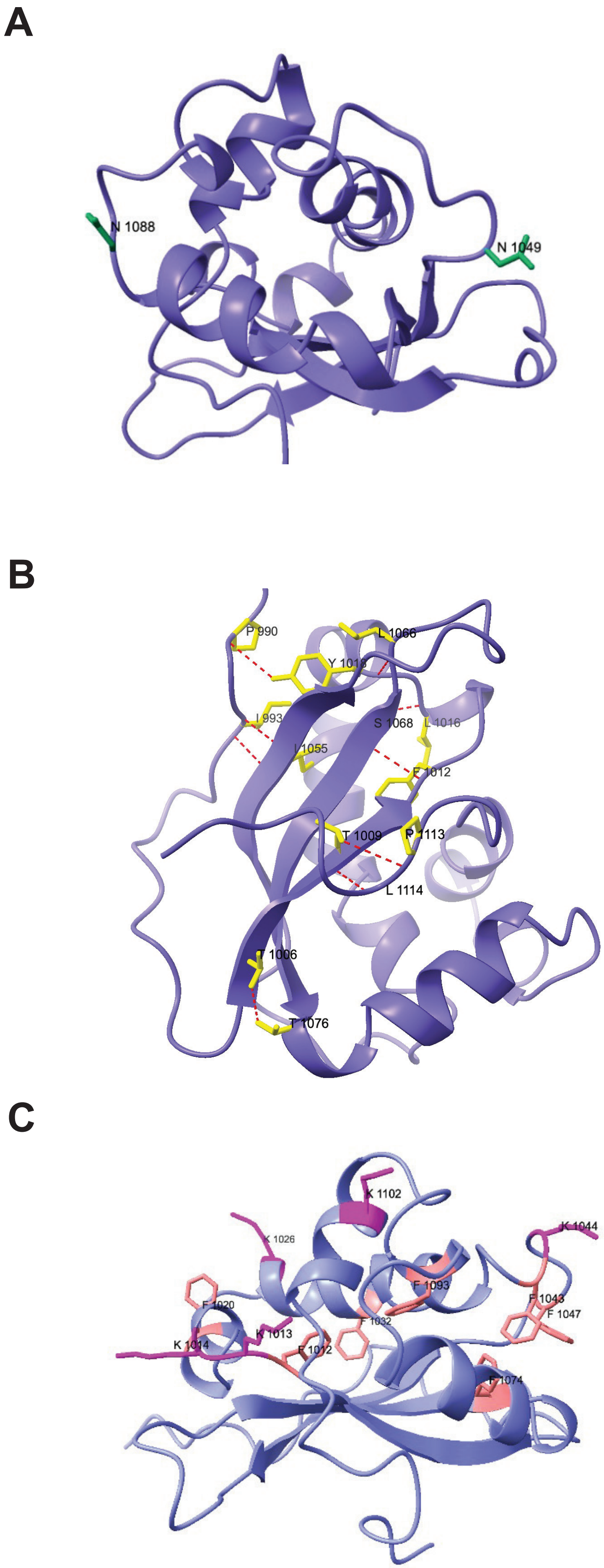
A
N 1088
N 1049
B
P 990
L 1066
I 993
S 1068
L 1016
I 1055
F 1012
T 1009
P 1113
L 1114
T 1006
T 1076
C
K 1102
K 1026
K 1044
F 1093
F 1043
F 1047
F 1020
F 1032
K 1013
K 1014
F 1012
F 1074

*Fig._S11*

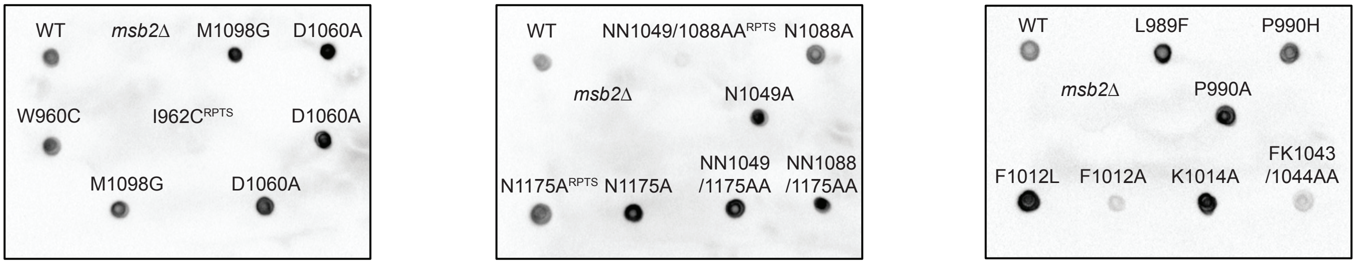


| | | | | | | | | |
|---|---|---|---|---|---|---|---|---|
| pRS316 | pHA-Msb2 | W961C | F1012A | S935A | N945A | L989A | P990R | P990W |
| T1009A | I1010A | G1011A | L1016A | N1017A | Y1018A | E1019A | F1020A | P1036A |
| N1040A | L1082A | V1108A | I1112A | P1113A | T1115A | G1126A | S1128A | S1232A |
| C1198A | W960A* | S930A* | P941A* | I993C* | V1072A* | L1114A* | G1116A* | S1129A* |
| | | | G1196A* | S1256Atr | G1196Atr | G1126Atr | | |

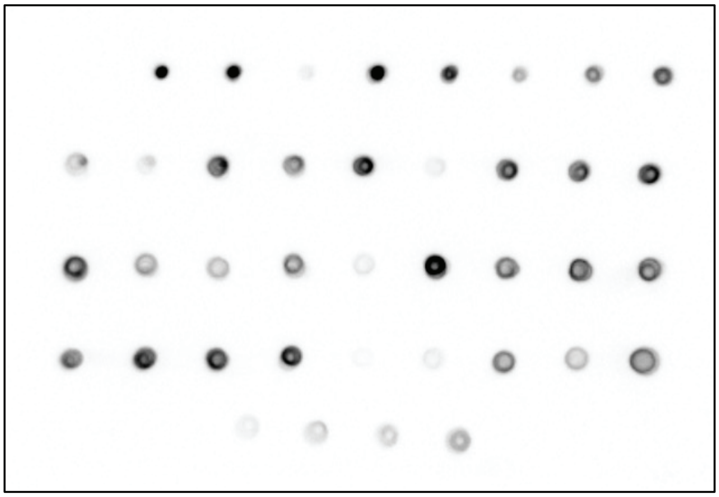

*Fig._S12*

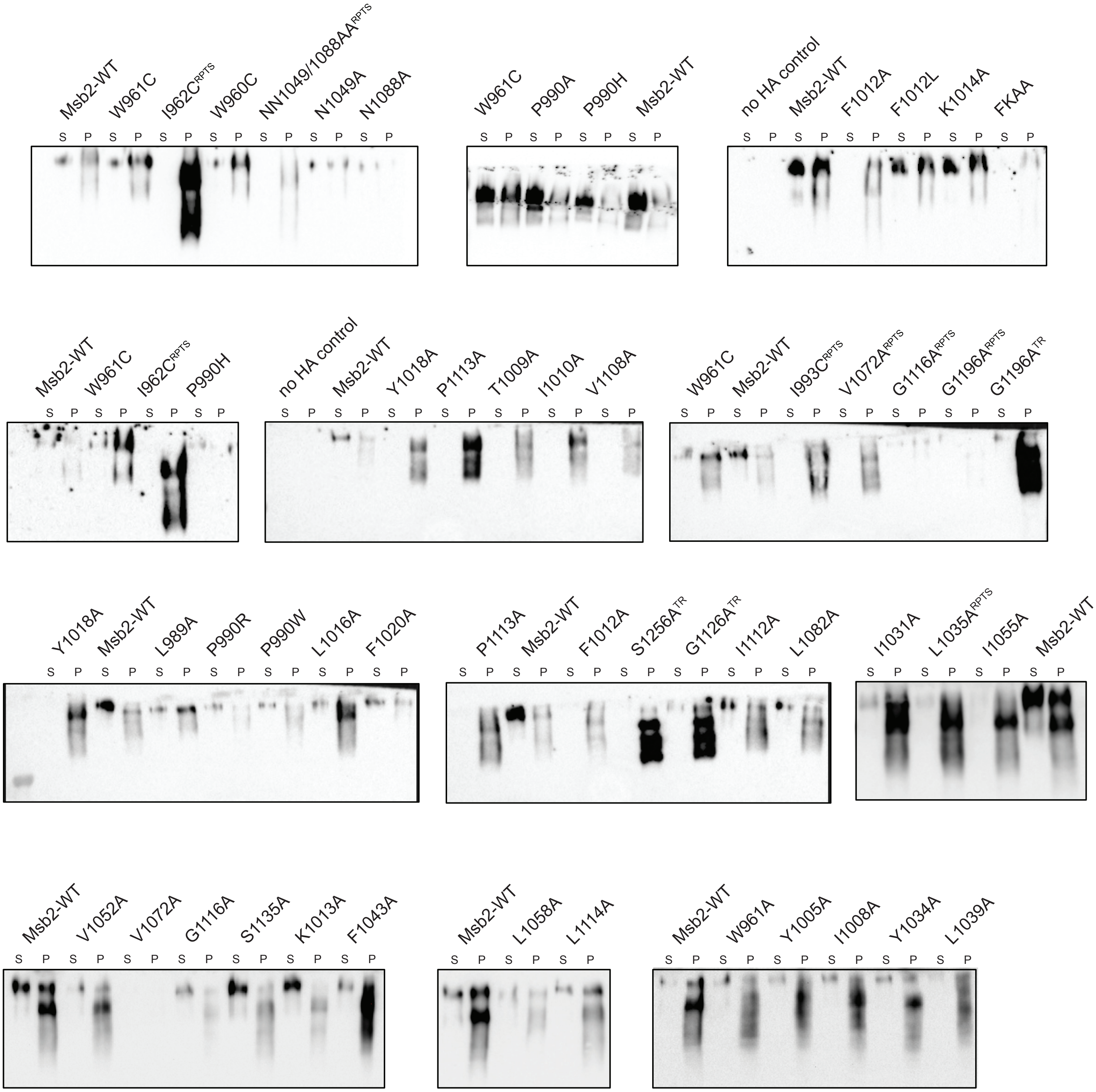

Msb2-WT
W961C
I962C$^{RPTS}$
W960C
NN1049/1088AA$^{RPTS}$
N1049A
N1088A
W961C
P990A
P990H
Msb2-WT
no HA control
Msb2-WT
F1012A
F1012L
K1014A
FKAA
Msb2-WT
W961C
I962C$^{RPTS}$
P990H
no HA control
Msb2-WT
Y1018A
P1113A
T1009A
I1010A
V1108A
W961C
Msb2-WT
I993C$^{RPTS}$
V1072A$^{RPTS}$
G1116A$^{RPTS}$
G1196A$^{RPTS}$
G1196A$^{TR}$
Y1018A
Msb2-WT
L989A
P990R
P990W
L1016A
F1020A
P1113A
Msb2-WT
F1012A
S1256A$^{TR}$
G1126A$^{TR}$
I1112A
L1082A
I1031A
L1035A$^{RPTS}$
I1055A
Msb2-WT
Msb2-WT
V1052A
V1072A
G1116A
S1135A
K1013A
F1043A
Msb2-WT
L1058A
L1114A
Msb2-WT
W961A
Y1005A
I1008A
Y1034A
L1039A
S P

Fig._S13

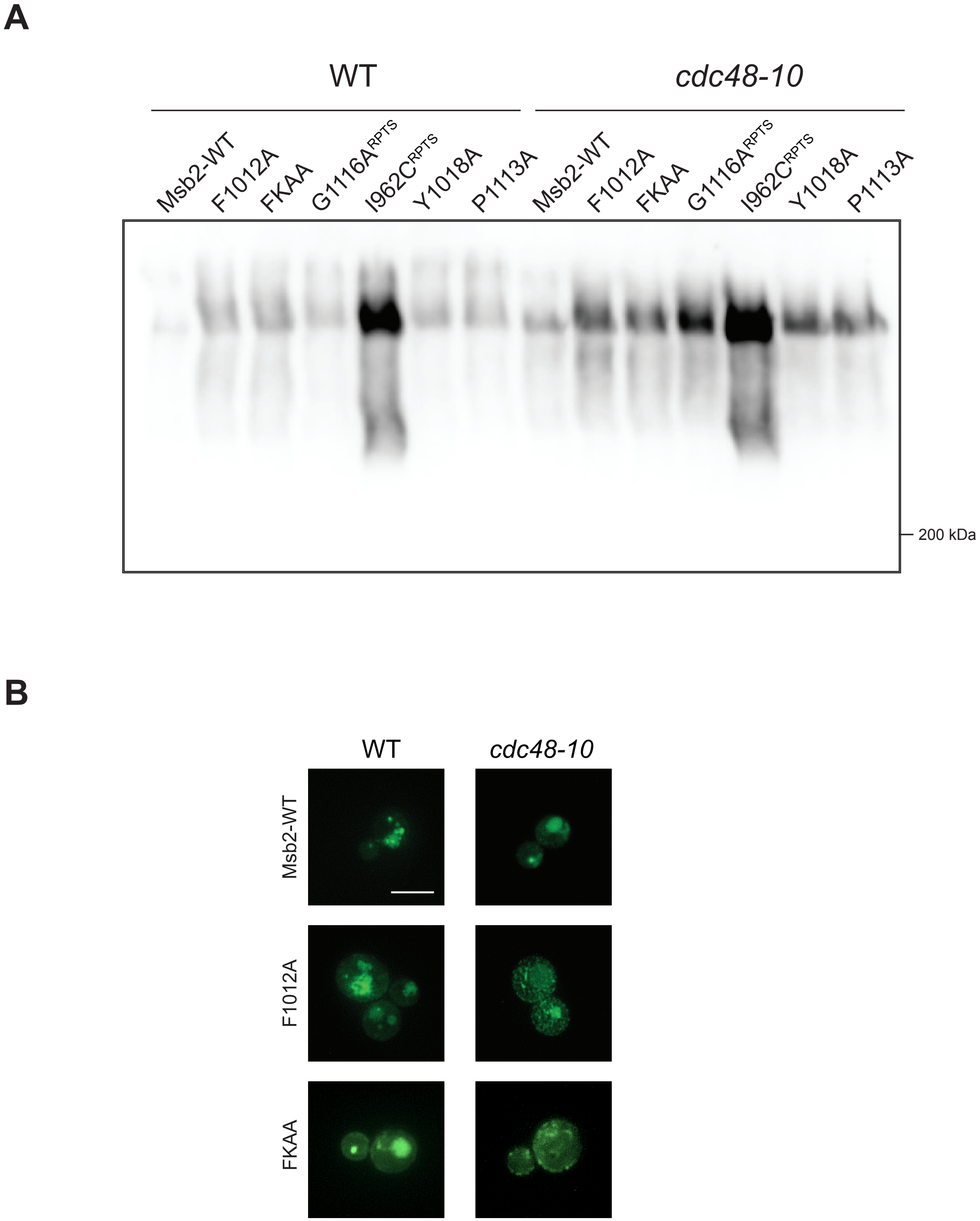
A
WT
cdc48-10
Msb2-WT
F1012A
FKAA
G1116A^RPTS
I962C^RPTS
Y1018A
P1113A
Msb2-WT
F1012A
FKAA
G1116A^RPTS
I962C^RPTS
Y1018A
P1113A
200 kDa
B
WT
cdc48-10
Msb2-WT
F1012A
FKAA

Fig._S14

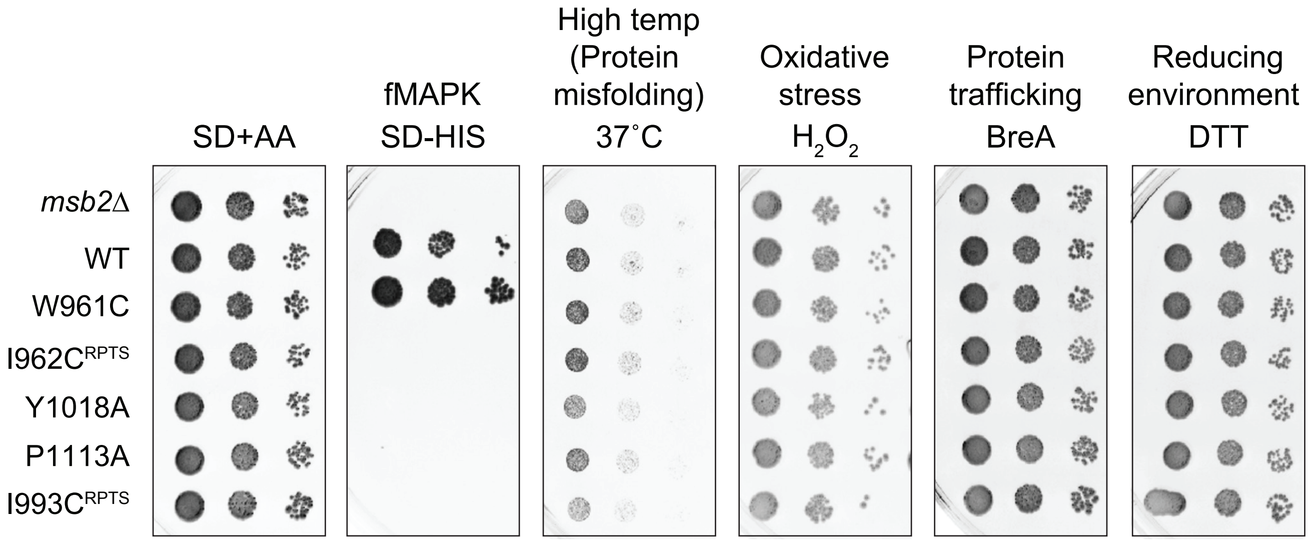
SD+AA
fMAPK
SD-HIS
High temp
(Protein
misfolding)
37˚C
Oxidative
stress
$H_2O_2$
Protein
trafficking
BreA
Reducing
environment
DTT
*msb2Δ*
WT
W961C
I962C RPTS
Y1018A
P1113A
I993C RPTS

Fig._S15

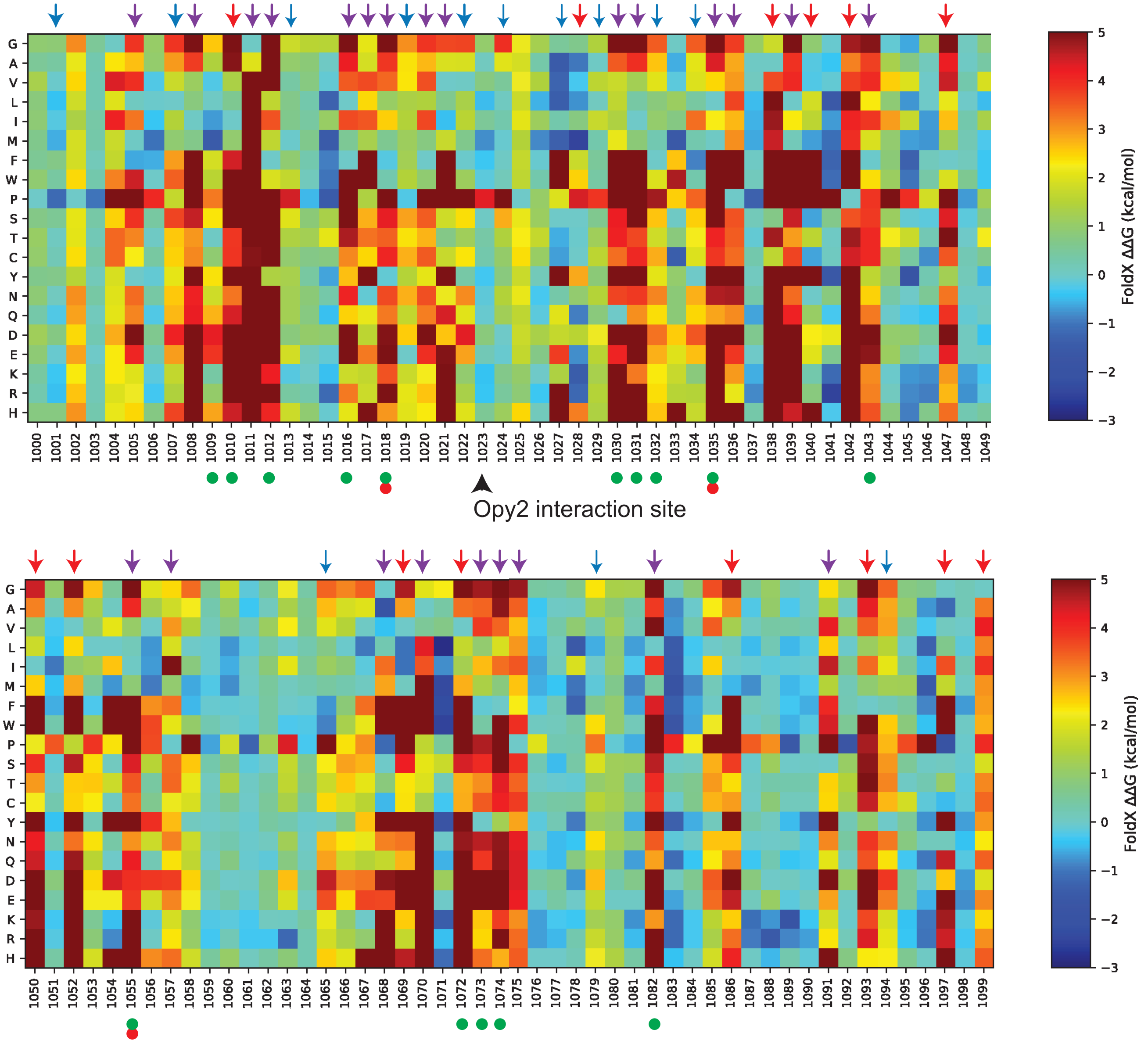


Key

- critical for fMAPK (includes LOF and partial LOF)
- critical for HOG
- conserved and critical for structure
- critical for structure
- conserved residue

Fig._S16

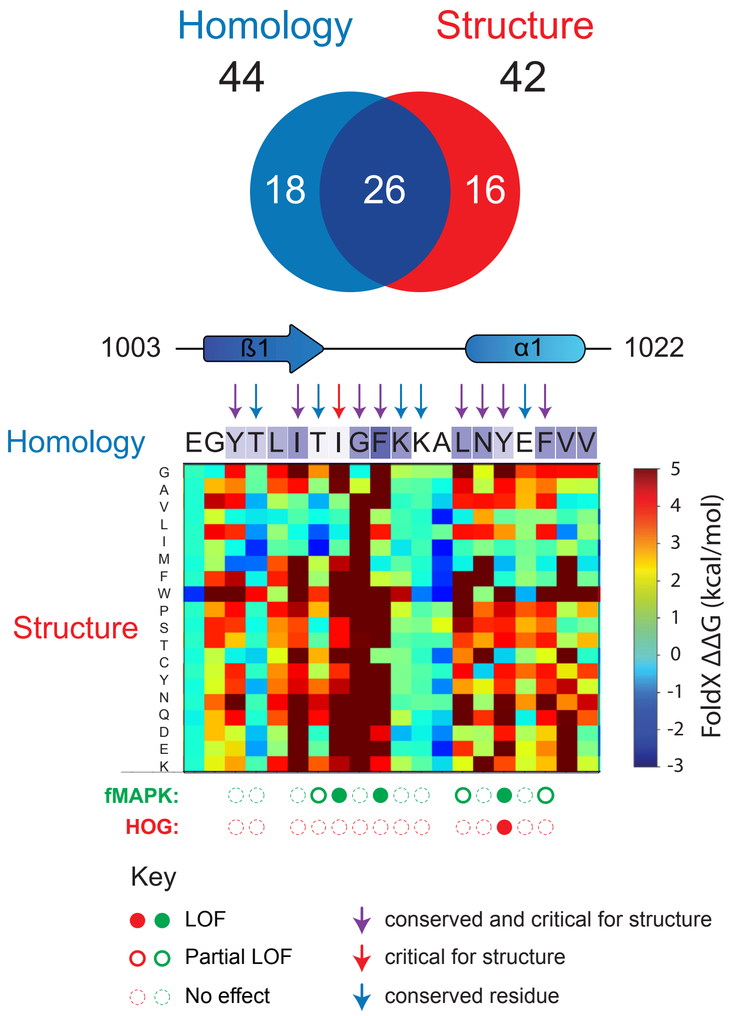

Homology
44
Structure
42
18
26
16
1003
ß1
α1
1022
Homology
EGYTLITIGFKKALNYEFVV
G A V L I M F W P S T C Y N Q D E K
Structure
5 4 3 2 1 0 -1 -2 -3
FoldX ΔΔG (kcal/mol)
fMAPK:
HOG:
Key
LOF
Partial LOF
No effect
conserved and critical for structure
critical for structure
conserved residue

A

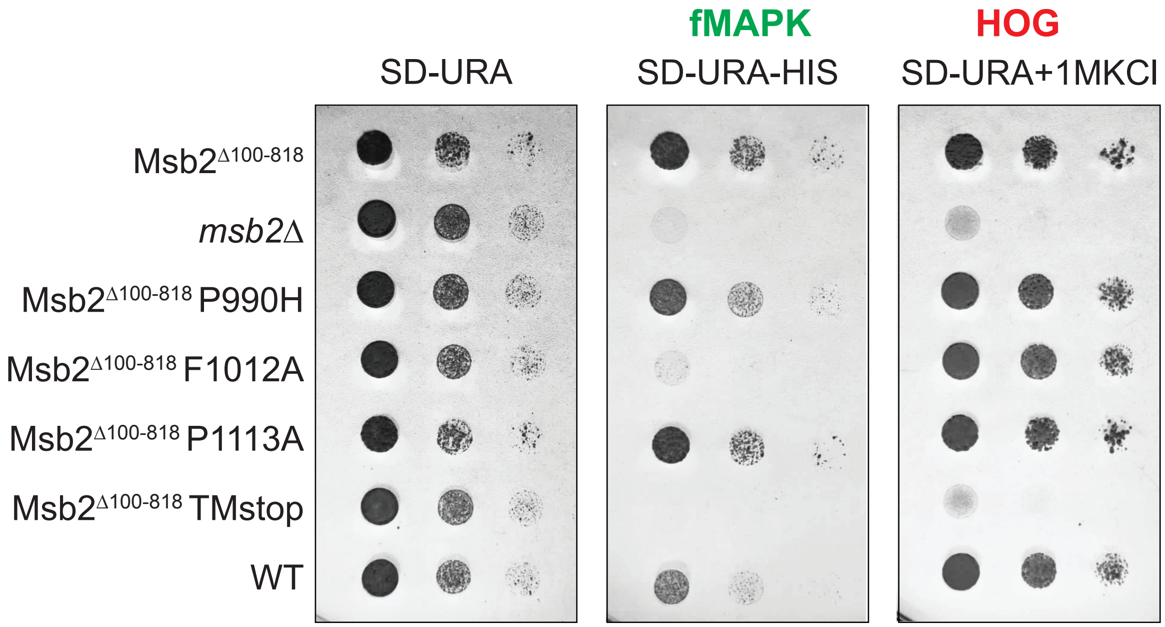
fMAPK
HOG
SD-URA
SD-URA-HIS
SD-URA+1MKCl
Msb2Δ100-818
msb2Δ
Msb2Δ100-818 P990H
Msb2Δ100-818 F1012A
Msb2Δ100-818 P1113A
Msb2Δ100-818 TMstop
WT

Msb2

B

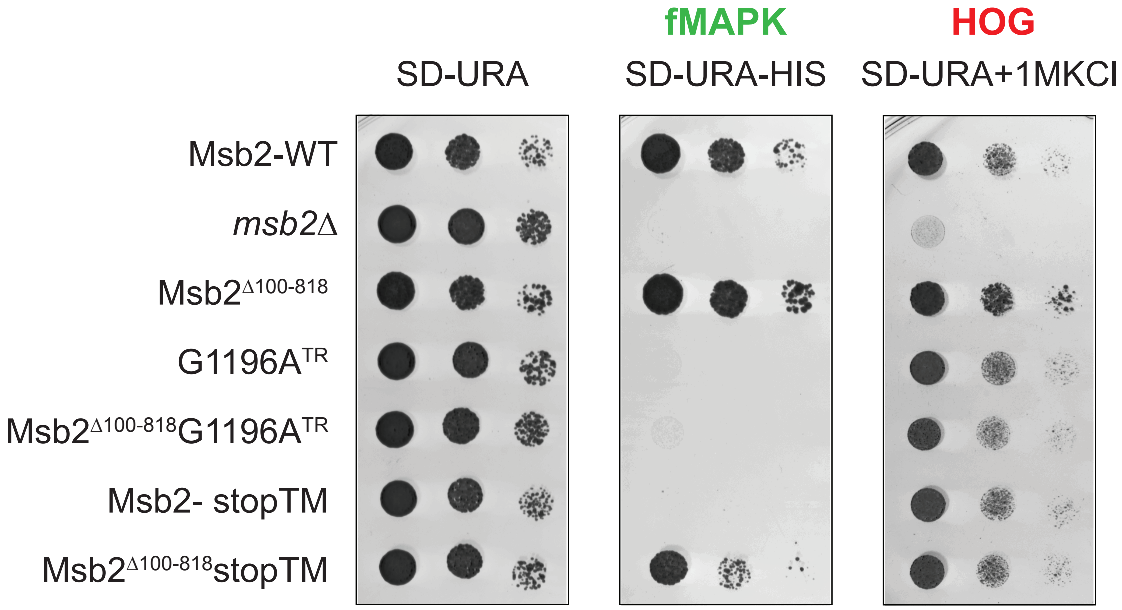
fMAPK
HOG
SD-URA
SD-URA-HIS
SD-URA+1MKCl
Msb2-WT
msb2Δ
Msb2Δ100-818
G1196ATR
Msb2Δ100-818G1196ATR
Msb2- stopTM
Msb2Δ100-818stopTM

*Fig._S18*

**A**

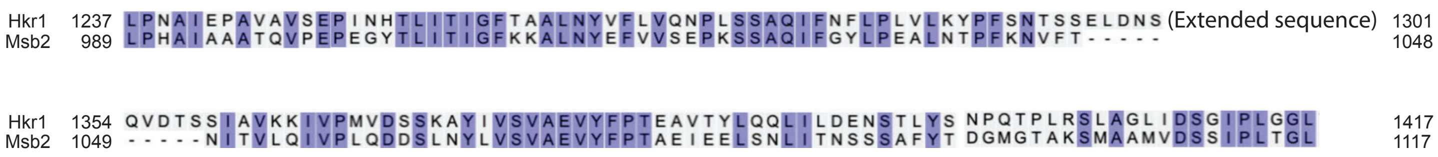
Hkr1 1237 LPNAIEPAVAVSEPINHTLITIGFTAALNYVFLVQNPLSSAQIFNFLPLVLKYPFSNTSSELDNS (Extended sequence) 1301
Msb2 989 LPHAIAAATQVPEPEGYTLITIGFKKALNYEFVVSEPKSSAQIFGYLPEALNTPFKNVFT----- 1048
Hkr1 1354 QVDTSSIAVKKIVPMVDSSKAYIVSVAEVYFPTEAVTYLQQLILDENSTLYS NPQTPLRSLAGLIDSGIPLGGL 1417
Msb2 1049 -----NITVLQIVPLQDDSLNYLVSVAEVYFPTAEIEELSNLITNSSSAFYT DGMGTAKSMAAMVDSSIPLTGL 1117

**B**

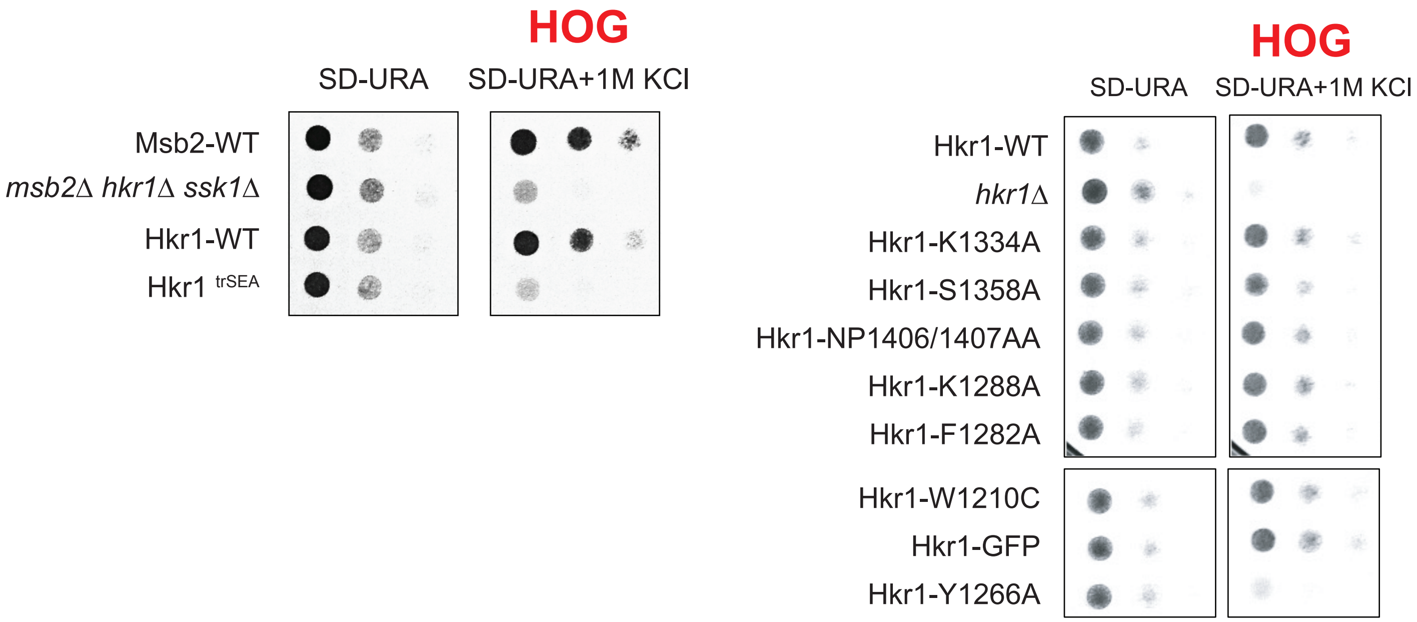
HOG
SD-URA
SD-URA+1M KCl
Msb2-WT
msb2Δ hkr1Δ ssk1Δ
Hkr1-WT
Hkr1 trSEA
HOG
SD-URA
SD-URA+1M KCl
Hkr1-WT
hkr1Δ
Hkr1-K1334A
Hkr1-S1358A
Hkr1-NP1406/1407AA
Hkr1-K1288A
Hkr1-F1282A
Hkr1-W1210C
Hkr1-GFP
Hkr1-Y1266A

Fig._S19

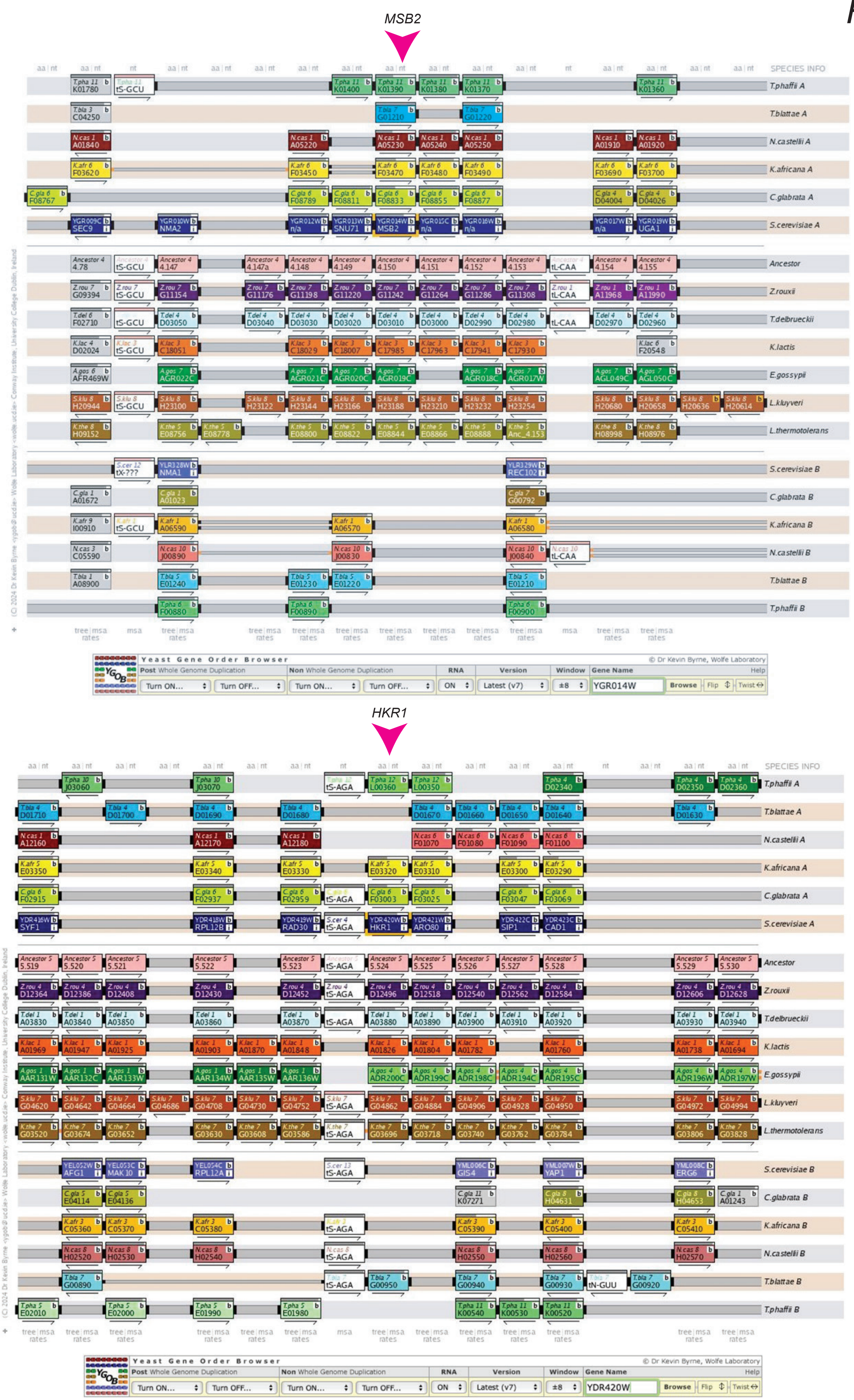
MSB2
SPECIES INFO
T.phaffii A
T.blattae A
N.castellii A
K.africana A
C.glabrata A
S.cerevisiae A
Ancestor
Z.rouxii
T.delbrueckii
K.lactis
E.gossypii
L.kluyveri
L.thermotolerans
S.cerevisiae B
C.glabrata B
K.africana B
N.castellii B
T.blattae B
T.phaffii B
(C) 2024 Dr Kevin Byrne <> Wolfe Laboratory <wolfe ucd.ie> Conway Institute, University College Dublin, Ireland
Yeast Gene Order Browser
© Dr Kevin Byrne, Wolfe Laboratory
Post Whole Genome Duplication
Non Whole Genome Duplication
RNA
Version
Window
Gene Name
Help
Turn ON...
Turn OFF...
ON
Latest (v7)
±8
YGR014W
Browse
Flip
Twist
HKR1
SPECIES INFO
T.phaffii A
T.blattae A
N.castellii A
K.africana A
C.glabrata A
S.cerevisiae A
Ancestor
Z.rouxii
T.delbrueckii
K.lactis
E.gossypii
L.kluyveri
L.thermotolerans
S.cerevisiae B
C.glabrata B
K.africana B
N.castellii B
T.blattae B
T.phaffii B
Yeast Gene Order Browser
© Dr Kevin Byrne, Wolfe Laboratory
YDR420W
Browse

Fig._S20

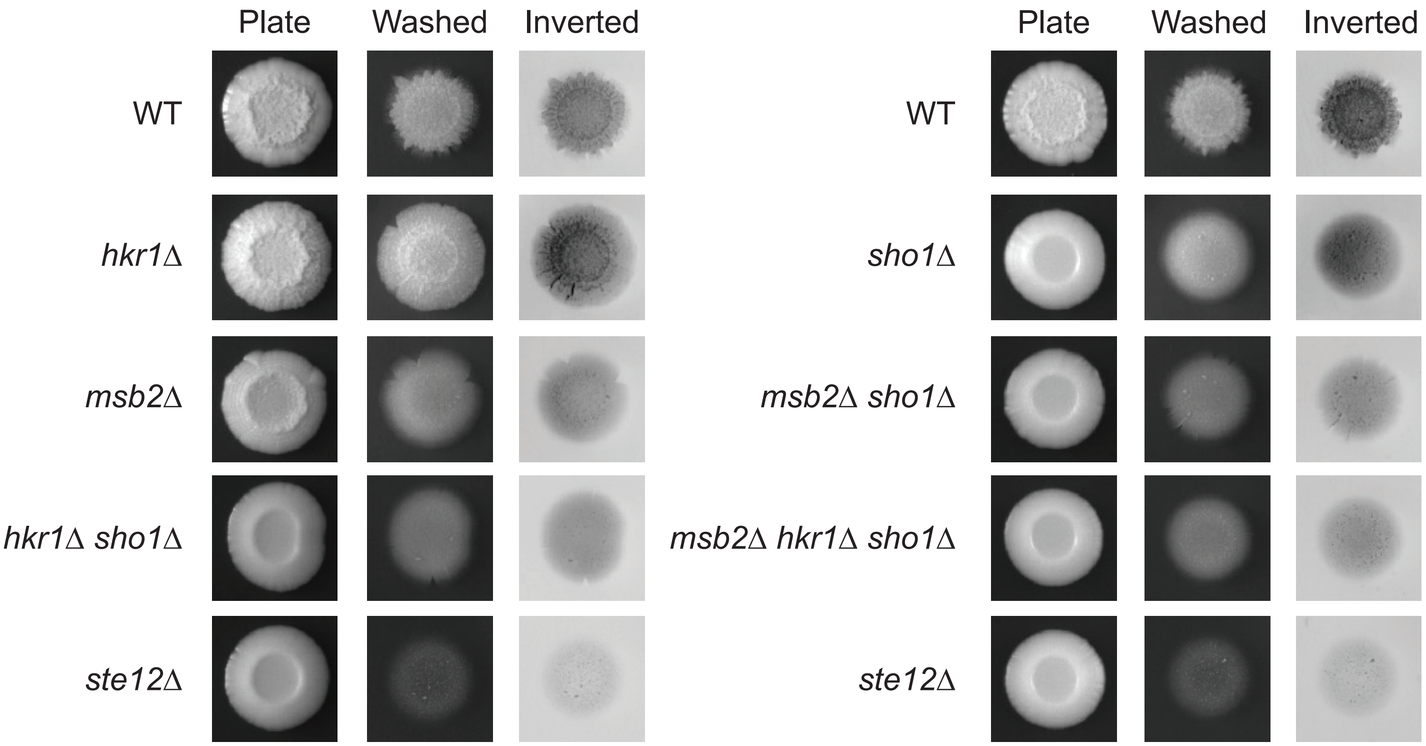
Plate
Washed
Inverted
WT
hkr1Δ
msb2Δ
hkr1Δ sho1Δ
ste12Δ
Plate
Washed
Inverted
WT
sho1Δ
msb2Δ sho1Δ
msb2Δ hkr1Δ sho1Δ
ste12Δ

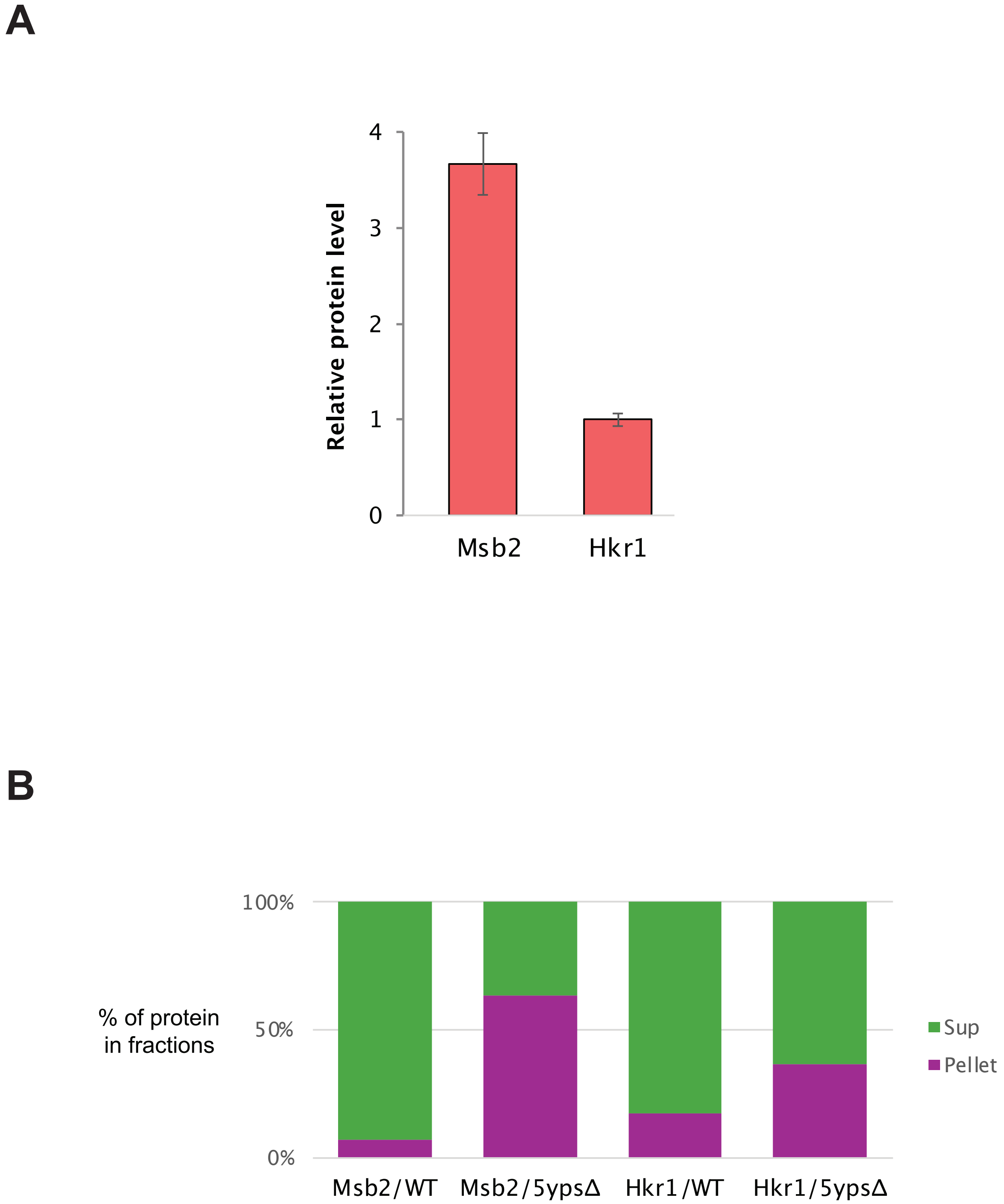
A
Relative protein level
4
3
2
1
0
Msb2
Hkr1
B
% of protein in fractions
100%
50%
0%
Msb2/WT
Msb2/5ypsΔ
Hkr1/WT
Hkr1/5ypsΔ
Sup
Pellet

Fig._S22

**A**

Msb2
Opy2
Sho1

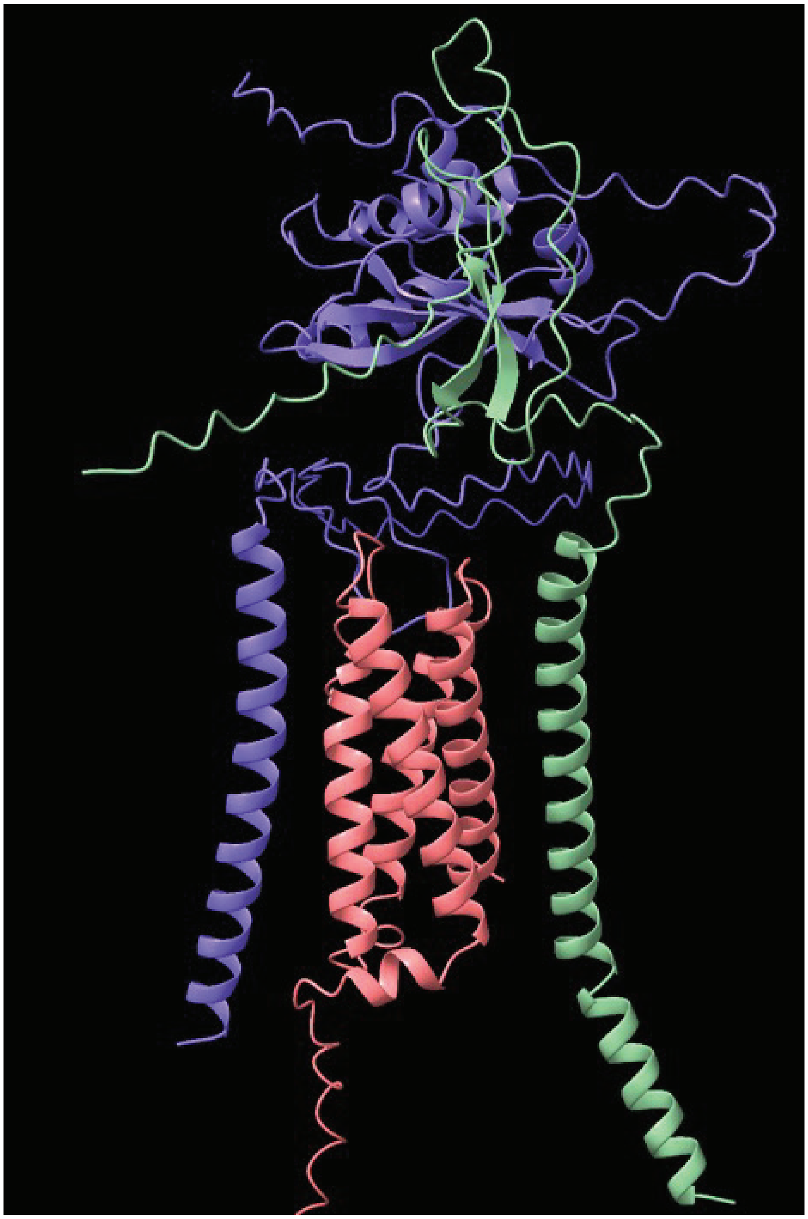

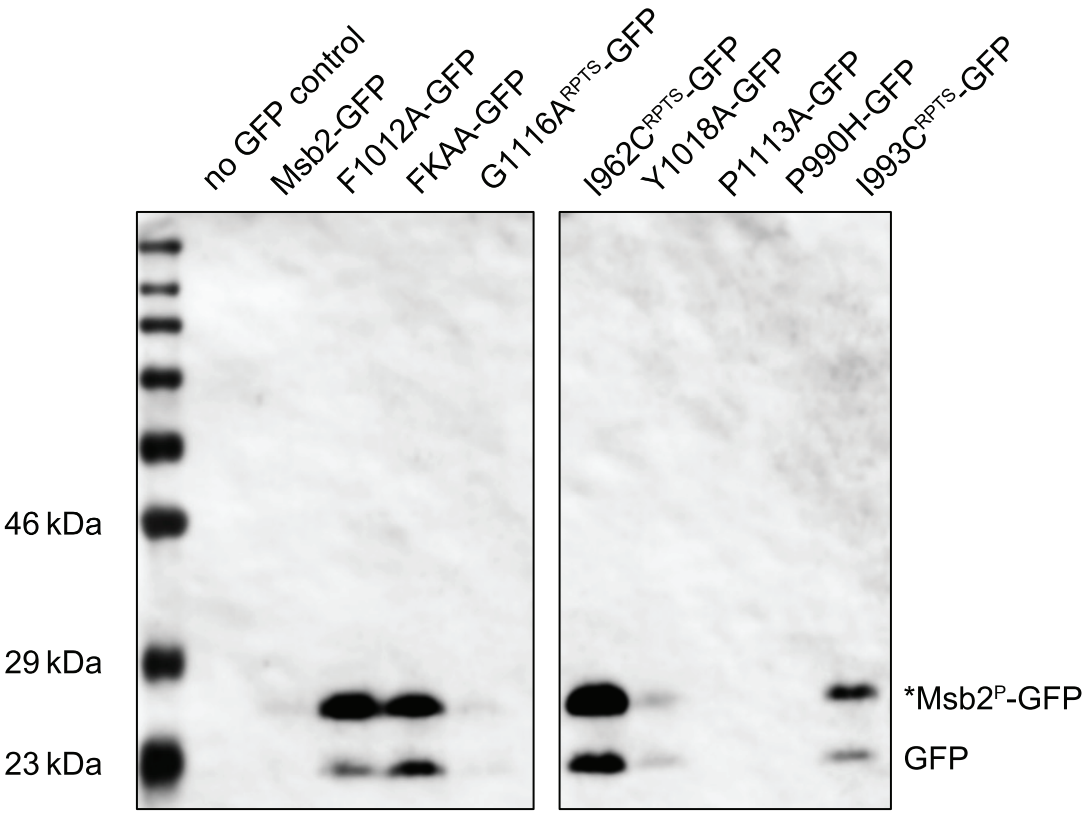
no GFP control
Msb2-GFP
F1012A-GFP
FKAA-GFP
G1116A^RPTS-GFP
I962C^RPTS-GFP
Y1018A-GFP
P1113A-GFP
P990H-GFP
I993C^RPTS-GFP
46 kDa
29 kDa
23 kDa
*Msb2^P-GFP
GFP
